# A synthetic tear protein resolves dry eye through promoting corneal nerve regeneration

**DOI:** 10.1101/2022.05.07.491018

**Authors:** Yael Efraim, Feeling Yu Ting Chen, Ka Neng Cheong, Eliza A. Gaylord, Nancy A. McNamara, Sarah M. Knox

## Abstract

Corneal architecture is essential for vision and is greatly perturbed by the absence of tears due to the highly prevalent disorder dry eye. With no regenerative therapies available, pathological alterations of the ocular surface in response to dryness, including persistent epithelial defects and poor wound healing, result in lifelong morbidity. Here, using a mouse model of aqueous-deficient dry eye, we reveal that topical application of the synthetic tear protein lacripep reverses the pathological outcomes of dry eye through restoring the extensive network of corneal nerves that are essential for tear secretion, barrier function, epithelial homeostasis and wound healing. Intriguingly, the restorative effects of lacripep occur despite extensive immune cell infiltration, suggesting tissue reinnervation and regeneration can be achieved under chronic inflammatory conditions. In summary, our data highlight lacripep as a first-in-class regenerative therapy for returning the cornea to a near homeostatic state in individuals who suffer from dry eye.

**Teaser:** Topical application of a synthetic tear protein repairs dry eye-mediated corneal damage through driving functional sensory reinnervation.

## Introduction

Tear deficiency due to lacrimal gland dysfunction (aqueous-deficient dry eye) is among the most common and debilitating outcomes of systemic autoimmune diseases including Sjogren’s, rheumatoid arthritis, scleroderma and systemic lupus erythematosus (*1*). A healthy tear film provides an aqueous coating necessary for optimal vision and tissue function while also shielding the ocular surface from environmental, inflammatory, and microbial insult. Due to the essential requirement of tears in maintaining ocular health, corruption of tissue integrity and loss of homeostasis in response to prolonged dryness induce a vast array of pathological outcomes (*2*) Yet, despite the extensive ramifications of dry eye on ocular health and its significant impact on vision, quality of life, and the psychological/physical consequences of chronic pain (*3*), there are currently only three clinically-approved therapies for the treatment of dry eye disease that specifically target T-cell mediated inflammatory pathways believed to be the primary driver of dry eye pathogenesis. However, these anti-inflammatory treatments are not regenerative and only promote modest improvements in the signs and symptoms of dry eye (*2*), which results in life-long corneal dysfunction and reduced quality of life.

Due to the fundamental requirement for tears in corneal maintenance and the clear role of desiccating stress as a principal driver of dry eye pathogenesis, there has been a recent focus on the application of tear-promoting factors to relieve dry eye. Lacritin, an endogenous glycoprotein identified in tears that is deficient in dry eye patients (*4*), has been found to possess pro-secretory properties in healthy and diseased animal models (*4, 5*), and to promote corneal epithelial cell proliferation in vitro (6). These findings led to the development of lacripep, a stable synthetic peptide consisting of lacritin’s active C-terminal fragment, that has also been shown to stabilize the human tear film (*7*). However, its impact on cornea regeneration, including tissue architecture and epithelial cell identity, integrity and homeostasis, as well as physiological tear secretion, during dry eye disease progression has not been investigated. Furthermore, whether application of lacripep to the desiccated cornea can restore the significantly depleted functional nerve supply that is essential for basal tear secretion, ocular surface integrity, and corneal wound healing/tissue regeneration, remains unknown.

Here, using the well-characterized, autoimmune regulator (*Aire*)-deficient mouse model of spontaneous, autoimmune-mediated dry eye, we show that topical administration of lacripep resolves dry eye disease through reactivating basal tear secretion, restoring progenitor cell identity, and rescuing epithelial barrier function. Lacripep achieves these outcomes through re-establishing functional sensory innervation of the corneal epithelium and can do so despite the presence of chronic ocular inflammation. Thus, we have identified the first regenerative ocular therapeutic for dry eye disease that restores the secretory and epithelial integrity of the desiccated cornea through promoting sensory reinnervation.

## Results

### Basal tear secretion, epithelial integrity and basal progenitor cell identity are restored with lacripep treatment during dry eye disease progression

The process of reflex (basal) tear secretion that occurs in response to changes at the ocular surface is essential to corneal function, homeostasis and wound healing and is profoundly reduced in human patients with dry eye disease. Although lacritin has been shown to promote basal tearing in a healthy rabbit model (*5*), whether lacripep is also capable of increasing basal tear production and can do so during dry eye disease progression is unknown. To test this, we evaluated basal tearing by utilizing the *Aire* mouse model of aqueous deficient dry eye (*8*). *Aire* knockout (KO) female mice develop classic signs of dry eye disease over a 2-week time period. This begins with small lymphocytic infiltrates in the lacrimal glands and mildly reduced tear secretion, corneal innervation, and barrier function at 5 weeks (wks), consistent with mild aqueous-deficient dry eye, that progresses to extensive CD4+ T cell-mediated exocrinopathy, severe tear reduction, and corneal pathologies at the epithelial, stromal and nerve-associated levels by 7 wks (*8–10*).

To determine the impact of lacripep on physiological (basal) tear secretion, lacripep (4 µM) or phosphate buffered saline (PBS, control) was topically applied to *Aire* KO corneas 3 times (3x) per day for 14 consecutive days (from 5 to 7 wk of age) and tear production was measured at day 0, 7 and 15 (Fig.1A). Basal tear levels for the untreated *Aire* KO mice were significantly reduced at day 0 (77% of WT levels) and continued to decrease over time (67% at day 7 and 37% at day 15) (Fig.1B). Tearing was also significantly decreased for the PBS-treated mice, with tear volumes being 50% of the WT controls at day 15 (Fig.1B). Although lacripep-treated *Aire* KO mice showed similar levels of tear production as untreated and PBS-treated mice at day 7, at day 15 tear volumes remained at day 0 levels (83% of WT) (Fig.1B), demonstrating lacripep retains pro-secretory functions during dry eye disease progression.

**Fig. 1.**
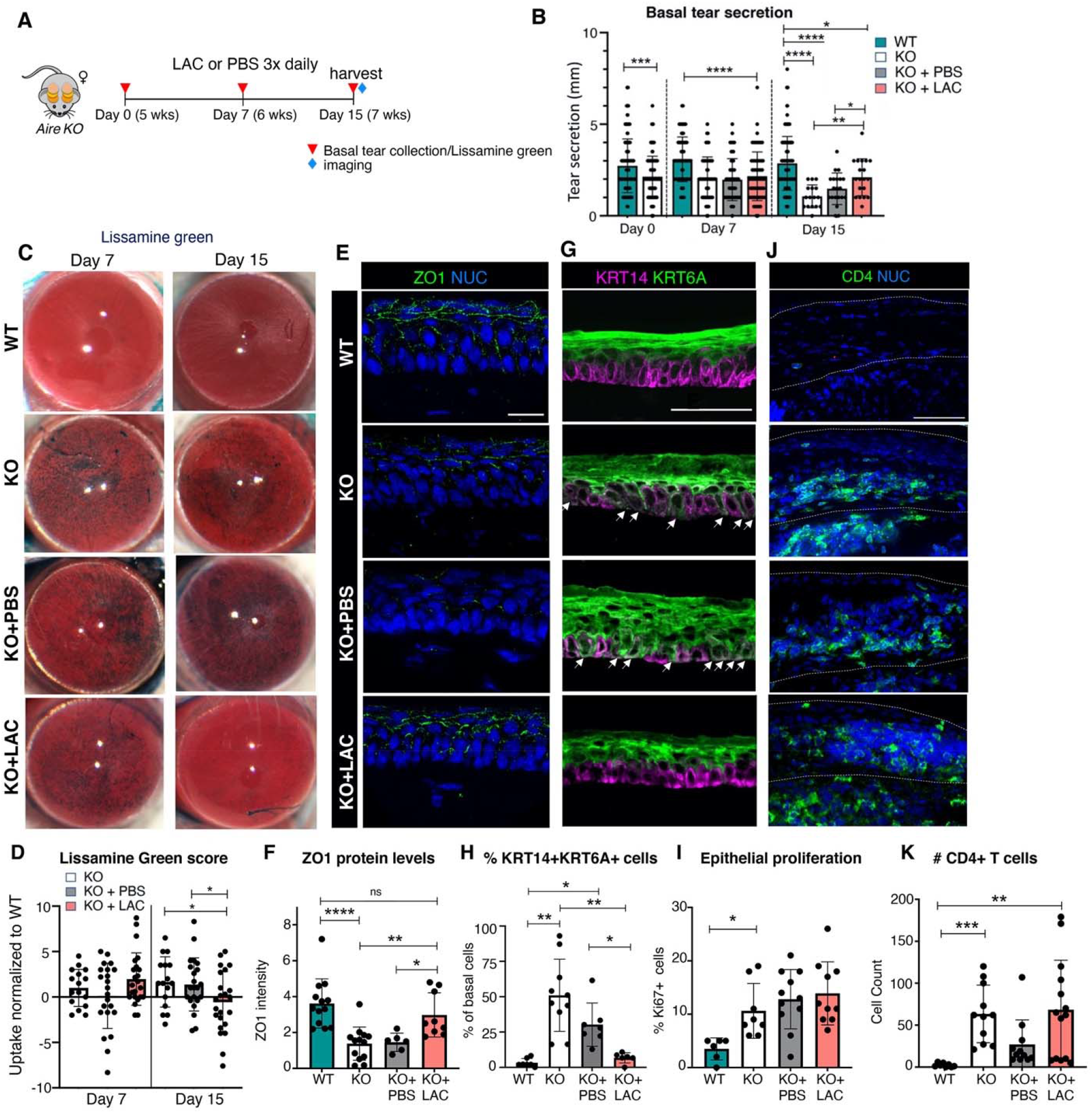
Basal tear secretion, epithelial integrity and basal progenitor cell identity are restored with lacripep treatment during dry eye disease progression. **A**, Schematic of treatment regimen showing data collection time points for tissue analysis (top). **B**, Levels of physiological (basal) tear secretion at day 0, 7 and 15. **C-D**, Lissamine green uptake in untreated/treated *Aire KO* corneas compared to age matched WT as assessed by scoring intensity of stain at day 7 and 15. The day 7 score was normalized to day 0 (before treatment) and the day 15 score was normalized to day 7. Data points above 1 indicate increased lissamine green uptake while points below 1 indicate reduced uptake. NUC = nuclei. **E-H**, Immunofluorescent analysis and quantification of the tight junction protein ZO1 (**E**,**F**), and basal progenitor cell marker KRT14 and differentiation marker KRT6A (**G**,**H**) at day 15. Arrows in **G** highlight basal cells co-expressing KRT6A and KRT14. The graph in **H** shows the percentage of basal KRT14+ basal cells co-expressing KRT6A. **I**, Quantification of proliferating epithelial cells at day 15. **J-K**, Immunofluorescent analysis and quantification of CD4+ T cells in WT and untreated/treated *Aire* KO corneas at day 15. Scale bars = 50 μm. *p < 0.05; **p < 0.01; ***p < 0.001; ****p < 0.0001; a one-way analysis of variance was applied to graphs B, D, F, H and K with the following post-hoc tests: a Tukey’s multiple comparisons test was used in B, D and F, and a Dunett’s T3 multiple comparisons test was used in H and K. Graph I was analyzed using a one-way analysis of variance with correction for multiple comparisons using a False Discovery rate of 0.05 and a two-stage step-up method of Benjamini, Krieger and Yekutieli. Each dot in the bar graph represents a biological replicate. All data are expressed as mean + s.d. n > 4 mice per group.

As dry eye results in disruption of ocular surface integrity resulting in cornea barrier dysfunction and, consequently, fluid loss and pathogen invasion (*11*), we next determined whether lacripep maintains corneal epithelial architecture by analyzing lissamine green uptake at day 7 and 15. At day 7 all *Aire* KO treatment groups exhibited increased lissamine green uptake compared to WT controls (Fig.1C,D). However, at day 15, epithelial barrier function in lacripep-but not PBS-treated or untreated corneas was significantly improved, with uptake levels resembling those of WT corneas (Fig.1C,D). Restoration of barrier function was further confirmed by analyzing the superficial epithelial layers for the tight junction protein ZO1 which is essential to the structural integrity of the corneal epithelium (*12*). Levels of ZO1 were greatly reduced in the untreated and PBS treated *Aire* KO cornea (Fig.1E,F). In contrast, ZO1 expression in lacripep-treated corneas mimicked that of WT controls (Fig.1E,F), thus, demonstrating lacripep is sufficient to rescue functional integrity of the corneal epithelium.

Although it has been well established that basal progenitor cells (also referred to as transit amplifying cells (*13*)) undergo mitosis and migration (*14*) to give rise to the barrier forming suprabasal epithelial and stratified squamous epithelial cells, the impact of dry eye disease on basal cell identity and differentiation remains largely unknown. Thus, we questioned whether basal cell identity was altered in the desiccated cornea through defining the cellular location of KRT6A, a marker of early epithelial cell differentiation in skin (*15*) and cultured cornea epithelial cells (*16*). Accordingly, we confirmed that KRT6A marks differentiated corneal epithelial cell types in vivo, as shown by its exclusive expression in KRT14-deficient suprabasal and superficial epithelial cells of healthy cornea at 7 wks of age (Fig.1G). In contrast, KRT6A appeared in a large cohort of KRT14+ basal cells in the untreated *Aire* KO cornea (Fig.1G, arrows), suggesting progenitors were undergoing aberrant differentiation. Treatment of *Aire* KO cornea with PBS for 2 wks did not reverse this outcome, with the number of KRT6A+KRT14+ cells being similar to untreated controls (Fig.1G, H). Strikingly, however, cell identity in lacripep-treated corneas returned to that of WT tissue, with KRT6A remaining exclusively expressed by suprabasal/superficial cells (Fig.1G, H). To determine if changes in cell identity alters progenitor cell division, we analyzed the percentage of proliferating K14+ basal cells in treated and untreated corneas. In the untreated *Aire* KO corneas, K14+ cell division was significantly increased, as denoted by Ki67, compared to WT controls (Fig.1I). A similar outcome was found for both PBS-and lacripep-treated *Aire* KO corneas, indicating cell identity but not cell division is impacted by lacripep.

We next tested whether lacripep restored basal cell identity and barrier function by reducing immune cell infiltration. Prolonged desiccating stress from reduced tear production elicits an immune response in which activated CD4+ T cells home to the corneal epithelium and stroma (primarily accumulating at the limbal region) to modulate epithelial differentiation (*17*) and direct the development of autoimmune mediated dry eye (*18*). Intriguingly, at day 15 we found all *Aire* KO corneas (Fig.1I,J), as well as their corresponding lacrimal glands (Fig.S1A, B), regardless of treatment, to be extensively infiltrated by CD4+ T cells. Furthermore, the extent of immune mediated damage to *Aire* KO gland function under each condition was comparable as there was no difference between maximal tear secretion in response to systemic injection of the muscarinic agonist pilocarpine (*9,19*). This outcome was further supported by RNAseq analysis at day 7 that showed (i) the expression of T cell markers such as *Cd3, Cd7*, and *Cd8*, (ii) T cell specific response genes, such as T cell-specific guanine nucleotide triphosphate-binding protein 1 and 2 (*Tgtp1* and *Tgtp2*), and (iii) macrophage markers, such as *Cd64*, remained significantly elevated in the untreated and PBS/lacripep treated *Aire* KO corneas compared to the WT controls (Fig.S1D; Data S1 and Table S1). In addition, key inflammatory mediators that are typically induced in desiccated corneas, including interleukin-1 beta (IL1b), and interferon response genes (*Irgm1, Ifi203, Ifit1, Ccl22, Ccl5*, and *Stat1*) were expressed at similar levels in all diseased corneas (Fig.S1D and Data S1 and Table S1).

Thus, together these data indicate that, despite the ongoing, large-scale inflammatory response that occurs during dry eye disease progression, lacripep is capable of improving corneal barrier function and rescuing cell identity in the desiccated cornea without dampening inflammation.

### Sensory innervation of the desiccated cornea is restored in response to lacripep treatment

As basal tear production is dependent on sensory innervation of the cornea, and dry eye leads to diminished corneal innervation in humans (*20, 21*) and mice (*8, 22*), we next determined whether the maintenance of reflex tearing in response to lacripep was due to restoration of the nerve supply. Sensory nerves derived from the ophthalmic lobe of the trigeminal ganglion primarily enter the corneal stroma at the limbal region and extend along the basement membrane, with axon fibers branching upward through the multi-layered epithelium to establish nerve-epithelial interactions. Here, they serve an essential role in sensing changes at the ocular surface (e.g., dryness) and maintaining corneal epithelial homeostasis, in part, through activation of tear production from the lacrimal gland via the lacrimal reflex. Thus, the degree of basal tearing reflects the function and quality of sensory nerves within the corneal epithelium. Consistent with this requirement, basal tear production and corneal nerve fiber density are significantly reduced in the *Aire KO* mouse model (beginning at 5 wks (*9*)) similar to human patients suffering from dry eye disease e.g., due to Sjögren’s (*21, 23*), an outcome that strongly correlates with loss of active lacritin in human tears (*24*).

To test for changes in innervation, corneas isolated from the four treatment groups at day 15 were immunostained for growth-associated protein 43 (GAP43), a marker of remodeling axons (*25*), along with two common sensory neuropeptides, substance P (SP) and calcitonin-gene related protein (CGRP). SP and CGRP are differentially expressed to carry out discrete functions: SP is released in response to trigeminal activation to modulate tear secretion and goblet cell function (*26*) while CGRP is involved in multiple homeostatic processes, including corneal epithelium regeneration and regulation of vasculature (*27, 28*). As shown in the whole mount images in Figure 2A, WT corneas showed extensive innervation by GAP43+ nerve fibers expressing SP and/or CGRP (Fig.2A,B), consistent with the constant remodeling and functionality of corneal nerves (*24*). In contrast to WT, nerve density was dramatically reduced in the untreated and PBS-treated *Aire* KO corneas, indicating lubrication alone is not sufficient to maintain a functional nerve supply (Fig.2A). Strikingly, however, lacripep-treated corneas showed extensive innervation throughout the tissue, with the density of highly branched GAP43+ nerve fibers expressing SP and CGRP being nearly equivalent to that of the WT controls (Fig.2A,B). Thus, lacripep treatment successfully regenerates the sensory nerve supply to the inflamed cornea during disease progression.

**Fig. 2.**
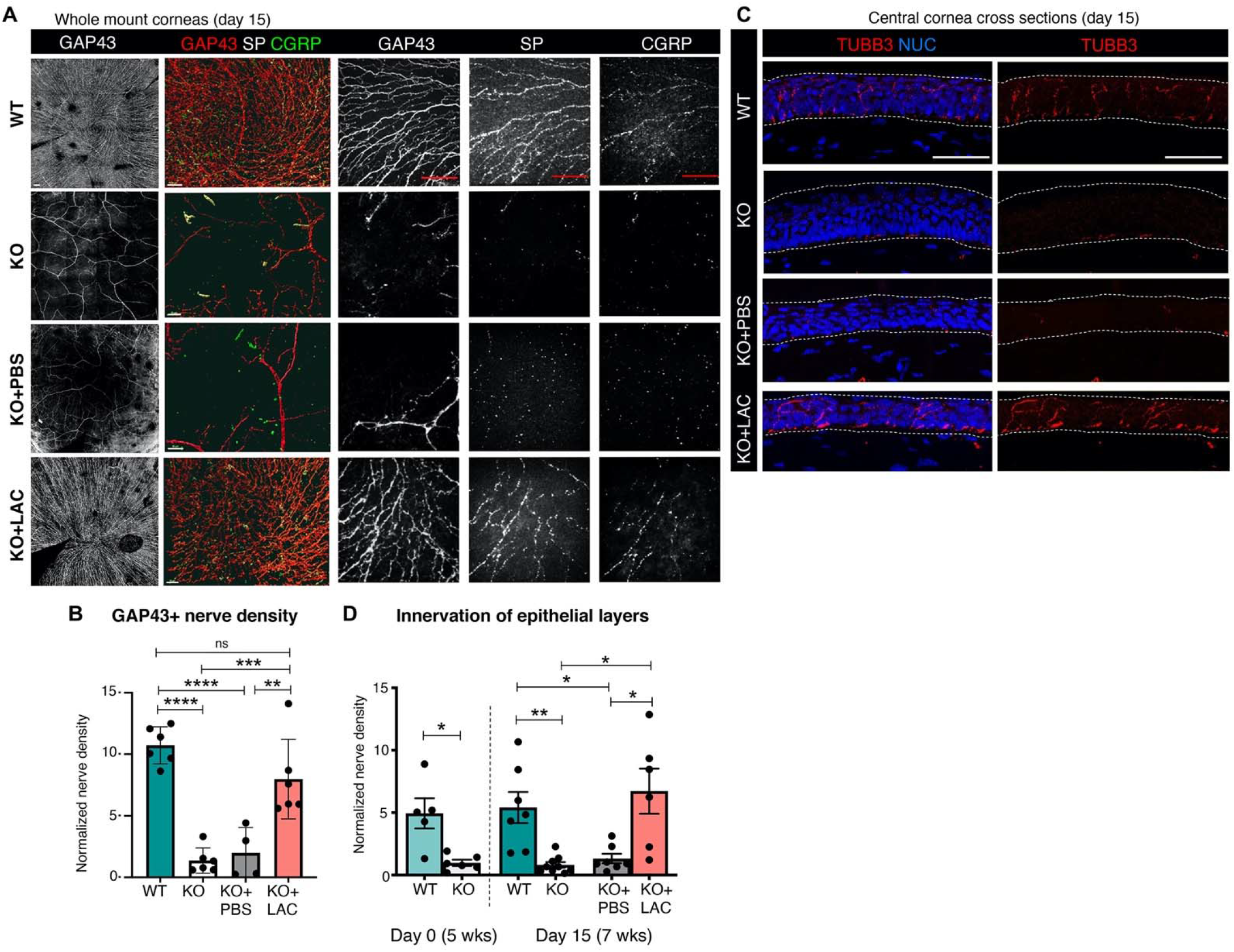
Sensory innervation of the desiccated cornea is restored in response to lacripep treatment. **A**, 3D projections of nerve fibers across the peripheral and central regions of WT and untreated, PBS-or lacripep-treated *Aire KO* cornea (whole mount). Nerves were immunolabeled for the remodeling neuronal marker GAP43, and sensory neurotransmitters substance P (SP), and calcitonin-gene-related peptide (CGRP). **B**, Quantification of GAP43+ nerves in WT, and untreated/treated *Aire* KO corneas. **C**, Central corneal cross-sections (10-12 µm) were immunolabeled for the pan-neuronal marker TUBB3 and nuclei (NUC). Scale bar = 70 μm. **D**, Quantification of TUBB3+ nerves in the central corneal epithelium of WT, untreated and treated *Aire* KO. Scale bar = 50 μm. *p < 0.05; **p < 0.01; ***p < 0.001; ****p < 0.0001; ns = not significant. Data in graph B was subjected to a one-way analysis of variance with a post-hoc Tukey’s test. In graph D, a Student’s T test was applied to WT vs KO at day 0 and a one-way analysis of variance with a post-hoc Tukey’s test was applied to day 15. Each dot in the bar graph represents a biological replicate. All data are expressed as mean + s.d. n >4 mice per group.

### Lacripep re-establishes functional corneal nerve-epithelial interactions during dry eye disease progression

Given nerve-epithelial interactions are essential for mediating physiological tear secretion, we next assessed innervation to corneal epithelial cell layers at 2 wks of treatment (day 15, 7 wks of age) through immunofluorescent analysis. As shown in Figure 2C, the corneal epithelium of untreated and PBS-treated mice were highly deficient in axons compared to the extensively innervated WT controls (Fig.2C,D). In contrast, innervation of lacripep-treated epithelial cell layers resembled that of WT tissue, with TUBB3+ axons extending apically through the epithelial layers to reach the most superficial cells (Fig.2C,D). Thus, the reduced epithelial innervation at 5 wks is effectively restored through sustained lacripep treatment.

Next, we questioned the timing of epithelial reinnervation in response to lacripep treatment by analyzing the nerve supply at the 7-day time point (6 wks of age) through 3D imaging of intact whole mount corneas (Fig.3A). As shown in Figure 3B, PBS-treated *Aire* KO corneal epithelia showed a severe reduction in innervation at day 7, with the central cornea possessing only 18% of the nerve density of WT tissue (Fig.3C), an outcome that remained at this level over the 2 wk treatment window (see Fig.2). In contrast, corneas treated with lacripep exhibited significantly greater innervation at 7 days, reaching 45% of the nerve density of WT tissue (Fig.3B,C), with levels returning to that of WT cornea by day 15 (see Fig.2). Consistent with lacripep promoting nerve regeneration rather than nerve maintenance, lacripep-treated *Aire* KO corneas displayed a greater proportion of newly regenerating GAP43+ (TUBB3−) nerves relative to GAP43+TUBB3+ nerves than the WT corneas (Fig.3B,C). Specifically, the central corneas of lacripep-treated KO mice were populated with 50% GAP43+ nerves and 26% GAP43+TUBB3+ nerves while the WT tissue showed 43% and 37%, respectively (Fig.3C). Of note, the pro-regenerative cellular changes after axon injury result in neurons switching from an active, electrically transmitting state back to an electrically silent, growth-competent state (*29–31*), thereby likely impairing increased basal tear production at the 7-day time point for lacripep treated mice compared to untreated *Aire* KO controls (Fig.1B).

**Fig. 3.**
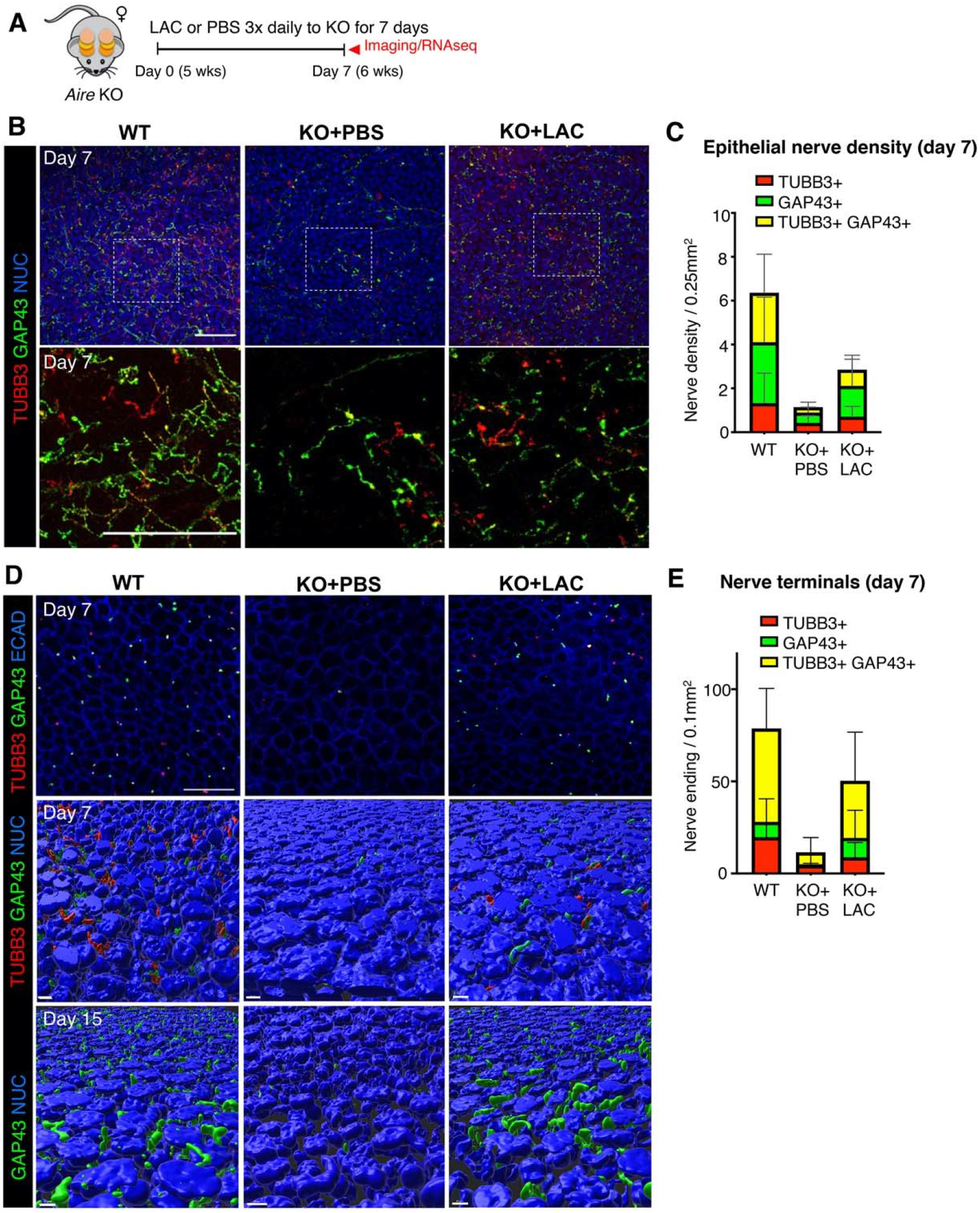
Lacripep re-establishes functional corneal nerve-epithelial interactions during dry eye disease progression. **A**, Schematic of the treatment regimen showing data collection time points for RNAseq and tissue analyses. **B-C**, Immunofluorescent analysis (**B**) and quantification (**C**) of regenerating nerves in the corneal epithelium at day 7. Graph in **C** shows the relative proportion of newly regenerating GAP43+ nerves in the central cornea. **D**, 3D reconstruction of whole mount immunofluorescent images of newly regenerating (GAP43+) and existing (TUBB3+) intraepithelial nerve terminals at day 7 (upper panel) and 15 (lower panel) in the corneal epithelium. **E**, Graph showing relative proportion of GAP43+, GAP43+TUBB3+ and TUBB3+ nerve terminals in the central cornea at day 7. NUC = nuclei. Scale bar = 25 μm, 10 μm. Each dot in the bar graphs represents a biological replicate. All data are expressed as mean + s.d. n > 4 mice per group.

We next determined the extent of functional reinnervation of the cornea at day 7 by assessing the distribution of sensory nerve endings (nerve terminals) within the epithelial layers in lacripep-treated *Aire* KO corneas compared to WT and PBS KO controls (Fig.3D). Sensory nerve terminals within the corneal epithelium respond to changes in the thickness and stability of the tear film to promote basal tear secretion, while also triggering epithelial responses to restore homeostasis through the release of neurotransmitters. Using 3D imaging and topographical reconstruction by IMARIS, we identified an extensive array of intraepithelial sensory nerve terminals within the WT controls (78 per 0.1mm^2^) contrasting with the severely reduced numbers found in PBS-treated *Aire* KO (Fig.3D,E). However, lacripep-treated *Aire* KO epithelia showed a significant increase in nerve terminals (50 per 0.1mm^2^), indicating nerve-epithelial cross-talk was being re-established. Consistent with the activation of nerve regeneration rather than nerve maintenance, the proportion of the sensory nerve terminals that expressed GAP43, as opposed to TUBB3 alone, in the lacripep-treated corneas was significantly greater than WT (p<0.05) and PBS-treated tissue (p<0.05; Fig.3D,E). Moreover, at day 15 the total number of nerve terminals in the lacripep treated *Aire* KO cornea epithelium was similar to that of WT controls highlighting the extensive recovery of innervation taking place during disease progression (Fig.3D). Collectively, our data illustrate the therapeutic efficiency of lacripep as the first topical dry eye treatment capable of regenerating functional corneal nerves and nerve-epithelial connections required for ocular integrity, signaling, and tear secretion.

### Lacripep activates master regulators of nerve regeneration

Finally, we questioned the potential mechanism by which lacripep restores the nerve supply. Lacritin and lacripep bind and activate syndecan-1 (SDC1), a transmembrane heparan sulfate proteoglycan enriched in the corneal epithelium, which has been shown to increase epithelial cell proliferation and migration in vitro (*32–33)*. SDC1 is also heavily involved in corneal nerve regeneration after injury (*34*), with *Sdc1*-deficient mice exhibiting a profound reduction in corneal innervation after ocular damage that is subsequently followed by corneal pathologies consistent with dry eye (*34*). Thus, we first confirmed that SDC1 remains expressed in the epithelium during dry eye disease progression through application of in-situ hybridization (RNAscope) to healthy and diseased corneas at 7 wks of age. As shown in Figure 4A, *Sdc1* transcripts were highly enriched in the basal and suprabasal cell populations of the WT corneas in a manner consistent with previous studies (*35*). Similarly, we found *Sdc1* transcripts were also abundantly located in the basal and suprabasal cells of the *Aire* KO corneal epithelium (Fig.4A), thus demonstrating that SDC1-lacripep interactions within the epithelium can take place during dry eye disease progression.

**Fig. 4.**
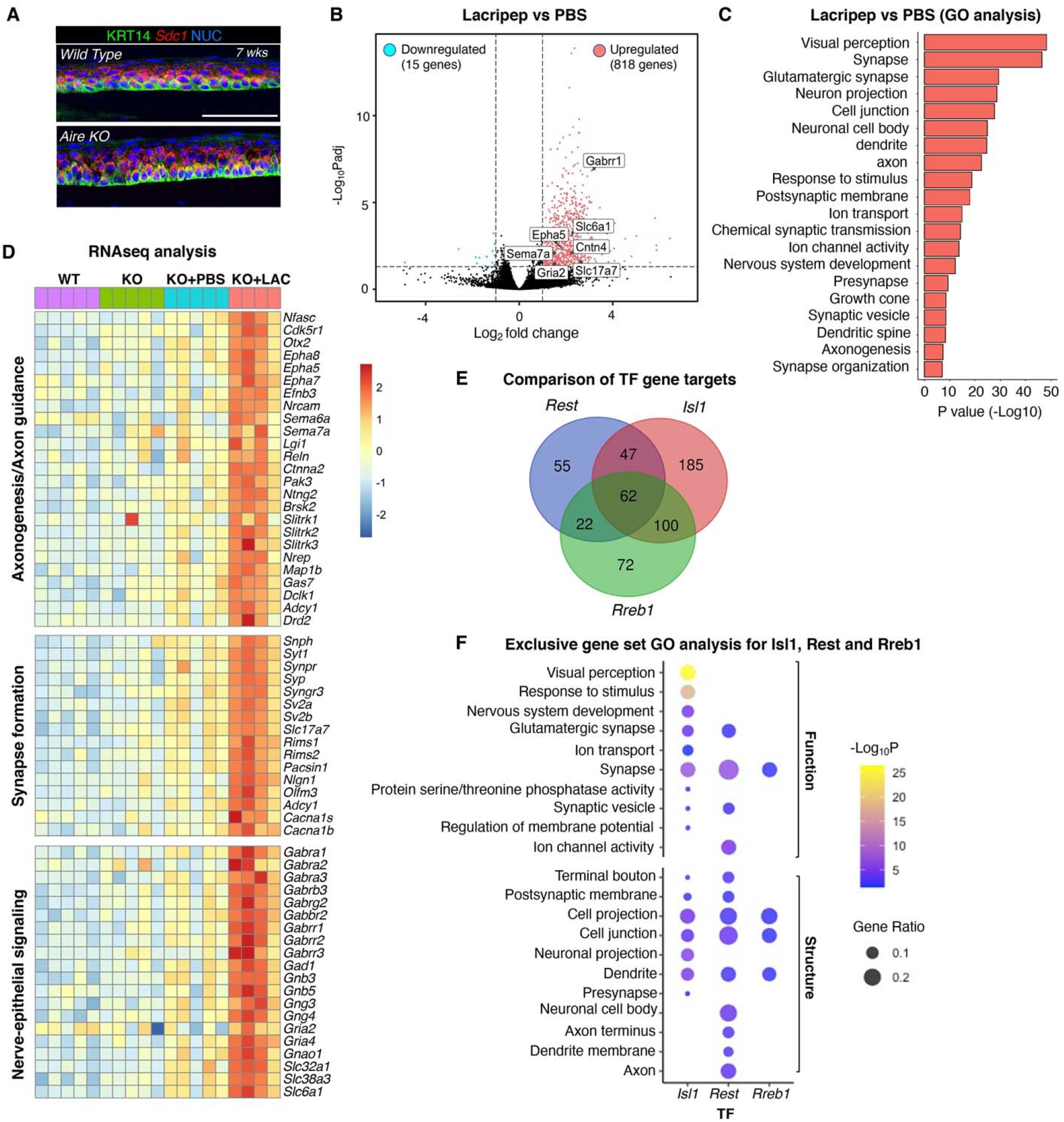
Lacripep activates master regulators of nerve regeneration in the desiccated cornea. **A**, RNA transcripts for syndecan-1 (*Sdc1*), co-receptor for lacripep, in the basal and suprabasal epithelial cells of WT and Aire KO cornea at 7 wks of age. Scale bar = 50 μm. **B**, Volcano plot of differentially expressed corneal genes in response to lacripep versus PBS treatment at day 7. **C**, Gene Ontology (GO) analysis highlighting upregulated pathways in lacripep versus PBS-treated *Aire* KO corneas. **D**, Heatmap featuring lacripep-induced upregulated genes associated with axonogenesis, axon guidance, nerve-epithelial signaling and synapse formation at day 7 of treatment. **E**, Venn diagram of the top transcription factors *Isl1, Rest* and *Rreb1* regulon targets upregulated in response to lacripep. Normalized enrichment score (NES) > 3, # Targets >100. **F**, GO analysis of the exclusive target gene sets for each TF in **E**, highlighting individual and combined roles in the regulation of function-and structure-based neuronal processes. p<0.05.

To identify potential downstream targets of lacripep treatment, we elucidated changes in corneal gene expression and signaling pathways at the day 7 time point through RNAseq. Principal component analysis (PCA) identified major transcriptional differences between each treatment group (Fig.S2A). Strikingly, differential gene expression analysis (DESeq2 (*36*); *P*adj<0.05; 2-fold change cutoff) of lacripep-versus PBS-treated corneas revealed an extensive alteration in gene expression that was predominantly associated with increased transcription, with 818 genes being significantly upregulated while only 15 were downregulated (Fig.4B, Table S2). Subsequent analysis yielded enriched gene sets that were highly associated with neuronal gene ontology (GO) terms e.g., synapse (P = 3.6E-47), visual perception (P = 5.8E-49), neuron projection (P = 2.1E-29), and response to stimulus (P = 1.8E-19) (Fig.4C, Data S2). These included cohorts of genes expressed by nerves and/or corneal epithelial cells involved in axonogenesis and axon guidance e.g., *Neurod2, Cntn4*, ephrins (*Epha5, Epha7, Epha8*) and semaphorins (*Sema6a*; *Sema7a*); synapse formation and function e.g., synaptic vesicle membrane proteins (*Syt1*; synaptophysin-*Syp*) and synaptic vesicle exocytosis regulators (*Rims1, Rims2*); neuronal signaling e.g., GABA receptors (*Gabra2, Gabrr1*), glutamate receptors (*Gria2, Gria4)*, and G protein signal-transducing mediators (*Gnb3, Gng4*); and neurotransmitter transport e.g. glutamate transporters (*Slc17a7*) and GABA transporters (*Slc6a1, Slc32a1*) (Fig.4D, Table S3). Analysis of corneas treated with lacripep or PBS versus untreated corneas further emphasized the ability of lacripep and not PBS alone to induce transcription of neuronal gene sets (Fig.S2B). Combined, these data clearly demonstrate that lacripep positively promotes neuroregeneration by upregulating a gene signature associated with axon migration, synapse formation and neuronal-epithelial signaling.

To identify candidate transcription factors that regulate these gene sets, we performed transcription factor (TF)-gene target regulatory network analysis using iRegulon (*37*). iRegulon identifies transcription factor-binding motifs that are enriched in the genomic regions of a query gene set and predicts transcription factors that bind to them. This revealed an abundant up-regulation of target gene sets from 3 top master transcriptional regulators, RE1 Silencing Transcription Factor (*Rest*, 184 genes), ISL LIM Homeobox 1 (*Isl1*, 394 genes), and Ras-Responsive Element-Binding Protein 1 (*Rreb1*, 254 genes) (Fig.S2C, Fig.4E, Table S3). ISL1 plays an essential role in the generation of sensory and sympathetic neurons (*38 39*), REST is a master regulator of neurogenesis that plays a role in modulating synaptic plasticity (*40–42*) and RREB1 regulates axon injury (*43*). Notably, we identified a significant number of genes per gene set that are uniquely regulated by 1 of the 3 different TFs (Fig 4E, Table S4). This was particularly the case for *Isl1*, a TF previously shown to be enriched in the limbal cells (*44*) and corneal sensory nerves (*45*), where 47% of gene targets (185 out of 394) did not overlap with the other TFs, compared to 29% and 28% for *Rest and Rreb1*, respectively (Fig.4E, Table S3). The *Isl1* exclusive gene set was also extensively enriched in genes involved with functional and structural nerve-mediated processes including visual perception (P=3.5E-27, e.g., *Capb4, Vsx1*), response to stimulus (P=9.7E-19; e.g., *Slc17a7, Ush2a*), synapse (P=9.4E-8, e.g. *Magi2, Synpr*) and neuron projection (P=6.4E-6, e.g., *Cdh23, Kif1a*) (Fig.4F, Table S4). Although the identified gene sets found to be exclusively regulated by *Rest* and *Rreb1* were less extensive than that of Isl1, each of the 3 TFs regulated specific genes involved in synapse formation and function (Fig.4F), suggesting that *Isl1, Rest* and *Rreb1* jointly regulate nerve-epithelial communication in response to lacripep.

Thus, together these data suggest that lacripep regenerates sensory nerves and re-establishes nerve-epithelial communication during dry eye disease progression through upregulation of master regulators of neuronal-associated gene sets.

## Discussion

To date, none of the dry eye therapies, including those currently in clinical trials, have been shown to promote the comprehensive restoration of corneal structure and function at the cellular and tissue level in pre-clinical animal models. Indeed, current medications for multiple ocular surface diseases serve to inhibit inflammation but fail to act on the tissue itself to retain or regain structure and function. Here, we demonstrate that lacripep treatment reverses the multifaceted pathogenesis of dry eye disease through its prosecretory, pro-regenerative, and neurotrophic functions. Using a murine model that recapitulates many of the features of dry eye disease, we show that lacripep restores the structural and functional integrity of the cornea and basal progenitor cell identity by promoting functional reinnervation of the epithelium, thereby serving to disrupt disease development and support active wound healing. Furthermore, we show that this outcome occurs in the presence of chronic inflammation, thus indicating that the regeneration and reinnervation of inflamed tissues can occur in the diseased context.

Loss of corneal nerves is an established clinical consequence of dry eye pathogenesis in Sjögrens, diabetes (*46*), rheumatoid arthritis (*47*), scleroderma (*48*), thyroid associated disorders (e.g., thyroid-associated ophthalmopathy)(*49*) and chronic graft-versus-host disease (*50*). However, despite this, the clinically approved therapeutics utilized to treat dry eye to date are directed at dampening inflammation but none have been shown to promote the restoration of corneal nerves or to regenerate tissue. With recent studies showing a vital role for T-cell signaling pathways in promoting the stability of ocular surface homeostasis and the wound repair process (*51, 52*), reducing immune cell infiltration and/or altering their function may inhibit cornea/nerve regeneration. In contrast, the highly utilized anti-inflammatory drug cyclosporin A has been shown to retard the regeneration of surgically transected nerves (*53*). These outcomes have consequently fueled a growing interest in the development of novel therapies that target disrupted innervation. To date, several factors with neuroregenerative potential have been tested in animal models, including corneal matrix repair product cacicol (*54*), insulin growth factor-1 (*55*), the neuropeptide pituitary adenylate cyclase-activating polypeptide (*56*), pigment-epithelium derived factor (*57*), and *N*-(1-acetylpiperidin-4-yl)-4-fluorobenzamide (FK962)(*58*) but results have been limited. More recently in preclinical and clinical studies for treating neurotrophic keratitis, an uncommon degenerative disease of the cornea resulting in denervation, breakdown of the epithelium, ulceration and perforation, (*59, 60*), recombinant nerve growth factor (cenegermin) stimulated the reinnervation of rabbit corneas after refractive surgery (*61*) and boosted corneal wound healing in human patients. However, in controlled clinical studies it has failed to show significant benefit over vehicle (artificial tears) in terms of corneal sensitivity or visual acuity (*62*). Moreover, its direct contribution to the regeneration and reinnervation of corneas from patients with neurotrophic keratitis and whether it can resolve dry eye pathologies in the setting of chronic inflammation are yet to be addressed.

Our study highlights a novel and significant link between neuroregeneration and the restoration of progenitor cell identity in a model of chronic, autoimmune-mediated dry eye. Multiple studies in other organ systems, such as the salivary glands (*63*), tooth (*64*) and skin (*65, 66*), have established a critical role for nerves in the regulation of tissue structure, and function, as well as progenitor cell maintenance. In the cornea, mouse models of both chemically-and physically-induced denervation (*67, 68*) have implicated sensory innervation as a regulator of progenitor cell identity, as determined by a reduction in the expression of epithelial progenitor cell markers such as p63 (*67*), an outcome that can be partially reversed by topical NGF treatment (*68*). Yet, the impact of denervation and reinnervation on the differentiation status of basal progenitors has not been addressed. Our data suggest that lacripep’s restorative effects on corneal innervation are mediated through its impact on epithelial cell identity/differentiation. Further studies are needed to identify the specific signaling systems activated by lacripep and the mechanism whereby sensory innervation regulates epithelial progenitor identity and cell fate.

Although the precise mechanism by which lacripep stimulates corneal reinnervation remains unclear, its known interaction with SDC1 suggests that lacripep mediates its neuroregenerative effects via binding and activating SDC1. Indeed, the nerve phenotype of the *Aire* KO cornea during dry eye disease progression strongly resembles that of the injured *Sdc1*-knockout cornea (*34*), as shown by the substantial reduction in intraepithelial nerve terminals and reinnervation. However, little is known about the downstream pathways that mediate neuroregeneration in response to SDC1. Our RNAseq analysis of the *Aire* KO cornea provides new insight into the potential mechanism of corneal reinnervation through the identification of 3 master regulators of the highly enriched neuronal gene sets, namely *Isl1, Rest*, and *Rreb1*, potentially directing lacripep-induced nerve remodeling, neurite outgrowth, and synapse formation and function. Specifically, these factors have been reported to orchestrate processes such as neurogenesis, synaptogenesis and/or axon regeneration (*38–43)*). Additionally, REST acts as a critical factor linking neuronal activity to the activation of intrinsic homeostasis and restoring physiological levels of activity throughout the entire neuronal network (*69)*. Furthermore, in addition to regulating innervation, ISL1, RREB1 and REST are also involved in stem cell/progenitor maintenance, epithelial differentiation and proliferation, as well as epithelial architecture and integrity (*70–74*), suggesting lacripep delivers regenerative instructions to the epithelium to coordinate the regulation of tissue structure and cell identity. Thus, lacripep may act by fine-tuning the nerves and the epithelium via these TFs to achieve functional reinnervation.

In summary, our study highlights lacripep as a new therapeutic capable of resolving ocular damage through promoting functional reinnervation of the cornea. Future studies exploring lacripep’s effects at the cellular level will allow us to identify and target specific signaling pathways essential for corneal re-innervation and restoration in patients with dry eye and other vision-threatening ocular surface disorders that impact corneal nerves (e.g., herpes simplex virus, interstitial keratitis, and neurotrophic keratitis).

## Materials and Methods

### Animal Model

All procedures were approved by the UCSF Institutional Animal Care and Use Committee (IACUC) and adhered to the NIH Guide for the Care and Use of Laboratory Animals (Approval number: 332AN089075-02). Wild type (WT) and *Aire*-deficient mice on the BALB/c background (BALB/c *Aire* KO) were the gift of Mark Anderson, University of California, San Francisco. Adult female mice (aged between 5 and 7 weeks) were used in all experiments. Mice were housed in the University of California, San Francisco Parnassus campus Laboratory Animal Resource Center (LARC), which is AAALAC accredited. Mice were housed in groups of up to five per cage where possible, in individually ventilated cages (IVCs), with fresh water, regular cleaning, and environmental enrichment. Appropriate sample size was calculated using power calculations. Genomic DNA isolated from tail clippings was genotyped for the Aire mutations by PCR with the recommended specific primers and their optimized PCR protocols (Jackson Laboratories Protocol 17936).

### Treatment Regimen

*Aire KO* female mice aged 5 weeks were used for the study. All mice were topically treated with 5 μL per eye of 4 μM lacripep. Dosing was three times daily for 14 consecutive days. Lissamine green staining assessment and tear secretion assays were performed on each eye before treatment at baseline, 7 days and 14 days post treatment

### Analysis of epithelial barrier function

After mice were anesthetized with isoflurane, 5 μL of lissamine green dye (1%) was applied to the lower conjunctival cul-de-sac. Images of the cornea were then taken using an Olympus Zoom Stereo Microscope (Olympus, CenterValley, PA). Lissamine green staining was scored by dividing the cornea into four quadrants, the extent of staining in each quadrant was classified as Grade 0, no staining; Grade 1, sporadic (<25%); Grade 2, diffuse punctate (25-75%), or Grade 3, coalesced punctate staining (75% or more). The total score was calculated separately for each eye and equaled the sum of all four quadrants ranging from 0 (no staining) to 12 (most severe staining). Scoring was conducted by three masked observers with each data point representing the fold change of each eye relative to its baseline score before treatment.

### Tear Secretion Measurements

Mice were anesthetized with isoflurane and basal tear secretion was then measured using a Zone-Quick phenol red thread (as indicated by the length of the tear-absorbed region in 15 seconds). Stimulated tear secretion was measured after 4.5 mg/kg of pilocarpine diluted in saline was injected into the peritoneum (i.p.). Ten minutes later, mice were anesthetized with isoflurane and tear secretion was measured using a Zone-Quick phenol red thread (Showa Yakuhin Kako Co. Ltd., Tokyo, Japan).

### Tissue processing and immunohistological analyses

Immunohistological and immunofluorescent analyses of cornea and lacrimal gland samples were performed as previously described (*8*). Briefly, enucleated eyes were embedded in OCT Tissue Tek freezing media. 7μm and 20 μm sections were prepared from fresh frozen tissues using a cryostat (Leica, lzar, Germany) and mounted on SuperFrost Plus slides. Sections were fixed for 20 min in 4% paraformaldehyde (PFA) at room temperature (RT) and permeabilized using 0.3% Triton X100 in phosphate buffered saline for 15 min. Sections were then washed in PBS-Tween 20 (PBST) for 10 min, before being blocked with 5% normal donkey serum (Jackson Laboratories, ME) in PBST for 1 hour at RT. After blocking, slides were incubated with primary antibodies diluted in blocking buffer overnight at 4°C. Following 3 washes with PBST, slides were incubated with secondary antibodies diluted in blocking solution at RT for 1 hour.

For immunofluorescent analysis, 7 or 20 μm tissue sections were incubated with the following primary antibodies: mouse anti-ZO1 conjugated to Alexa594 (1:1000, Life Technologies, Cat 339194), rabbit anti-KRT6A (1:800, Cell Signaling, Cat 4912S), chicken anti-KRT14 (1:1500, Santa Cruz, Cat 515882), rat anti-Ki67 (1:200, Biolegend, 652405), rat anti-CD4 (1:200, Santa Cruz), rabbit anti-TUBB3 (1:500, Cell Signaling, Cat 5568S), and rat anti-Ecadherin (1:300, Life Technologies, Cat 131900). Antibodies were detected using Cy2-, Cy3-or Cy5-conjugated secondary Fab fragment antibodies (Jackson Laboratories), and nuclei were stained with Hoechst 33342 (1:3000, Sigma-Aldrich). Fluorescence was analyzed using a Zeiss LSM 900 confocal microscope or Zeiss Yokogawa Spinning disk confocal microscope with images assessed using NIH ImageJ software, as described below.

For corneal whole mount staining, PFA fixed corneas were blocked and incubated in primary antibodies for 48hr at 4□: Mouse anti-TUBB3 (1:300, R&D, MAB 1195), rabbit anti-GAP43 (1:400, Novus bio, NB 300-143), goat anti-CGRP (1:300, Thermo fisher, PA1-85250), and rat anti-Substance P (1:500, Millipore, MAB 356). This was followed by extensive washes in PBST before tissue was incubated in secondary antibodies overnight at 4□.

### *In situ* Hybridization

Manual chromogenic RNAscope (ACDBio) was performed using company protocols on fresh frozen cornea tissue sections to detect target RNA at single cell level. Tissue pre-treatment included fixation for 15 min in 4% paraformaldehyde (PFA) at 4°C, RNAscope® Hydrogen Peroxide (ACD# 322335) treatment for 10 min at RT followed by protease treatment (RNAscope® Protease Plus ACD# 322331) for 10 min at 40□ using the HybEZ Oven. Detection of specific probe binding sites was with RNAscope Reagent kit— from ACD (Cat. 323110). Single ISH detection for mouse *Sdc1* (ACD Probe: 813921), Mouse Positive Control Probe (ACD Probe: 320881) and Negative Control Probe (ACD Probe: 320871) was performed manually. Target probes were hybridized for 2 hr at 40□ using HybEZ oven followed by amplification steps according to the manufacture’s protocol. Positive staining was indicated by fluorescent dots in the cell cytoplasm or nucleus.

### Image Analysis

Quantification of tight junction protein ZO1. ZO-1 fluorescent intensity was quantified in the apical squamous epithelial layer within a region of interest (ROI) containing a 350 µm section of central cornea epithelium. A Tsai’s thresholding method (*75*) (Moments) was then applied to the ROI, and integrated densities within the ROI of the thresholded image were recorded.

Quantification of corneal epithelial basal progenitor identity and proliferation. The number of basal KRT14+ and KRT14+KRT6A+ basal epithelial cells were counted in 350 µm sections of central cornea. To obtain the percentage of KRT6A+ basal cells per respective region, the number of KRT14+KRT6A+ basal cells was divided by the total number of KRT14+ basal cells. The number of proliferating epithelial progenitors (Ki67+KRT14+) was presented as a percentage of total KRT14+ cells counted in 350 µm sections of central cornea.

Quantification of CD4+ T cells in cornea and Lacrimal gland. The numbers of CD4+ T cells in the limbal stroma were counted and graphed. CD4+ T cells in the lacrimal gland were presented as percent coverage of total CD4+ T cells within the whole lacrimal gland tissue area.

Quantification of corneal epithelial nerve density and synapses. Tissue sections: TUBB3+ nerve density within 300 µm sections of central cornea epithelial ROI was quantified by applying Tsai’s thresholding method (Moments), with integrated densities within the ROI of the thresholded image being recorded. Corneal whole mounts: The density of nerves expressing GAP43, TUBB3, or GAP43 and TUBB3 (% Area of 354 µm x 396 µm, double positive nerve % Area was calculated from colocalization threshold image) was quantified from central cornea images of whole mount cornea and plotted in a stacked bar graph to visualize the proportion of each of the three nerve types. Similarly, numbers of synapses marked by expression of GAP43, TUBB3, or GAP43 and TUBB3 were counted per respective region of central cornea (100 µm^2^) and plotted in a stacked bar graph. The density of GAP43+ nerves were quantified from the whole mount cornea images (354 µm x 396 µm) applying Tsai’s threshold as described above.

3D reconstruction of nerve fibers. Acquired confocal Z-stacks comprising epithelial and stroma were reconstructed into 3D images using Imaris image analysis software (Bitplane AG, Zurich, Switzerland). The surface of the nuclei (Hoechst) and nerves (TUBB3/GAP43/SP/CGRP) were segmented using the background subtraction threshold. Each channel was obtained separately. Touching nuclei were split by the calculated seed points inside each area.

### RNA isolation and RNAseq library generation

Total RNA was collected at 7 days of treatment and purified using RNAaqueous and DNase reagents according to the manufacturer’s instructions (Ambion, Houston, TX, USA). RNA quality was assessed using the Agilent 2100 BioAnalyzer, and samples with an RNA integrity ≥ 6.0 were included for RNA sequencing. The synthesis of mRNA libraries was performed by Novogene Corporation Inc. according to their protocols. RNA library was formed by ployA capture (or rRNA removal), RNA fragmentation by covaris or enzyme digestion and reverse transcription of cDNA. Sequencing was performed as described below.

### RNAseq analysis

RNA libraries were sequenced on an Illumina NovaSeq 6000. Depths of 20–30 million 150 bp paired-end reads were generated for each sample. Quality control metrics were performed on raw sequencing reads using the FASTQC v0.11.9 application (*76*). Reads were mapped to the UCSC Mus musculus genome mm10 (NCBI build v38) using Spliced Transcripts Alignment to a Reference (STAR) (*77*). At least 90% of the reads were successfully mapped. Reads aligning to the mm10 build were quantified against Ensembl Transcripts release 93 using Partek® E/M (Partek’s optimization of the expectation maximization algorithm, Partek Inc, St.Louis, MO, USA), which disregarded any reads that aligned to more than one location or more than one gene at a single location. Data was normalized by two procedures: 1. total count normalization, 2. addition of a small offset (0.0001). DEseq2 was then used to detect differential gene expression between WT, untreated *Aire* KO, PBS treated *Aire* KO and lacripep treated *Aire* KO corneas based on the normalized count data. Genes were considered differentially expressed if the log2 Fold Change between samples was at least 1, with the adjusted p-value held to 0.05 (*36*). Heatmaps and Volcano plot of differentially expressed genes were created using “pheatmap”, and “EnhancedVolcano” R packages, respectively (*78, 79*).

A list of all significantly modulated genes (p < 0.05) was used as input for gene ontology (GO) analysis using the online Database for Annotation, Visualization and Integrated Discovery (DAVID, 2021 update (*80*). We consider an attribute to be significant if its adjusted p value is less than 0.05 relative to an appropriate background gene set.

To identify regulatory networks underlying the impact of lacripep on tissue regeneration, we used the cytoscape application *iRegulon* (*37*) to predict master regulators, i.e., transcription factors whose regulons (transcriptional target sets) are highly enriched with the input gene list. A list of all significantly modulated genes (FC>1.5, p < 0.05) was used as input and the normalized enrichment score (NES) threshold was set to 3, which corresponds to an FDR of the TF recovery between 3%-9%. Exclusively upregulated genes for each TF were further used as input for GO analysis using DAVID as indicated above.

### Statistical Analysis

A minimum of three independent repeats were conducted in all experiments. Bar graphs are used to summarize the means and standard deviations of each outcome obtained using all data collected from wild type (WT) and *Aire KO* mice. A Student’s t-test was used for two group comparisons; a one-way analysis of variance (ANOVA) was used for multiple group comparisons with either Tukey’s multiple comparisons test, Dunnett T3, or corrected for multiple comparisons by controlling the False Discovery Rate using Benjamini, Krieger and Yekutieli; P < 0.05 was considered statistically significant. A false discovery rate of 0.05 was applied to RNAseq data. A Wald test with Benjamini and Hochberg correction was used for differential gene expression, and a Fisher’s exact test was used for Gene Ontology analysis.

### Study approval

All procedures were approved by the UCSF Institutional Animal Care and Use Committee (IACUC) and were adherent to the NIH Guide for the Care and Use of Laboratory Animals.

## Acknowledgments

The authors would like to thank Drs. John Whitelock, Alison May, Ophir Klein, Jeffrey Bush and Matilda Chan for their constructive input towards this manuscript as well as the sharing of reagents and instruments.

## Author contributions

Conceptualization: YE, FY, NAM, SMK

Methodology: YE, FY, KNC, EG, NAM, SMK

Investigation: YE, FY, KNC, EG

Visualization: YE, FY, KNC, SMK

Funding acquisition: SMK, NAM

Project administration: SMK, NAM

Supervision: SMK, NAM

Writing – original draft: YE, FY, NAM, SMK

Writing – review & editing: YE, FY, NAM, SMK

## Data and materials availability

Data are available in the main text or the supplementary materials. All transcriptomic data underlying the study will be deposited in Gene Expression Omnibus (GEO)

## Supplementary Materials (separate file)

Please see attached “Efraim, Chen et al., Supplementary materials.pdf” and data files.

