## Supplementary Figures and Tables for "A synthetic tear protein resolves dry eye through promoting corneal nerve regeneration"

**This PDF file includes:**

Figs. S1 to S2

Tables S1 to S4

**Other Supplementary Materials for this manuscript include the following:**

Data S1 to S2

**Fig. S1.**

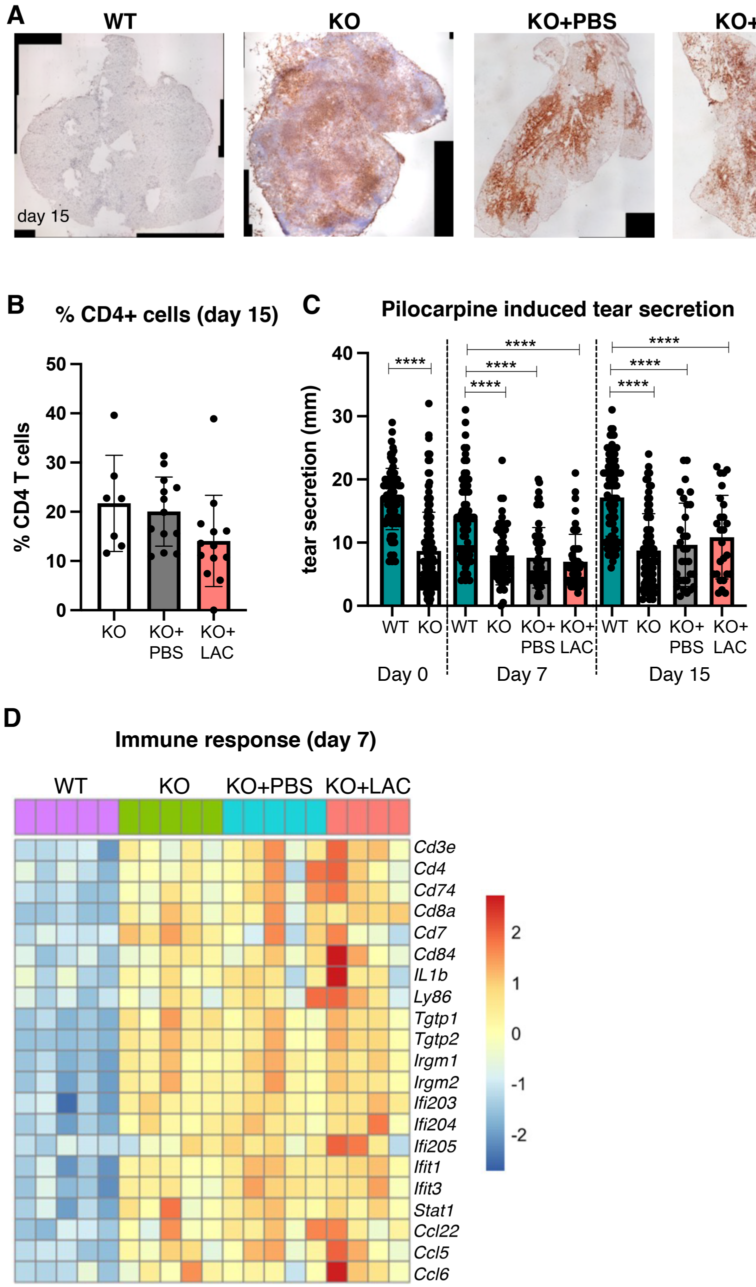

**Fig.S1. Lacripep improves tear secretion and barrier function despite chronic inflammation. A-B,** Immunohistological analysis and quantification of CD4+ T cells in WT, untreated and treated *Aire* KO lacrimal glands. Graph in **B** shows the percentage of CD4+ T cells in each treatment group normalized to WT controls. **C,** Levels of pilocarpine induced (maximal) tear secretion at day 7 and 15 compared to baselin**e. D,** Heatmap highlighting genes associated with immune responses are upregulated in all treatment conditions in the *Aire* KO cornea but not in the WT.*$p < 0.05$; **$p < 0.01$; ***$p < 0.001$; one-way analysis of variance was used in B; Student's T test and one-way analysis of variance with Tukey's test was used in C. Each dot in the bar graph represents a biological replicate. Error bars represent standard deviation. $n \geq 4$ mice per group.

**Fig. S2.**

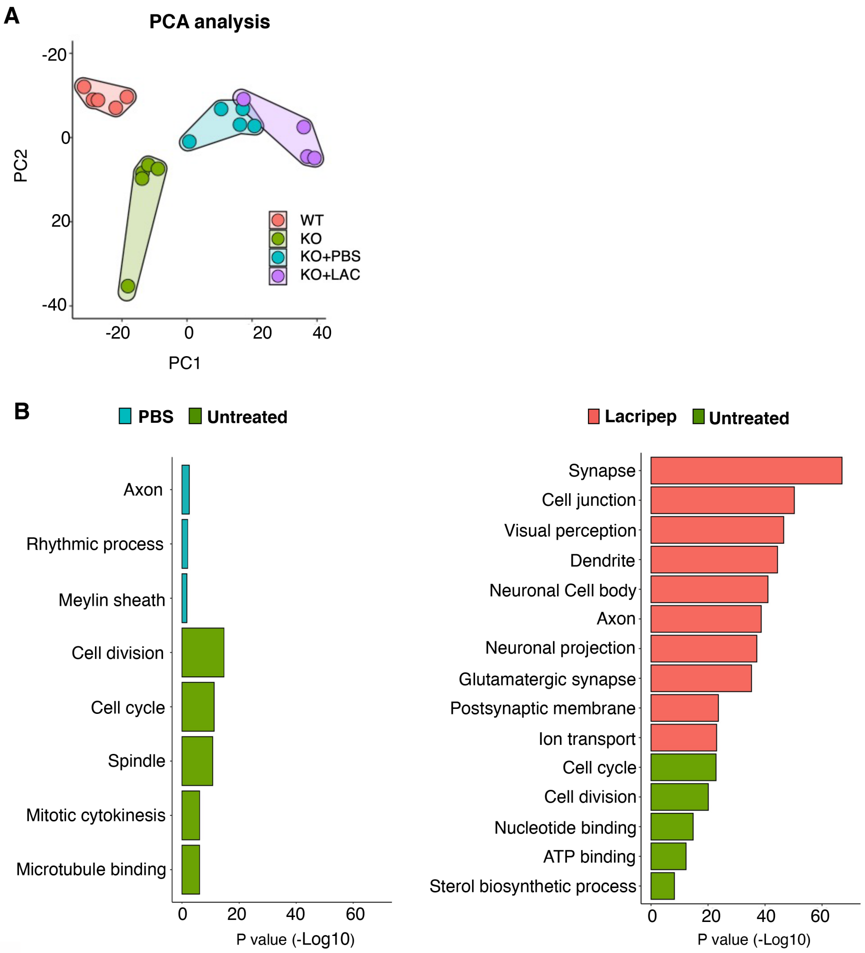

**C**

| TF | Motif | NES | #Targets |
|---|---|---|---|
| ***Rest*** | 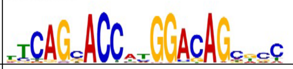 | 7.888 | 184 |
| ***Isl1*** | 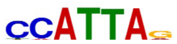 | 4.199 | 394 |
| ***Rreb1*** | 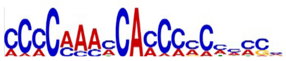 | 3.148 | 254 |

**Fig.S2. RNAseq analysis of cornea samples at 7 days of treatment**. (**A**) Principal component analysis (PCA) plot of the different treatment groups. (**B**) Gene Ontology (GO) analysis highlighting upregulated pathways in PBS- and lacripep-treated versus untreated *Aire* KO controls; FC>1.5,p<0.05. (**C**) Identification of top master transcriptional regulators enriched in lacripep-treated corneas when compared to PBS. iRegulon was employed on 886 upregulated genes. FC>1.5, p<0.05. Table shows binding motif, Normalized enrichment score (NES) and number of targets (# Targets). NES>3, # Targets>100.

**Table S1. Differential gene expression analysis of *Aire* KO vs WT corneas at 6 wks of age**

| ***Gene*** | **baseMean** | **log2FoldChange** | **lfcSE** | **stat** | **pvalue** | **padj** |
|---|---|---|---|---|---|---|
| *Sprr1b* | 910.84 | 11.13 | 1.58 | -7.05 | 1.77E-12 | 4.94E-10 |
| *Cxcl9* | 101.35 | 7.56 | 1.27 | -5.97 | 2.42E-09 | 2.60E-07 |
| *Gbp10* | 16.20 | 7.53 | 1.25 | -6.01 | 1.82E-09 | 2.04E-07 |
| *Gm4841* | 30.64 | 7.28 | 1.11 | -6.57 | 5.14E-11 | 9.25E-09 |
| *Gbp11* | 9.67 | 6.78 | 2.12 | -3.20 | 1.36E-03 | 1.49E-02 |
| *Tgtp2* | 151.73 | 6.72 | 0.51 | -13.26 | 3.80E-40 | 1.25E-36 |
| *Cxcl10* | 51.32 | 6.56 | 1.38 | -4.74 | 2.14E-06 | 8.82E-05 |
| *Iqcn* | 18.26 | 6.54 | 1.16 | -5.65 | 1.59E-08 | 1.38E-06 |
| *Apol6* | 7.78 | 6.47 | 1.22 | -5.29 | 1.19E-07 | 7.93E-06 |
| *Opn1mw* | 14.00 | 6.13 | 1.93 | -3.18 | 1.47E-03 | 1.57E-02 |
| *Tgtp1* | 287.87 | 5.98 | 0.76 | -7.89 | 2.98E-15 | 1.44E-12 |
| *Cxcl11* | 7.75 | 5.87 | 1.12 | -5.22 | 1.77E-07 | 1.10E-05 |
| *Gm12250* | 292.93 | 5.75 | 0.36 | -15.89 | 7.47E-57 | 7.39E-53 |
| *C1ql3* | 4.11 | 5.55 | 1.43 | -3.87 | 1.08E-04 | 2.19E-03 |
| *Ubd* | 9.20 | 5.52 | 1.09 | -5.09 | 3.68E-07 | 2.01E-05 |
| *Mmp13* | 201.43 | 5.49 | 1.01 | -5.43 | 5.58E-08 | 4.19E-06 |
| *Gm12946* | 5.95 | 5.49 | 1.20 | -4.58 | 4.72E-06 | 1.67E-04 |
| *Gja3* | 5.77 | 5.44 | 1.15 | -4.72 | 2.36E-06 | 9.56E-05 |
| *Gvin1* | 8.54 | 5.42 | 1.11 | -4.89 | 9.84E-07 | 4.62E-05 |
| *A930031H19Rik* | 5.60 | 5.40 | 1.44 | -3.75 | 1.79E-04 | 3.25E-03 |
| *A230001M10Rik* | 3.66 | 5.39 | 1.53 | -3.51 | 4.46E-04 | 6.60E-03 |
| *Apol10b* | 3.45 | 5.30 | 1.19 | -4.43 | 9.28E-06 | 2.96E-04 |
| *Gbx2* | 3.32 | 5.24 | 1.22 | -4.31 | 1.62E-05 | 4.76E-04 |
| *Gzmb* | 6.72 | 5.05 | 1.19 | -4.24 | 2.28E-05 | 6.26E-04 |
| *Gabrr3* | 2.85 | 5.03 | 1.59 | -3.16 | 1.58E-03 | 1.66E-02 |
| *Tcrg-C2* | 2.79 | 4.99 | 1.34 | -3.72 | 1.98E-04 | 3.52E-03 |
| *Atcayos* | 3.98 | 4.89 | 1.52 | -3.21 | 1.32E-03 | 1.47E-02 |
| *A930003A15Rik* | 5.96 | 4.88 | 1.25 | -3.90 | 9.76E-05 | 2.03E-03 |
| *BC023105* | 11.41 | 4.83 | 0.85 | -5.71 | 1.10E-08 | 9.86E-07 |
| *Lrrc30* | 3.82 | 4.83 | 1.75 | -2.76 | 5.78E-03 | 4.24E-02 |
| *Cck* | 2.47 | 4.82 | 1.33 | -3.62 | 2.90E-04 | 4.74E-03 |
| *Car4* | 2.34 | 4.74 | 1.60 | -2.96 | 3.08E-03 | 2.71E-02 |
| *Ly6i* | 2.33 | 4.73 | 1.52 | -3.12 | 1.84E-03 | 1.87E-02 |
| *Csn3* | 410.45 | 4.72 | 0.70 | -6.74 | 1.61E-11 | 3.29E-09 |
| *H2-Q6* | 519.94 | 4.69 | 0.29 | -16.09 | 3.03E-58 | 5.99E-54 |
| *Tmem236* | 2.25 | 4.68 | 1.55 | -3.03 | 2.44E-03 | 2.29E-02 |
| *Atf3* | 799.57 | 4.65 | 0.37 | -12.44 | 1.66E-35 | 4.10E-32 |

| | | | | | | |
|---|---|---|---|---|---|---|
| *Ifi27l2a* | 1451.60 | 4.63 | 0.58 | -8.01 | 1.17E-15 | 6.25E-13 |
| *H2-Q7* | 304.70 | 4.57 | 0.33 | -13.76 | 4.52E-43 | 2.24E-39 |
| *Klrc1* | 3.16 | 4.56 | 1.18 | -3.86 | 1.14E-04 | 2.28E-03 |
| *Gm4951* | 34.64 | 4.52 | 0.66 | -6.84 | 7.88E-12 | 1.68E-09 |
| *Lim2* | 53.96 | 4.51 | 0.94 | -4.81 | 1.54E-06 | 6.73E-05 |
| *Mmp25* | 13.58 | 4.49 | 0.74 | -6.06 | 1.36E-09 | 1.61E-07 |
| *Gm5970* | 2.99 | 4.47 | 1.54 | -2.90 | 3.74E-03 | 3.11E-02 |
| *Fasl* | 2.92 | 4.43 | 1.27 | -3.50 | 4.64E-04 | 6.78E-03 |
| *Slc34a3* | 10.21 | 4.43 | 0.95 | -4.65 | 3.34E-06 | 1.28E-04 |
| *Nkg7* | 8.49 | 4.42 | 0.92 | -4.80 | 1.61E-06 | 7.03E-05 |
| *Dusp27* | 4.20 | 4.35 | 1.44 | -3.02 | 2.57E-03 | 2.37E-02 |
| *Pon1* | 15.38 | 4.31 | 0.86 | -5.03 | 5.02E-07 | 2.60E-05 |
| *Gulo* | 4.06 | 4.30 | 1.26 | -3.42 | 6.34E-04 | 8.62E-03 |
| *Cd8a* | 13.85 | 4.29 | 0.71 | -6.02 | 1.71E-09 | 1.93E-07 |
| *Ebf2* | 2.57 | 4.25 | 1.41 | -3.02 | 2.56E-03 | 2.37E-02 |
| *Prdm13* | 3.83 | 4.24 | 1.36 | -3.12 | 1.78E-03 | 1.82E-02 |
| *Erich3* | 3.92 | 4.24 | 1.52 | -2.79 | 5.25E-03 | 3.96E-02 |
| *Tcap* | 126.53 | 4.21 | 1.56 | -2.70 | 6.94E-03 | 4.84E-02 |
| *Lmod3* | 3.74 | 4.20 | 1.55 | -2.71 | 6.73E-03 | 4.75E-02 |
| *Mmp10* | 91.02 | 4.20 | 0.70 | -6.00 | 1.96E-09 | 2.18E-07 |
| *Nlrc5* | 372.28 | 4.20 | 0.28 | -15.08 | 2.30E-51 | 1.52E-47 |
| *Cst7* | 2.47 | 4.19 | 1.30 | -3.22 | 1.30E-03 | 1.45E-02 |
| *Fhl2* | 53.87 | 4.17 | 0.50 | -8.36 | 6.25E-17 | 4.13E-14 |
| *Fgf16* | 2.45 | 4.17 | 1.50 | -2.78 | 5.46E-03 | 4.08E-02 |
| *Lctl* | 27.44 | 4.13 | 1.10 | -3.77 | 1.64E-04 | 3.04E-03 |
| *Cntn5* | 5.38 | 4.09 | 1.35 | -3.03 | 2.46E-03 | 2.31E-02 |
| *Rd3* | 30.68 | 4.09 | 1.06 | -3.86 | 1.14E-04 | 2.28E-03 |
| *Sprr2h* | 22.28 | 4.03 | 0.90 | -4.47 | 7.83E-06 | 2.55E-04 |
| *Rnf165* | 5.94 | 4.02 | 1.10 | -3.64 | 2.71E-04 | 4.49E-03 |
| *BC147527* | 3.23 | 3.97 | 1.26 | -3.14 | 1.69E-03 | 1.75E-02 |
| *Hist1h1a* | 12.74 | 3.97 | 1.01 | -3.91 | 9.23E-05 | 1.95E-03 |
| *Gbp5* | 61.00 | 3.96 | 0.53 | -7.48 | 7.68E-14 | 3.04E-11 |
| *Rcvrn* | 135.54 | 3.95 | 1.23 | -3.21 | 1.31E-03 | 1.46E-02 |
| *Ngf* | 148.72 | 3.92 | 0.86 | -4.55 | 5.36E-06 | 1.86E-04 |
| *F830016B08Rik* | 16.55 | 3.92 | 0.81 | -4.83 | 1.39E-06 | 6.12E-05 |
| *Pde6h* | 11.87 | 3.91 | 1.09 | -3.59 | 3.33E-04 | 5.28E-03 |
| *Gngt1* | 106.88 | 3.89 | 1.01 | -3.87 | 1.08E-04 | 2.19E-03 |
| *Iigp1* | 720.06 | 3.87 | 0.59 | -6.59 | 4.42E-11 | 8.03E-09 |
| *Sprr1a* | 3015.66 | 3.87 | 0.92 | -4.23 | 2.34E-05 | 6.33E-04 |

| | | | | | | |
|---|---|---|---|---|---|---|
| *Klk7* | 10.82 | 3.84 | 0.99 | -3.90 | 9.75E-05 | 2.03E-03 |
| *Slamf8* | 8.23 | 3.84 | 0.80 | -4.78 | 1.73E-06 | 7.43E-05 |
| *Gm42743* | 140.33 | 3.83 | 0.87 | -4.39 | 1.14E-05 | 3.54E-04 |
| *Serpina3g* | 55.72 | 3.82 | 0.52 | -7.36 | 1.90E-13 | 6.50E-11 |
| *Casp1* | 95.92 | 3.82 | 0.29 | -13.11 | 2.72E-39 | 7.68E-36 |
| *Lgsn* | 142.17 | 3.81 | 0.89 | -4.28 | 1.85E-05 | 5.30E-04 |
| *Htr3a* | 8.68 | 3.77 | 1.15 | -3.28 | 1.04E-03 | 1.23E-02 |
| *Gm8989* | 4.33 | 3.76 | 1.30 | -2.90 | 3.74E-03 | 3.11E-02 |
| *Klhl30* | 7.67 | 3.76 | 1.12 | -3.35 | 7.95E-04 | 1.02E-02 |
| *Gm8979* | 12.77 | 3.74 | 0.68 | -5.53 | 3.20E-08 | 2.55E-06 |
| *Ptgs2* | 942.51 | 3.72 | 0.79 | -4.72 | 2.35E-06 | 9.53E-05 |
| *Card11* | 7.67 | 3.70 | 0.95 | -3.90 | 9.49E-05 | 1.99E-03 |
| *Dlx2* | 6.36 | 3.70 | 0.93 | -3.96 | 7.40E-05 | 1.63E-03 |
| *Retnla* | 7.49 | 3.67 | 0.98 | -3.73 | 1.93E-04 | 3.45E-03 |
| *Tspan32* | 3.69 | 3.67 | 1.26 | -2.90 | 3.76E-03 | 3.12E-02 |
| *4930546K05Rik* | 3.88 | 3.66 | 1.21 | -3.03 | 2.47E-03 | 2.31E-02 |
| *Gm13111* | 3.83 | 3.66 | 1.35 | -2.72 | 6.61E-03 | 4.68E-02 |
| *Ankrd33* | 29.18 | 3.66 | 1.22 | -3.00 | 2.66E-03 | 2.43E-02 |
| *Fosl1* | 254.18 | 3.64 | 0.65 | -5.64 | 1.66E-08 | 1.43E-06 |
| *Slamf6* | 5.25 | 3.64 | 0.97 | -3.75 | 1.74E-04 | 3.19E-03 |
| *Prph2* | 612.50 | 3.63 | 1.14 | -3.17 | 1.50E-03 | 1.60E-02 |
| *Crhr2* | 4.87 | 3.63 | 1.25 | -2.91 | 3.63E-03 | 3.04E-02 |
| *Rbp3* | 661.02 | 3.61 | 1.13 | -3.21 | 1.35E-03 | 1.49E-02 |
| *Pde6g* | 275.58 | 3.60 | 1.05 | -3.43 | 6.01E-04 | 8.32E-03 |
| *Cd3d* | 8.30 | 3.58 | 0.83 | -4.29 | 1.76E-05 | 5.11E-04 |
| *Crygd* | 911.80 | 3.57 | 0.84 | -4.26 | 2.07E-05 | 5.80E-04 |
| *Cryge* | 220.54 | 3.57 | 0.82 | -4.34 | 1.45E-05 | 4.34E-04 |
| *Mylk2* | 12.36 | 3.57 | 1.15 | -3.10 | 1.96E-03 | 1.95E-02 |
| *Rho* | 4082.53 | 3.56 | 1.14 | -3.12 | 1.81E-03 | 1.84E-02 |
| *Cd6* | 16.37 | 3.55 | 0.57 | -6.25 | 4.04E-10 | 5.67E-08 |
| *Ccl5* | 20.61 | 3.54 | 0.67 | -5.29 | 1.22E-07 | 8.03E-06 |
| *Slc17a7* | 268.92 | 3.53 | 1.06 | -3.33 | 8.57E-04 | 1.06E-02 |
| *Gjd2* | 10.81 | 3.52 | 0.83 | -4.23 | 2.30E-05 | 6.30E-04 |
| *Crygc* | 697.52 | 3.51 | 0.87 | -4.04 | 5.38E-05 | 1.26E-03 |
| *Gnat1* | 1543.56 | 3.50 | 1.07 | -3.27 | 1.08E-03 | 1.27E-02 |
| *Crybb1* | 438.92 | 3.49 | 0.85 | -4.11 | 3.90E-05 | 9.63E-04 |
| *Mc1r* | 4.57 | 3.48 | 1.09 | -3.18 | 1.48E-03 | 1.58E-02 |
| *Gp5* | 4.53 | 3.47 | 1.23 | -2.81 | 4.90E-03 | 3.77E-02 |
| *AW112010* | 40.22 | 3.47 | 0.47 | -7.36 | 1.79E-13 | 6.22E-11 |

| | | | | | | |
|---|---|---|---|---|---|---|
| *Pmaip1* | 77.64 | 3.46 | 0.77 | -4.49 | 7.01E-06 | 2.33E-04 |
| *Kcnj14* | 63.35 | 3.45 | 1.17 | -2.95 | 3.21E-03 | 2.79E-02 |
| *Fabp12* | 14.47 | 3.45 | 0.91 | -3.81 | 1.41E-04 | 2.68E-03 |
| *Six1* | 8.14 | 3.45 | 0.89 | -3.86 | 1.14E-04 | 2.28E-03 |
| *Hist4h4* | 4.47 | 3.44 | 1.17 | -2.93 | 3.34E-03 | 2.87E-02 |
| *Gimap5* | 4.28 | 3.44 | 1.03 | -3.34 | 8.29E-04 | 1.04E-02 |
| *Nrxn3* | 13.53 | 3.44 | 0.83 | -4.12 | 3.78E-05 | 9.37E-04 |
| *Rgr* | 174.79 | 3.43 | 0.71 | -4.82 | 1.42E-06 | 6.22E-05 |
| *Vtn* | 104.50 | 3.43 | 1.06 | -3.22 | 1.29E-03 | 1.44E-02 |
| *Slfn5* | 325.65 | 3.42 | 0.36 | -9.52 | 1.80E-21 | 2.23E-18 |
| *Gm2895* | 3.38 | 3.42 | 1.27 | -2.69 | 7.07E-03 | 4.90E-02 |
| *Gbp6* | 143.05 | 3.42 | 0.41 | -8.30 | 1.02E-16 | 6.53E-14 |
| *9330175E14Rik* | 15.45 | 3.41 | 0.59 | -5.81 | 6.09E-09 | 5.83E-07 |
| *Fam167a* | 55.80 | 3.41 | 0.84 | -4.06 | 4.96E-05 | 1.18E-03 |
| *Rpl3l* | 25.82 | 3.41 | 1.26 | -2.71 | 6.79E-03 | 4.77E-02 |
| *Has1* | 5.39 | 3.40 | 0.94 | -3.64 | 2.78E-04 | 4.58E-03 |
| *Ly6c2* | 7.50 | 3.40 | 0.77 | -4.41 | 1.05E-05 | 3.30E-04 |
| *Nxnl1* | 37.84 | 3.40 | 1.21 | -2.81 | 4.96E-03 | 3.80E-02 |
| *Gzma* | 6.65 | 3.40 | 0.89 | -3.81 | 1.37E-04 | 2.63E-03 |
| *Actl6b* | 5.49 | 3.39 | 0.88 | -3.86 | 1.12E-04 | 2.26E-03 |
| *Crygf* | 146.93 | 3.37 | 0.86 | -3.90 | 9.57E-05 | 2.01E-03 |
| *Cryba1* | 1200.10 | 3.37 | 0.95 | -3.54 | 4.01E-04 | 6.08E-03 |
| *Gm33347* | 10.02 | 3.36 | 1.13 | -2.98 | 2.90E-03 | 2.60E-02 |
| *Mip* | 29.35 | 3.36 | 0.78 | -4.29 | 1.82E-05 | 5.26E-04 |
| *Cryba2* | 339.87 | 3.35 | 0.95 | -3.52 | 4.28E-04 | 6.40E-03 |
| *Elovl2* | 6.82 | 3.34 | 1.05 | -3.18 | 1.45E-03 | 1.56E-02 |
| *Crygs* | 794.75 | 3.34 | 0.86 | -3.90 | 9.48E-05 | 1.99E-03 |
| *Cd2* | 9.40 | 3.34 | 0.64 | -5.23 | 1.74E-07 | 1.08E-05 |
| *A930036K24Rik* | 8.22 | 3.33 | 0.92 | -3.63 | 2.81E-04 | 4.62E-03 |
| *Lrit1* | 46.12 | 3.33 | 1.07 | -3.11 | 1.90E-03 | 1.91E-02 |
| *Tulp1* | 179.64 | 3.33 | 1.16 | -2.87 | 4.15E-03 | 3.34E-02 |
| *Tex52* | 5.27 | 3.32 | 0.91 | -3.67 | 2.40E-04 | 4.09E-03 |
| *Des* | 116.33 | 3.30 | 1.10 | -3.01 | 2.65E-03 | 2.43E-02 |
| *Pdc* | 210.10 | 3.30 | 1.07 | -3.09 | 2.03E-03 | 2.00E-02 |
| *Slfn8* | 193.23 | 3.29 | 0.36 | -9.26 | 2.08E-20 | 2.29E-17 |
| *Pitpnm2os2* | 11.79 | 3.27 | 1.07 | -3.06 | 2.24E-03 | 2.16E-02 |
| *1110002E22Rik* | 17.03 | 3.27 | 0.86 | -3.81 | 1.40E-04 | 2.67E-03 |
| *Cryaa* | 3983.86 | 3.25 | 0.87 | -3.73 | 1.91E-04 | 3.44E-03 |
| *Gm4759* | 14.58 | 3.24 | 0.76 | -4.27 | 1.98E-05 | 5.61E-04 |

| | | | | | | |
|---|---|---|---|---|---|---|
| *Impg2* | 60.58 | 3.24 | 1.00 | -3.26 | 1.12E-03 | 1.30E-02 |
| *5830417I10Rik* | 26.87 | 3.24 | 1.16 | -2.78 | 5.38E-03 | 4.03E-02 |
| *Pde6b* | 303.06 | 3.24 | 1.06 | -3.05 | 2.27E-03 | 2.18E-02 |
| *Rgs1* | 9.02 | 3.24 | 0.81 | -3.99 | 6.49E-05 | 1.48E-03 |
| *Crybb2* | 1661.39 | 3.23 | 0.90 | -3.59 | 3.26E-04 | 5.20E-03 |
| *Crygb* | 886.87 | 3.22 | 0.88 | -3.65 | 2.63E-04 | 4.39E-03 |
| *Rpe65* | 99.63 | 3.21 | 0.81 | -3.95 | 7.73E-05 | 1.68E-03 |
| *Crb1* | 47.15 | 3.20 | 1.12 | -2.86 | 4.30E-03 | 3.43E-02 |
| *Ttr* | 241.94 | 3.20 | 0.73 | -4.41 | 1.06E-05 | 3.31E-04 |
| *Gm5941* | 47.42 | 3.19 | 0.59 | -5.43 | 5.55E-08 | 4.18E-06 |
| *Sectm1a* | 6.89 | 3.19 | 0.81 | -3.96 | 7.53E-05 | 1.65E-03 |
| *Cd7* | 5.67 | 3.19 | 0.79 | -4.02 | 5.77E-05 | 1.34E-03 |
| *Htr1d* | 5.68 | 3.19 | 0.89 | -3.56 | 3.66E-04 | 5.64E-03 |
| *Cd247* | 7.67 | 3.18 | 0.74 | -4.32 | 1.59E-05 | 4.70E-04 |
| *Neurod4* | 14.84 | 3.17 | 0.90 | -3.54 | 4.03E-04 | 6.09E-03 |
| *Dusp8* | 126.21 | 3.17 | 0.43 | -7.39 | 1.44E-13 | 5.18E-11 |
| *Ifit3* | 808.74 | 3.17 | 0.59 | -5.40 | 6.53E-08 | 4.76E-06 |
| *Pinlyp* | 24.37 | 3.16 | 0.62 | -5.12 | 3.08E-07 | 1.74E-05 |
| *Igtp* | 1088.19 | 3.16 | 0.31 | -10.29 | 7.66E-25 | 1.38E-21 |
| *Gm32443* | 8.56 | 3.15 | 0.95 | -3.31 | 9.30E-04 | 1.13E-02 |
| *Gm47647* | 10.34 | 3.14 | 1.11 | -2.84 | 4.50E-03 | 3.55E-02 |
| *Klra2* | 5.35 | 3.14 | 1.00 | -3.13 | 1.76E-03 | 1.81E-02 |
| *Foxf2* | 13.04 | 3.13 | 0.74 | -4.25 | 2.17E-05 | 6.01E-04 |
| *Olfm3* | 5.22 | 3.13 | 1.01 | -3.09 | 2.02E-03 | 1.99E-02 |
| *Epha8* | 10.08 | 3.13 | 0.84 | -3.74 | 1.86E-04 | 3.36E-03 |
| *Olfr56* | 41.49 | 3.13 | 0.39 | -8.07 | 7.15E-16 | 4.04E-13 |
| *Neurod1* | 47.83 | 3.12 | 1.15 | -2.71 | 6.80E-03 | 4.77E-02 |
| *Six6* | 11.51 | 3.12 | 0.82 | -3.81 | 1.39E-04 | 2.65E-03 |
| *Hhatl* | 16.48 | 3.11 | 0.94 | -3.33 | 8.84E-04 | 1.09E-02 |
| *Slc24a1* | 212.56 | 3.11 | 1.10 | -2.84 | 4.54E-03 | 3.57E-02 |
| *Lhx4* | 3.57 | 3.10 | 1.14 | -2.72 | 6.46E-03 | 4.61E-02 |
| *Ahrr* | 10.31 | 3.09 | 0.94 | -3.28 | 1.05E-03 | 1.24E-02 |
| *Gm45518* | 5.54 | 3.09 | 1.09 | -2.85 | 4.40E-03 | 3.49E-02 |
| *Vgf* | 7.40 | 3.09 | 1.02 | -3.03 | 2.41E-03 | 2.27E-02 |
| *Upb1* | 27.15 | 3.08 | 0.54 | -5.69 | 1.24E-08 | 1.09E-06 |
| *Lpar3* | 3.55 | 3.06 | 1.11 | -2.75 | 5.93E-03 | 4.32E-02 |
| *Calhm6* | 5.66 | 3.06 | 0.98 | -3.13 | 1.73E-03 | 1.78E-02 |
| *Plaur* | 148.28 | 3.05 | 0.73 | -4.17 | 2.99E-05 | 7.76E-04 |
| *Alox15* | 857.91 | 3.05 | 0.76 | -4.02 | 5.75E-05 | 1.33E-03 |

| | | | | | | |
|---|---|---|---|---|---|---|
| *Platr17* | 19.65 | 3.05 | 0.90 | -3.37 | 7.43E-04 | 9.72E-03 |
| *Gbp9* | 401.22 | 3.04 | 0.35 | -8.65 | 5.22E-18 | 3.97E-15 |
| *Ctla4* | 16.07 | 3.03 | 0.65 | -4.69 | 2.79E-06 | 1.10E-04 |
| *B2m* | 9727.45 | 3.02 | 0.22 | -13.57 | 5.95E-42 | 2.35E-38 |
| *Slfn1* | 54.86 | 3.02 | 0.42 | -7.20 | 6.00E-13 | 1.80E-10 |
| *Sh3bgr* | 17.55 | 3.01 | 0.86 | -3.51 | 4.51E-04 | 6.65E-03 |
| *Ccl7* | 6.61 | 3.01 | 0.96 | -3.15 | 1.63E-03 | 1.70E-02 |
| *Gimap3* | 15.63 | 2.99 | 0.52 | -5.73 | 1.00E-08 | 9.04E-07 |
| *Cfap73* | 3.22 | 2.99 | 1.02 | -2.93 | 3.43E-03 | 2.92E-02 |
| *Vash1* | 8.62 | 2.98 | 1.09 | -2.75 | 6.04E-03 | 4.38E-02 |
| *Zc3h12d* | 5.32 | 2.98 | 0.87 | -3.42 | 6.21E-04 | 8.53E-03 |
| *Tshr* | 6.80 | 2.98 | 0.91 | -3.28 | 1.04E-03 | 1.23E-02 |
| *Cxcr3* | 6.89 | 2.97 | 0.85 | -3.51 | 4.50E-04 | 6.64E-03 |
| *Pdcd1* | 19.53 | 2.97 | 0.59 | -5.04 | 4.60E-07 | 2.44E-05 |
| *AI847159* | 13.77 | 2.97 | 0.86 | -3.43 | 6.00E-04 | 8.31E-03 |
| *Hspb7* | 21.09 | 2.97 | 0.89 | -3.34 | 8.46E-04 | 1.06E-02 |
| *Ptger2* | 3.43 | 2.96 | 1.01 | -2.92 | 3.47E-03 | 2.95E-02 |
| *Tlr12* | 4.05 | 2.95 | 1.08 | -2.73 | 6.39E-03 | 4.57E-02 |
| *Klrd1* | 4.08 | 2.93 | 0.90 | -3.25 | 1.14E-03 | 1.31E-02 |
| *Sntg2* | 5.35 | 2.93 | 0.89 | -3.28 | 1.02E-03 | 1.21E-02 |
| *Fosb* | 725.29 | 2.93 | 0.37 | -7.99 | 1.39E-15 | 7.07E-13 |
| *Tdrd9* | 4.88 | 2.92 | 1.06 | -2.74 | 6.13E-03 | 4.43E-02 |
| *Gm6611* | 7.65 | 2.91 | 0.79 | -3.67 | 2.41E-04 | 4.10E-03 |
| *Irgm2* | 831.46 | 2.91 | 0.33 | -8.82 | 1.13E-18 | 1.02E-15 |
| *Sag* | 1020.90 | 2.90 | 1.04 | -2.79 | 5.32E-03 | 4.00E-02 |
| *Il2rb* | 28.87 | 2.89 | 0.63 | -4.61 | 3.98E-06 | 1.47E-04 |
| *Inhba* | 7.94 | 2.88 | 0.71 | -4.07 | 4.77E-05 | 1.14E-03 |
| *Gpr3* | 10.10 | 2.87 | 0.59 | -4.84 | 1.30E-06 | 5.83E-05 |
| *Sh2d5* | 417.27 | 2.87 | 0.56 | -5.16 | 2.52E-07 | 1.49E-05 |
| *Slco1c1* | 6.70 | 2.86 | 0.96 | -2.97 | 2.97E-03 | 2.64E-02 |
| *Slc26a10* | 9.66 | 2.86 | 0.63 | -4.50 | 6.69E-06 | 2.24E-04 |
| *Apol9b* | 1174.88 | 2.84 | 0.28 | -10.06 | 8.10E-24 | 1.34E-20 |
| *Cabp5* | 19.06 | 2.84 | 0.94 | -3.01 | 2.60E-03 | 2.40E-02 |
| *Vsx2* | 38.25 | 2.84 | 0.69 | -4.08 | 4.46E-05 | 1.08E-03 |
| *Serpinb2* | 72.76 | 2.84 | 0.83 | -3.42 | 6.34E-04 | 8.62E-03 |
| *Itgal* | 32.27 | 2.82 | 0.37 | -7.69 | 1.48E-14 | 6.37E-12 |
| *Cryga* | 20.54 | 2.82 | 0.62 | -4.56 | 5.06E-06 | 1.77E-04 |
| *Kcnj12* | 7.58 | 2.81 | 0.99 | -2.82 | 4.73E-03 | 3.67E-02 |
| *Gm4070* | 22.62 | 2.80 | 0.43 | -6.56 | 5.26E-11 | 9.38E-09 |

| | | | | | | |
|---|---|---|---|---|---|---|
| *Kcnj8* | 4.34 | 2.80 | 0.95 | -2.96 | 3.11E-03 | 2.73E-02 |
| *Gdf15* | 9.38 | 2.80 | 0.86 | -3.27 | 1.09E-03 | 1.27E-02 |
| *Casp12* | 59.75 | 2.80 | 0.32 | -8.66 | 4.69E-18 | 3.71E-15 |
| *Gm49485* | 6.15 | 2.80 | 0.96 | -2.91 | 3.59E-03 | 3.03E-02 |
| *Bhlhe22* | 25.58 | 2.80 | 0.62 | -4.52 | 6.06E-06 | 2.07E-04 |
| *Ccl8* | 19.60 | 2.79 | 0.49 | -5.72 | 1.05E-08 | 9.45E-07 |
| *Gpr152* | 17.01 | 2.79 | 0.95 | -2.93 | 3.42E-03 | 2.92E-02 |
| *Birc7* | 5.79 | 2.78 | 1.02 | -2.72 | 6.51E-03 | 4.63E-02 |
| *Nxnl2* | 14.08 | 2.78 | 1.03 | -2.71 | 6.76E-03 | 4.76E-02 |
| *Itk* | 13.35 | 2.78 | 0.70 | -3.96 | 7.54E-05 | 1.65E-03 |
| *Foxp3* | 8.80 | 2.78 | 0.66 | -4.19 | 2.79E-05 | 7.34E-04 |
| *Pax8* | 6.46 | 2.78 | 0.99 | -2.79 | 5.24E-03 | 3.96E-02 |
| *Myo18b* | 14.26 | 2.78 | 0.91 | -3.06 | 2.18E-03 | 2.11E-02 |
| *Trbc2* | 14.92 | 2.76 | 0.50 | -5.50 | 3.87E-08 | 3.05E-06 |
| *Zbp1* | 873.14 | 2.76 | 0.30 | -9.23 | 2.76E-20 | 2.88E-17 |
| *Crybb3* | 471.82 | 2.75 | 0.54 | -5.06 | 4.27E-07 | 2.29E-05 |
| *Svop* | 15.12 | 2.75 | 0.77 | -3.56 | 3.68E-04 | 5.66E-03 |
| *Selp* | 10.47 | 2.75 | 0.89 | -3.07 | 2.13E-03 | 2.07E-02 |
| *Cd3g* | 10.36 | 2.73 | 0.67 | -4.10 | 4.18E-05 | 1.02E-03 |
| *Rab37* | 5.22 | 2.72 | 0.93 | -2.93 | 3.37E-03 | 2.89E-02 |
| *Clec12a* | 7.32 | 2.72 | 0.75 | -3.64 | 2.69E-04 | 4.46E-03 |
| *Ifi44* | 1544.36 | 2.71 | 0.66 | -4.13 | 3.65E-05 | 9.15E-04 |
| *Zdhhc22* | 8.37 | 2.71 | 0.81 | -3.32 | 8.98E-04 | 1.10E-02 |
| *Cnn1* | 7.55 | 2.71 | 0.95 | -2.84 | 4.51E-03 | 3.55E-02 |
| *Rgs7* | 6.98 | 2.70 | 0.88 | -3.09 | 2.02E-03 | 1.99E-02 |
| *Ccl22* | 12.12 | 2.70 | 0.71 | -3.81 | 1.40E-04 | 2.67E-03 |
| *Igsf11* | 12.35 | 2.70 | 0.79 | -3.41 | 6.54E-04 | 8.83E-03 |
| *AC164883.2* | 11.24 | 2.69 | 0.62 | -4.34 | 1.43E-05 | 4.29E-04 |
| *Slc29a4* | 7.17 | 2.68 | 0.77 | -3.49 | 4.77E-04 | 6.91E-03 |
| *Chrm4* | 5.03 | 2.68 | 0.93 | -2.88 | 3.95E-03 | 3.24E-02 |
| *Phf11a* | 23.10 | 2.68 | 0.55 | -4.85 | 1.21E-06 | 5.52E-05 |
| *Nr4a1* | 269.72 | 2.68 | 0.38 | -6.97 | 3.06E-12 | 7.76E-10 |
| *5430430K15Rik* | 5.50 | 2.67 | 0.78 | -3.45 | 5.69E-04 | 7.96E-03 |
| *Gm26954* | 8.06 | 2.67 | 0.96 | -2.77 | 5.60E-03 | 4.15E-02 |
| *Gm16310* | 21.38 | 2.66 | 0.77 | -3.46 | 5.44E-04 | 7.70E-03 |
| *Il2ra* | 18.51 | 2.66 | 0.55 | -4.83 | 1.39E-06 | 6.12E-05 |
| *Pex5l* | 63.85 | 2.66 | 0.83 | -3.19 | 1.43E-03 | 1.55E-02 |
| *Rlbp1* | 124.42 | 2.65 | 0.73 | -3.64 | 2.76E-04 | 4.56E-03 |
| *Lhx6* | 8.05 | 2.65 | 0.71 | -3.71 | 2.06E-04 | 3.64E-03 |

| | | | | | | |
|---|---|---|---|---|---|---|
| *Cdk5r2* | 42.77 | 2.64 | 0.91 | -2.91 | 3.60E-03 | 3.03E-02 |
| *Chga* | 26.17 | 2.64 | 0.51 | -5.16 | 2.50E-07 | 1.48E-05 |
| *Ccr7* | 10.32 | 2.63 | 0.59 | -4.44 | 8.80E-06 | 2.84E-04 |
| *Ms4a4b* | 11.11 | 2.63 | 0.63 | -4.16 | 3.21E-05 | 8.22E-04 |
| *B4galnt3* | 9.90 | 2.62 | 0.67 | -3.91 | 9.41E-05 | 1.98E-03 |
| *Sprr2g* | 113.95 | 2.62 | 0.88 | -2.97 | 2.94E-03 | 2.62E-02 |
| *Cabp4* | 34.21 | 2.62 | 0.75 | -3.48 | 4.96E-04 | 7.12E-03 |
| *C430002N11Rik* | 18.12 | 2.62 | 0.72 | -3.63 | 2.85E-04 | 4.67E-03 |
| *Gm14275* | 9.81 | 2.62 | 0.83 | -3.15 | 1.64E-03 | 1.71E-02 |
| *Sh2d2a* | 9.27 | 2.61 | 0.61 | -4.28 | 1.83E-05 | 5.26E-04 |
| *Kcnq1* | 5.56 | 2.61 | 0.82 | -3.18 | 1.47E-03 | 1.58E-02 |
| *Rbm24* | 20.50 | 2.60 | 0.88 | -2.94 | 3.24E-03 | 2.80E-02 |
| *H2-DMb1* | 27.44 | 2.59 | 0.58 | -4.48 | 7.37E-06 | 2.43E-04 |
| *Scimp* | 6.17 | 2.59 | 0.87 | -2.96 | 3.04E-03 | 2.69E-02 |
| *Grifin* | 36.16 | 2.58 | 0.43 | -5.98 | 2.26E-09 | 2.45E-07 |
| *Lrat* | 45.94 | 2.57 | 0.86 | -2.99 | 2.79E-03 | 2.52E-02 |
| *Gm15433* | 103.80 | 2.57 | 0.76 | -3.37 | 7.64E-04 | 9.89E-03 |
| *Ccr5* | 27.51 | 2.57 | 0.46 | -5.63 | 1.82E-08 | 1.55E-06 |
| *Gm7582* | 34.35 | 2.57 | 0.53 | -4.89 | 1.01E-06 | 4.74E-05 |
| *Alox12e* | 1746.58 | 2.56 | 0.89 | -2.86 | 4.17E-03 | 3.36E-02 |
| *Mmp12* | 37.00 | 2.56 | 0.51 | -4.98 | 6.37E-07 | 3.15E-05 |
| *Slamf7* | 5.34 | 2.56 | 0.85 | -3.01 | 2.58E-03 | 2.38E-02 |
| *Il27ra* | 5.42 | 2.55 | 0.76 | -3.35 | 8.16E-04 | 1.03E-02 |
| *Slc39a12* | 13.60 | 2.55 | 0.88 | -2.90 | 3.71E-03 | 3.09E-02 |
| *Ms4a4c* | 14.91 | 2.53 | 0.53 | -4.78 | 1.77E-06 | 7.60E-05 |
| *Lingo3* | 15.18 | 2.52 | 0.76 | -3.31 | 9.45E-04 | 1.15E-02 |
| *Dmkn* | 63.20 | 2.52 | 0.60 | -4.22 | 2.41E-05 | 6.48E-04 |
| *Slc17a6* | 15.12 | 2.51 | 0.59 | -4.28 | 1.83E-05 | 5.26E-04 |
| *Slco1a4* | 13.38 | 2.51 | 0.66 | -3.79 | 1.52E-04 | 2.84E-03 |
| *Hist1h1b* | 16.07 | 2.51 | 0.93 | -2.70 | 6.95E-03 | 4.85E-02 |
| *Gm20628* | 12.10 | 2.50 | 0.85 | -2.96 | 3.10E-03 | 2.73E-02 |
| *Gm18445* | 91.53 | 2.50 | 0.53 | -4.74 | 2.18E-06 | 8.97E-05 |
| *Gnat2* | 20.99 | 2.50 | 0.69 | -3.61 | 3.01E-04 | 4.89E-03 |
| *Gdnf* | 52.58 | 2.49 | 0.33 | -7.46 | 8.88E-14 | 3.45E-11 |
| *Frrs1l* | 9.62 | 2.49 | 0.87 | -2.85 | 4.42E-03 | 3.50E-02 |
| *C230004F18Rik* | 5.03 | 2.48 | 0.90 | -2.75 | 5.90E-03 | 4.31E-02 |
| *Tnfrsf23* | 77.35 | 2.48 | 0.48 | -5.13 | 2.85E-07 | 1.65E-05 |
| *Chrna2* | 6.15 | 2.47 | 0.73 | -3.41 | 6.61E-04 | 8.90E-03 |
| *Gm12185* | 12.89 | 2.47 | 0.65 | -3.78 | 1.56E-04 | 2.91E-03 |

| | | | | | | |
|---|---|---|---|---|---|---|
| *Mafa* | 7.14 | 2.46 | 0.87 | -2.84 | 4.48E-03 | 3.54E-02 |
| *Itgae* | 13.85 | 2.46 | 0.60 | -4.08 | 4.55E-05 | 1.10E-03 |
| *Scg3* | 19.85 | 2.45 | 0.72 | -3.39 | 6.89E-04 | 9.17E-03 |
| *Sox2* | 6.46 | 2.45 | 0.76 | -3.21 | 1.34E-03 | 1.48E-02 |
| *Atp1a3* | 420.46 | 2.45 | 0.76 | -3.22 | 1.29E-03 | 1.44E-02 |
| *Cryba4* | 555.07 | 2.44 | 0.39 | -6.32 | 2.56E-10 | 3.83E-08 |
| *Dusp26* | 11.98 | 2.43 | 0.66 | -3.69 | 2.22E-04 | 3.84E-03 |
| *Fxyd2* | 5.94 | 2.42 | 0.81 | -3.00 | 2.69E-03 | 2.46E-02 |
| *Syt1* | 136.97 | 2.42 | 0.70 | -3.47 | 5.20E-04 | 7.41E-03 |
| *Synpo2l* | 16.47 | 2.41 | 0.79 | -3.07 | 2.15E-03 | 2.09E-02 |
| *Nptx2* | 10.44 | 2.40 | 0.65 | -3.68 | 2.36E-04 | 4.04E-03 |
| *Ciita* | 45.01 | 2.40 | 0.52 | -4.61 | 4.07E-06 | 1.48E-04 |
| *Mab21l2* | 17.28 | 2.40 | 0.70 | -3.45 | 5.61E-04 | 7.87E-03 |
| *Oas2* | 2255.74 | 2.39 | 0.35 | -6.92 | 4.66E-12 | 1.10E-09 |
| *Capn11* | 4.64 | 2.39 | 0.86 | -2.77 | 5.60E-03 | 4.15E-02 |
| *Clec10a* | 8.57 | 2.39 | 0.76 | -3.15 | 1.63E-03 | 1.71E-02 |
| *Cd36* | 40.28 | 2.39 | 0.70 | -3.39 | 7.08E-04 | 9.35E-03 |
| *Ptprz1* | 83.80 | 2.39 | 0.39 | -6.04 | 1.50E-09 | 1.74E-07 |
| *Srrm3* | 19.96 | 2.39 | 0.74 | -3.23 | 1.25E-03 | 1.41E-02 |
| *Ppp1r1a* | 18.32 | 2.38 | 0.60 | -3.97 | 7.32E-05 | 1.62E-03 |
| *Gimap4* | 29.47 | 2.37 | 0.41 | -5.75 | 8.75E-09 | 8.10E-07 |
| *H2-DMb2* | 45.15 | 2.37 | 0.48 | -4.93 | 8.31E-07 | 3.99E-05 |
| *Rasd2* | 15.05 | 2.36 | 0.65 | -3.66 | 2.50E-04 | 4.23E-03 |
| *Ap3b2* | 26.92 | 2.36 | 0.56 | -4.21 | 2.55E-05 | 6.82E-04 |
| *Cplx3* | 39.52 | 2.36 | 0.70 | -3.34 | 8.29E-04 | 1.04E-02 |
| *Apol9a* | 1019.82 | 2.35 | 0.27 | -8.77 | 1.82E-18 | 1.56E-15 |
| *Zfp385b* | 15.16 | 2.34 | 0.79 | -2.97 | 2.99E-03 | 2.65E-02 |
| *Slc28a2* | 5.69 | 2.33 | 0.76 | -3.07 | 2.13E-03 | 2.08E-02 |
| *Ighg2b* | 13.12 | 2.33 | 0.60 | -3.90 | 9.66E-05 | 2.02E-03 |
| *Sidt1* | 13.22 | 2.32 | 0.59 | -3.96 | 7.65E-05 | 1.67E-03 |
| *Rd3l* | 14.64 | 2.32 | 0.82 | -2.84 | 4.54E-03 | 3.56E-02 |
| *Irgm1* | 2384.03 | 2.32 | 0.23 | -9.96 | 2.20E-23 | 3.11E-20 |
| *Sgpp2* | 5.80 | 2.32 | 0.83 | -2.80 | 5.17E-03 | 3.92E-02 |
| *P2ry10* | 16.48 | 2.31 | 0.58 | -4.02 | 5.89E-05 | 1.35E-03 |
| *Tmod1* | 22.24 | 2.31 | 0.68 | -3.42 | 6.28E-04 | 8.56E-03 |
| *Bfsp2* | 74.66 | 2.31 | 0.55 | -4.20 | 2.66E-05 | 7.07E-04 |
| *Pcdhac2* | 5.72 | 2.31 | 0.81 | -2.84 | 4.48E-03 | 3.54E-02 |
| *Ctsw* | 10.45 | 2.30 | 0.64 | -3.63 | 2.88E-04 | 4.71E-03 |
| *Ifit1* | 1885.98 | 2.29 | 0.33 | -7.04 | 1.94E-12 | 5.33E-10 |

| | | | | | | |
|---|---|---|---|---|---|---|
| *Rgs8* | 24.23 | 2.29 | 0.60 | -3.80 | 1.46E-04 | 2.75E-03 |
| *Adm* | 155.61 | 2.29 | 0.46 | -4.98 | 6.37E-07 | 3.15E-05 |
| *Pou3f1* | 11.11 | 2.29 | 0.76 | -3.00 | 2.74E-03 | 2.49E-02 |
| *Kcnq3* | 8.88 | 2.28 | 0.71 | -3.24 | 1.20E-03 | 1.36E-02 |
| *Sh2d4b* | 16.78 | 2.27 | 0.57 | -4.01 | 6.11E-05 | 1.40E-03 |
| *Otx2* | 63.15 | 2.27 | 0.84 | -2.72 | 6.59E-03 | 4.68E-02 |
| *Rorb* | 22.82 | 2.27 | 0.71 | -3.19 | 1.43E-03 | 1.55E-02 |
| *Insl6* | 18.04 | 2.26 | 0.45 | -5.03 | 4.99E-07 | 2.60E-05 |
| *Paqr9* | 12.66 | 2.25 | 0.77 | -2.91 | 3.58E-03 | 3.02E-02 |
| *Adh6a* | 31.97 | 2.25 | 0.69 | -3.28 | 1.05E-03 | 1.24E-02 |
| *Lyz1* | 12.29 | 2.25 | 0.61 | -3.71 | 2.11E-04 | 3.70E-03 |
| *Gm8251* | 9.75 | 2.24 | 0.82 | -2.74 | 6.20E-03 | 4.48E-02 |
| *Phyhipl* | 105.64 | 2.24 | 0.49 | -4.56 | 5.23E-06 | 1.82E-04 |
| *Brsk2* | 29.78 | 2.23 | 0.67 | -3.35 | 8.15E-04 | 1.03E-02 |
| *Arc* | 247.71 | 2.23 | 0.32 | -6.95 | 3.71E-12 | 8.96E-10 |
| *Rax* | 18.19 | 2.22 | 0.64 | -3.49 | 4.82E-04 | 6.97E-03 |
| *Gm16464* | 13.07 | 2.22 | 0.58 | -3.85 | 1.18E-04 | 2.34E-03 |
| *Faxc* | 14.85 | 2.21 | 0.66 | -3.37 | 7.58E-04 | 9.84E-03 |
| *Pcdh10* | 18.87 | 2.21 | 0.60 | -3.65 | 2.58E-04 | 4.33E-03 |
| *Trim67* | 8.62 | 2.20 | 0.75 | -2.92 | 3.53E-03 | 2.99E-02 |
| *Gabrg2* | 14.05 | 2.20 | 0.66 | -3.34 | 8.30E-04 | 1.04E-02 |
| *Sez6* | 16.37 | 2.20 | 0.73 | -3.02 | 2.52E-03 | 2.34E-02 |
| *Phkg1* | 24.17 | 2.19 | 0.79 | -2.76 | 5.75E-03 | 4.23E-02 |
| *Aim2* | 15.82 | 2.19 | 0.51 | -4.26 | 2.08E-05 | 5.82E-04 |
| *H2-M2* | 8.94 | 2.18 | 0.66 | -3.33 | 8.65E-04 | 1.07E-02 |
| *Gm13822* | 5.77 | 2.18 | 0.76 | -2.89 | 3.86E-03 | 3.18E-02 |
| *Jun* | 4194.44 | 2.18 | 0.19 | -11.30 | 1.28E-29 | 2.81E-26 |
| *Rtp4* | 1371.16 | 2.18 | 0.25 | -8.87 | 7.40E-19 | 6.98E-16 |
| *Dusp10* | 309.84 | 2.18 | 0.37 | -5.90 | 3.70E-09 | 3.76E-07 |
| *Lgals3bp* | 1158.99 | 2.18 | 0.23 | -9.38 | 6.87E-21 | 8.00E-18 |
| *Sp110* | 269.20 | 2.18 | 0.22 | -10.05 | 8.79E-24 | 1.34E-20 |
| *Il21r* | 13.41 | 2.17 | 0.54 | -4.04 | 5.36E-05 | 1.26E-03 |
| *Slc17a8* | 6.00 | 2.17 | 0.78 | -2.77 | 5.56E-03 | 4.13E-02 |
| *Rasd1* | 23.15 | 2.17 | 0.57 | -3.81 | 1.41E-04 | 2.68E-03 |
| *Rom1* | 349.20 | 2.16 | 0.65 | -3.35 | 8.05E-04 | 1.02E-02 |
| *Egr3* | 8.97 | 2.16 | 0.76 | -2.86 | 4.26E-03 | 3.40E-02 |
| *H2-Ab1* | 215.29 | 2.16 | 0.31 | -7.07 | 1.55E-12 | 4.45E-10 |
| *Ppp2r2c* | 31.87 | 2.16 | 0.47 | -4.57 | 4.92E-06 | 1.73E-04 |
| *Rdh12* | 93.55 | 2.15 | 0.78 | -2.76 | 5.83E-03 | 4.27E-02 |

| | | | | | | |
|---|---|---|---|---|---|---|
| *Fabp3* | 44.29 | 2.15 | 0.60 | -3.59 | 3.28E-04 | 5.21E-03 |
| *Jph2* | 75.29 | 2.15 | 0.45 | -4.73 | 2.26E-06 | 9.24E-05 |
| *Rnase6* | 7.72 | 2.14 | 0.67 | -3.19 | 1.41E-03 | 1.53E-02 |
| *Gabrb3* | 22.45 | 2.14 | 0.56 | -3.80 | 1.47E-04 | 2.76E-03 |
| *Syt4* | 47.24 | 2.14 | 0.51 | -4.19 | 2.79E-05 | 7.34E-04 |
| *Cd74* | 1921.85 | 2.14 | 0.29 | -7.41 | 1.26E-13 | 4.72E-11 |
| *Gm7785* | 5.96 | 2.14 | 0.69 | -3.09 | 1.98E-03 | 1.96E-02 |
| *Usp18* | 546.19 | 2.14 | 0.30 | -7.12 | 1.11E-12 | 3.29E-10 |
| *Gm7897* | 937.45 | 2.14 | 0.32 | -6.75 | 1.50E-11 | 3.09E-09 |
| *Nell2* | 24.46 | 2.13 | 0.44 | -4.85 | 1.24E-06 | 5.60E-05 |
| *H2-Eb1* | 569.40 | 2.13 | 0.29 | -7.39 | 1.48E-13 | 5.21E-11 |
| *Trbc1* | 7.21 | 2.13 | 0.70 | -3.03 | 2.43E-03 | 2.29E-02 |
| *C4b* | 322.73 | 2.13 | 0.32 | -6.60 | 4.05E-11 | 7.72E-09 |
| *C1ql1* | 18.78 | 2.12 | 0.65 | -3.24 | 1.19E-03 | 1.35E-02 |
| *Dusp4* | 58.90 | 2.11 | 0.47 | -4.52 | 6.05E-06 | 2.07E-04 |
| *Slfn4* | 919.42 | 2.11 | 0.48 | -4.44 | 9.04E-06 | 2.90E-04 |
| *Gbp8* | 119.35 | 2.10 | 0.33 | -6.32 | 2.63E-10 | 3.88E-08 |
| *Ddx25* | 7.81 | 2.10 | 0.74 | -2.83 | 4.70E-03 | 3.65E-02 |
| *Mustn1* | 13.56 | 2.09 | 0.63 | -3.32 | 9.08E-04 | 1.11E-02 |
| *Unc80* | 26.59 | 2.09 | 0.64 | -3.25 | 1.15E-03 | 1.33E-02 |
| *Camk2a* | 17.32 | 2.08 | 0.77 | -2.71 | 6.73E-03 | 4.75E-02 |
| *Gm16025* | 55.11 | 2.08 | 0.33 | -6.32 | 2.58E-10 | 3.83E-08 |
| *Syp* | 94.29 | 2.08 | 0.72 | -2.90 | 3.73E-03 | 3.11E-02 |
| *Spink2* | 15.71 | 2.08 | 0.56 | -3.75 | 1.78E-04 | 3.25E-03 |
| *Camkv* | 26.24 | 2.08 | 0.52 | -3.98 | 7.03E-05 | 1.57E-03 |
| *Pygm* | 254.74 | 2.07 | 0.75 | -2.75 | 5.96E-03 | 4.34E-02 |
| *Camk2b* | 42.96 | 2.07 | 0.54 | -3.80 | 1.46E-04 | 2.75E-03 |
| *Cdh22* | 6.95 | 2.07 | 0.76 | -2.73 | 6.40E-03 | 4.58E-02 |
| *Lck* | 22.33 | 2.06 | 0.39 | -5.26 | 1.41E-07 | 8.95E-06 |
| *Rasal3* | 22.23 | 2.06 | 0.44 | -4.64 | 3.41E-06 | 1.30E-04 |
| *Cytip* | 34.64 | 2.06 | 0.35 | -5.90 | 3.63E-09 | 3.70E-07 |
| *Igfbp3* | 124.63 | 2.05 | 0.64 | -3.19 | 1.41E-03 | 1.53E-02 |
| *Ifi204* | 158.61 | 2.05 | 0.25 | -8.05 | 8.59E-16 | 4.73E-13 |
| *Fgl2* | 103.97 | 2.05 | 0.34 | -6.05 | 1.47E-09 | 1.72E-07 |
| *Dnase1l3* | 48.39 | 2.05 | 0.45 | -4.51 | 6.51E-06 | 2.19E-04 |
| *Col9a1* | 17.73 | 2.05 | 0.72 | -2.83 | 4.61E-03 | 3.60E-02 |
| *Hk3* | 12.77 | 2.05 | 0.63 | -3.24 | 1.21E-03 | 1.37E-02 |
| *Ivl* | 454.04 | 2.05 | 0.40 | -5.09 | 3.51E-07 | 1.93E-05 |
| *S100b* | 43.34 | 2.04 | 0.54 | -3.75 | 1.73E-04 | 3.18E-03 |

| | | | | | | |
|---|---|---|---|---|---|---|
| *Bst2* | 549.65 | 2.04 | 0.31 | -6.60 | 4.10E-11 | 7.73E-09 |
| *Samsn1* | 10.81 | 2.04 | 0.50 | -4.03 | 5.50E-05 | 1.29E-03 |
| *Ly6d* | 63.13 | 2.03 | 0.46 | -4.45 | 8.42E-06 | 2.72E-04 |
| *Sox10* | 29.67 | 2.03 | 0.47 | -4.30 | 1.68E-05 | 4.92E-04 |
| *Ccdc85a* | 8.54 | 2.03 | 0.69 | -2.93 | 3.34E-03 | 2.87E-02 |
| *Gpr65* | 10.58 | 2.02 | 0.70 | -2.87 | 4.10E-03 | 3.31E-02 |
| *Ank1* | 26.55 | 2.02 | 0.61 | -3.32 | 9.14E-04 | 1.11E-02 |
| *Cadm3* | 66.82 | 2.01 | 0.52 | -3.86 | 1.14E-04 | 2.28E-03 |
| *Pax9* | 8.92 | 2.01 | 0.74 | -2.71 | 6.75E-03 | 4.76E-02 |
| *Rgs16* | 45.97 | 2.01 | 0.48 | -4.16 | 3.15E-05 | 8.09E-04 |
| *Vstm2l* | 12.46 | 2.01 | 0.73 | -2.74 | 6.13E-03 | 4.44E-02 |
| *Ptges* | 1274.66 | 2.00 | 0.29 | -7.02 | 2.19E-12 | 5.78E-10 |
| *Kcnj13* | 20.98 | 2.00 | 0.68 | -2.93 | 3.41E-03 | 2.92E-02 |
| *Trpc1* | 59.84 | 2.00 | 0.42 | -4.73 | 2.24E-06 | 9.18E-05 |
| *Kcnma1* | 30.69 | 2.00 | 0.67 | -2.98 | 2.88E-03 | 2.59E-02 |
| *Oas1e* | 40.28 | 2.00 | 0.36 | -5.59 | 2.33E-08 | 1.94E-06 |
| *Prkcq* | 67.54 | 2.00 | 0.35 | -5.78 | 7.52E-09 | 7.02E-07 |
| *Il7r* | 19.10 | 1.99 | 0.50 | -3.99 | 6.72E-05 | 1.52E-03 |
| *Insrr* | 13.33 | 1.99 | 0.71 | -2.82 | 4.73E-03 | 3.67E-02 |
| *Pla2g4e* | 27.95 | 1.99 | 0.54 | -3.65 | 2.61E-04 | 4.37E-03 |
| *Tmem72* | 17.18 | 1.99 | 0.71 | -2.78 | 5.41E-03 | 4.05E-02 |
| *Gm15728* | 20.16 | 1.98 | 0.43 | -4.60 | 4.23E-06 | 1.52E-04 |
| *Slc11a1* | 18.69 | 1.97 | 0.38 | -5.21 | 1.94E-07 | 1.18E-05 |
| *Csprs* | 70.68 | 1.97 | 0.34 | -5.86 | 4.54E-09 | 4.45E-07 |
| *Guca1b* | 175.53 | 1.96 | 0.65 | -3.01 | 2.62E-03 | 2.41E-02 |
| *Ptprn* | 37.24 | 1.96 | 0.50 | -3.90 | 9.73E-05 | 2.03E-03 |
| *Pcsk1n* | 49.76 | 1.96 | 0.60 | -3.28 | 1.06E-03 | 1.24E-02 |
| *Oas3* | 2065.92 | 1.96 | 0.25 | -7.73 | 1.04E-14 | 4.70E-12 |
| *Thsd7a* | 20.94 | 1.96 | 0.58 | -3.36 | 7.71E-04 | 9.93E-03 |
| *Trpv2* | 17.88 | 1.96 | 0.55 | -3.55 | 3.82E-04 | 5.85E-03 |
| *Selplg* | 51.79 | 1.95 | 0.44 | -4.43 | 9.25E-06 | 2.95E-04 |
| *Lce3c* | 30.08 | 1.95 | 0.68 | -2.88 | 4.01E-03 | 3.26E-02 |
| *9930111J21Rik1* | 37.74 | 1.95 | 0.54 | -3.61 | 3.09E-04 | 4.96E-03 |
| *Sbsn* | 153.69 | 1.95 | 0.41 | -4.76 | 1.91E-06 | 8.04E-05 |
| *Lgals9* | 862.67 | 1.94 | 0.23 | -8.60 | 7.81E-18 | 5.72E-15 |
| *Nanos1* | 14.47 | 1.94 | 0.59 | -3.29 | 1.02E-03 | 1.21E-02 |
| *Crabp2* | 60.47 | 1.94 | 0.45 | -4.27 | 1.95E-05 | 5.56E-04 |
| *Clec4a3* | 9.02 | 1.93 | 0.64 | -3.02 | 2.56E-03 | 2.37E-02 |
| *Wdfy4* | 28.22 | 1.93 | 0.37 | -5.27 | 1.35E-07 | 8.71E-06 |

| | | | | | | |
|---|---|---|---|---|---|---|
| *Gucy1a1* | 38.45 | 1.93 | 0.44 | -4.44 | 9.02E-06 | 2.90E-04 |
| *Tbx1* | 19.49 | 1.93 | 0.60 | -3.20 | 1.38E-03 | 1.51E-02 |
| *Nkain1* | 7.32 | 1.93 | 0.66 | -2.91 | 3.57E-03 | 3.01E-02 |
| *Phf11b* | 88.73 | 1.93 | 0.31 | -6.14 | 8.04E-10 | 1.03E-07 |
| *Bst1* | 15.21 | 1.93 | 0.57 | -3.41 | 6.58E-04 | 8.88E-03 |
| *Gdap1l1* | 8.44 | 1.93 | 0.69 | -2.79 | 5.23E-03 | 3.96E-02 |
| *Mpeg1* | 232.53 | 1.92 | 0.30 | -6.35 | 2.22E-10 | 3.41E-08 |
| *Gpr37* | 21.46 | 1.92 | 0.57 | -3.39 | 6.89E-04 | 9.17E-03 |
| *Krt87* | 11.30 | 1.92 | 0.54 | -3.59 | 3.27E-04 | 5.21E-03 |
| *Clec4a2* | 8.87 | 1.92 | 0.64 | -3.00 | 2.72E-03 | 2.47E-02 |
| *Slc6a11* | 30.69 | 1.92 | 0.57 | -3.34 | 8.44E-04 | 1.05E-02 |
| *Pdlim3* | 19.51 | 1.92 | 0.68 | -2.82 | 4.74E-03 | 3.67E-02 |
| *Gm44419* | 12.59 | 1.91 | 0.71 | -2.69 | 7.17E-03 | 4.95E-02 |
| *Trac* | 12.28 | 1.91 | 0.52 | -3.70 | 2.12E-04 | 3.71E-03 |
| *Sp140* | 82.79 | 1.91 | 0.27 | -6.96 | 3.47E-12 | 8.48E-10 |
| *Chgb* | 48.23 | 1.91 | 0.63 | -3.01 | 2.60E-03 | 2.40E-02 |
| *Isg15* | 1974.25 | 1.90 | 0.28 | -6.91 | 4.95E-12 | 1.14E-09 |
| *Glipr1* | 12.67 | 1.90 | 0.59 | -3.20 | 1.36E-03 | 1.50E-02 |
| *B4galnt4* | 84.97 | 1.90 | 0.40 | -4.75 | 2.06E-06 | 8.50E-05 |
| *Trim47* | 55.99 | 1.90 | 0.43 | -4.38 | 1.16E-05 | 3.61E-04 |
| *Prox1* | 27.37 | 1.90 | 0.57 | -3.33 | 8.78E-04 | 1.08E-02 |
| *Gipc2* | 75.73 | 1.89 | 0.29 | -6.54 | 6.02E-11 | 1.04E-08 |
| *Cd3e* | 11.61 | 1.89 | 0.60 | -3.14 | 1.69E-03 | 1.75E-02 |
| *Dusp1* | 4130.98 | 1.89 | 0.30 | -6.19 | 6.16E-10 | 8.08E-08 |
| *Scg2* | 80.07 | 1.89 | 0.46 | -4.12 | 3.71E-05 | 9.25E-04 |
| *Cldn5* | 27.91 | 1.88 | 0.42 | -4.48 | 7.61E-06 | 2.50E-04 |
| *Hspa4l* | 887.14 | 1.88 | 0.51 | -3.68 | 2.32E-04 | 3.98E-03 |
| *Lrrc25* | 15.88 | 1.88 | 0.53 | -3.56 | 3.74E-04 | 5.74E-03 |
| *Prss16* | 21.85 | 1.88 | 0.51 | -3.70 | 2.16E-04 | 3.76E-03 |
| *Tnfrsf9* | 9.68 | 1.88 | 0.58 | -3.22 | 1.26E-03 | 1.42E-02 |
| *Itgax* | 44.92 | 1.88 | 0.35 | -5.40 | 6.85E-08 | 4.89E-06 |
| *Kcnj10* | 16.18 | 1.87 | 0.58 | -3.22 | 1.28E-03 | 1.43E-02 |
| *H2-T24* | 16.83 | 1.87 | 0.47 | -3.97 | 7.21E-05 | 1.60E-03 |
| *Kcnq5* | 9.14 | 1.86 | 0.65 | -2.85 | 4.40E-03 | 3.49E-02 |
| *Hcn3* | 8.81 | 1.86 | 0.67 | -2.78 | 5.44E-03 | 4.06E-02 |
| *H2-K1* | 489.55 | 1.86 | 0.23 | -8.00 | 1.30E-15 | 6.75E-13 |
| *AC133103.2* | 6.80 | 1.86 | 0.67 | -2.77 | 5.57E-03 | 4.13E-02 |
| *Gm17024* | 73.57 | 1.85 | 0.51 | -3.61 | 3.10E-04 | 4.98E-03 |
| *St6gal2* | 15.47 | 1.85 | 0.61 | -3.04 | 2.35E-03 | 2.24E-02 |

| | | | | | | |
|---|---|---|---|---|---|---|
| *Ankrd37* | 22.41 | 1.85 | 0.45 | -4.11 | 3.95E-05 | 9.76E-04 |
| *Btla* | 6.97 | 1.85 | 0.68 | -2.71 | 6.83E-03 | 4.78E-02 |
| *Trp73* | 66.65 | 1.85 | 0.32 | -5.82 | 5.93E-09 | 5.70E-07 |
| *Rgs5* | 118.06 | 1.85 | 0.47 | -3.95 | 7.79E-05 | 1.69E-03 |
| *Ccl6* | 43.55 | 1.85 | 0.40 | -4.58 | 4.75E-06 | 1.68E-04 |
| *Oas1g* | 258.89 | 1.84 | 0.25 | -7.42 | 1.15E-13 | 4.38E-11 |
| *Fcer1g* | 32.65 | 1.84 | 0.36 | -5.08 | 3.74E-07 | 2.04E-05 |
| *Dlgap3* | 26.95 | 1.84 | 0.52 | -3.52 | 4.29E-04 | 6.40E-03 |
| *Ifi203* | 1023.32 | 1.84 | 0.29 | -6.33 | 2.38E-10 | 3.59E-08 |
| *Eif4e3* | 58.89 | 1.84 | 0.32 | -5.76 | 8.50E-09 | 7.90E-07 |
| *Mlana* | 55.60 | 1.84 | 0.60 | -3.05 | 2.32E-03 | 2.21E-02 |
| *Abcb4* | 14.74 | 1.83 | 0.66 | -2.77 | 5.58E-03 | 4.14E-02 |
| *Tbxa2r* | 6.73 | 1.83 | 0.62 | -2.96 | 3.05E-03 | 2.69E-02 |
| *Amer2* | 36.56 | 1.83 | 0.59 | -3.10 | 1.95E-03 | 1.95E-02 |
| *Slfn2* | 1649.18 | 1.83 | 0.20 | -8.96 | 3.16E-19 | 3.13E-16 |
| *Ier5l* | 245.70 | 1.83 | 0.30 | -6.17 | 6.85E-10 | 8.92E-08 |
| *Six3* | 44.75 | 1.83 | 0.48 | -3.79 | 1.52E-04 | 2.85E-03 |
| *Fscn2* | 57.52 | 1.83 | 0.46 | -4.01 | 6.05E-05 | 1.39E-03 |
| *Ms4a6d* | 35.56 | 1.82 | 0.41 | -4.47 | 7.91E-06 | 2.58E-04 |
| *AC127341.3* | 62.24 | 1.82 | 0.45 | -4.06 | 4.88E-05 | 1.16E-03 |
| *Plk3* | 1246.43 | 1.82 | 0.51 | -3.54 | 4.05E-04 | 6.13E-03 |
| *Pla2g2e* | 9.17 | 1.82 | 0.67 | -2.70 | 6.83E-03 | 4.79E-02 |
| *Cd48* | 10.31 | 1.82 | 0.51 | -3.57 | 3.57E-04 | 5.53E-03 |
| *Fam161a* | 47.98 | 1.81 | 0.58 | -3.12 | 1.79E-03 | 1.83E-02 |
| *Nr4a3* | 54.01 | 1.81 | 0.42 | -4.32 | 1.59E-05 | 4.70E-04 |
| *Gm11517* | 6.40 | 1.81 | 0.66 | -2.72 | 6.53E-03 | 4.64E-02 |
| *Ifitm3* | 1832.03 | 1.80 | 0.23 | -7.95 | 1.89E-15 | 9.37E-13 |
| *Hba-a1* | 301.44 | 1.80 | 0.32 | -5.55 | 2.90E-08 | 2.34E-06 |
| *Flt1* | 68.59 | 1.80 | 0.47 | -3.80 | 1.44E-04 | 2.72E-03 |
| *Galnt6* | 8.97 | 1.80 | 0.56 | -3.23 | 1.23E-03 | 1.39E-02 |
| *Shisal1* | 20.06 | 1.80 | 0.46 | -3.86 | 1.12E-04 | 2.25E-03 |
| *Ptprcap* | 18.11 | 1.80 | 0.40 | -4.51 | 6.43E-06 | 2.18E-04 |
| *Slc6a13* | 25.83 | 1.79 | 0.63 | -2.84 | 4.51E-03 | 3.55E-02 |
| *Tcfl5* | 11.00 | 1.79 | 0.56 | -3.19 | 1.40E-03 | 1.53E-02 |
| *Ppp1r16b* | 20.60 | 1.79 | 0.48 | -3.76 | 1.67E-04 | 3.09E-03 |
| *9130208D14Rik* | 26.72 | 1.79 | 0.60 | -2.98 | 2.90E-03 | 2.60E-02 |
| *Cd84* | 13.17 | 1.79 | 0.51 | -3.49 | 4.90E-04 | 7.05E-03 |
| *Disp2* | 26.38 | 1.78 | 0.53 | -3.36 | 7.81E-04 | 1.01E-02 |
| *Pthlh* | 1202.28 | 1.78 | 0.18 | -9.89 | 4.60E-23 | 6.07E-20 |

| | | | | | | |
|---|---|---|---|---|---|---|
| *Slc15a3* | 17.78 | 1.78 | 0.46 | -3.84 | 1.26E-04 | 2.45E-03 |
| *Snca* | 11.01 | 1.78 | 0.54 | -3.27 | 1.09E-03 | 1.27E-02 |
| *Evi2a* | 9.59 | 1.77 | 0.64 | -2.76 | 5.81E-03 | 4.26E-02 |
| *Trim6* | 71.56 | 1.77 | 0.27 | -6.52 | 7.24E-11 | 1.23E-08 |
| *Hsh2d* | 25.61 | 1.77 | 0.40 | -4.39 | 1.13E-05 | 3.51E-04 |
| *Fcgr4* | 8.78 | 1.76 | 0.60 | -2.93 | 3.37E-03 | 2.89E-02 |
| *Gpr158* | 12.55 | 1.76 | 0.61 | -2.88 | 3.97E-03 | 3.24E-02 |
| *Ptprc* | 84.15 | 1.76 | 0.30 | -5.89 | 3.96E-09 | 3.96E-07 |
| *Trim30a* | 1424.99 | 1.76 | 0.28 | -6.35 | 2.13E-10 | 3.29E-08 |
| *Gm10434* | 7.54 | 1.75 | 0.62 | -2.83 | 4.70E-03 | 3.65E-02 |
| *Trim30d* | 136.25 | 1.74 | 0.35 | -5.00 | 5.71E-07 | 2.87E-05 |
| *Cd4* | 22.93 | 1.74 | 0.45 | -3.86 | 1.14E-04 | 2.28E-03 |
| *Crmp1* | 28.29 | 1.74 | 0.55 | -3.18 | 1.45E-03 | 1.56E-02 |
| *Kcnc1* | 36.40 | 1.74 | 0.53 | -3.30 | 9.76E-04 | 1.17E-02 |
| *Traf3ip3* | 13.40 | 1.73 | 0.59 | -2.92 | 3.48E-03 | 2.96E-02 |
| *Gm4610* | 174.55 | 1.73 | 0.42 | -4.16 | 3.21E-05 | 8.22E-04 |
| *Gm44645* | 9.96 | 1.73 | 0.56 | -3.08 | 2.10E-03 | 2.05E-02 |
| *Barx2* | 193.72 | 1.73 | 0.45 | -3.84 | 1.24E-04 | 2.43E-03 |
| *Gm11714* | 15.04 | 1.73 | 0.55 | -3.14 | 1.68E-03 | 1.74E-02 |
| *Gem* | 58.48 | 1.73 | 0.39 | -4.42 | 1.01E-05 | 3.18E-04 |
| *AB124611* | 13.76 | 1.73 | 0.49 | -3.50 | 4.61E-04 | 6.75E-03 |
| *Pcdh8* | 57.66 | 1.73 | 0.37 | -4.61 | 4.03E-06 | 1.48E-04 |
| *Il2rg* | 18.99 | 1.73 | 0.51 | -3.40 | 6.69E-04 | 9.00E-03 |
| *Plcl1* | 12.49 | 1.72 | 0.53 | -3.26 | 1.12E-03 | 1.30E-02 |
| *Nefh* | 23.12 | 1.72 | 0.54 | -3.21 | 1.33E-03 | 1.47E-02 |
| *Krt17* | 608.70 | 1.71 | 0.60 | -2.86 | 4.24E-03 | 3.40E-02 |
| *Mef2c* | 52.12 | 1.71 | 0.43 | -3.98 | 6.85E-05 | 1.54E-03 |
| *Vegfa* | 1270.06 | 1.71 | 0.36 | -4.78 | 1.72E-06 | 7.42E-05 |
| *Tspan11* | 17.15 | 1.71 | 0.45 | -3.79 | 1.48E-04 | 2.79E-03 |
| *Add2* | 22.76 | 1.71 | 0.46 | -3.68 | 2.29E-04 | 3.94E-03 |
| *Bbc3* | 46.30 | 1.70 | 0.37 | -4.61 | 4.04E-06 | 1.48E-04 |
| *Thy1* | 122.90 | 1.70 | 0.32 | -5.26 | 1.42E-07 | 9.01E-06 |
| *Hmga2* | 22.10 | 1.70 | 0.60 | -2.85 | 4.42E-03 | 3.50E-02 |
| *Socs3* | 220.94 | 1.70 | 0.41 | -4.18 | 2.89E-05 | 7.54E-04 |
| *Nptx1* | 23.89 | 1.70 | 0.50 | -3.38 | 7.28E-04 | 9.55E-03 |
| *Vat1l* | 56.22 | 1.69 | 0.59 | -2.88 | 3.95E-03 | 3.23E-02 |
| *Ildr2* | 16.95 | 1.69 | 0.56 | -3.05 | 2.33E-03 | 2.22E-02 |
| *Tagln3* | 22.13 | 1.69 | 0.56 | -3.01 | 2.58E-03 | 2.38E-02 |
| *Fcgr1* | 25.79 | 1.69 | 0.46 | -3.71 | 2.05E-04 | 3.63E-03 |

| | | | | | | |
|---|---|---|---|---|---|---|
| *Edn2* | 19.55 | 1.69 | 0.53 | -3.18 | 1.48E-03 | 1.58E-02 |
| *Fabp5* | 3978.14 | 1.69 | 0.25 | -6.66 | 2.73E-11 | 5.24E-09 |
| *Samhd1* | 2750.19 | 1.69 | 0.16 | -10.78 | 4.41E-27 | 8.73E-24 |
| *Six3os1* | 29.33 | 1.69 | 0.52 | -3.25 | 1.17E-03 | 1.33E-02 |
| *Oas1a* | 994.26 | 1.68 | 0.21 | -7.84 | 4.66E-15 | 2.14E-12 |
| *Crnn* | 34.53 | 1.68 | 0.47 | -3.61 | 3.02E-04 | 4.89E-03 |
| *Icos* | 19.23 | 1.68 | 0.54 | -3.10 | 1.93E-03 | 1.93E-02 |
| *Enpp2* | 201.62 | 1.68 | 0.35 | -4.75 | 2.05E-06 | 8.50E-05 |
| *Fam78a* | 11.52 | 1.68 | 0.61 | -2.76 | 5.72E-03 | 4.21E-02 |
| *Skida1* | 26.77 | 1.67 | 0.55 | -3.04 | 2.40E-03 | 2.27E-02 |
| *Sprn* | 7.06 | 1.67 | 0.58 | -2.89 | 3.90E-03 | 3.20E-02 |
| *Vwf* | 54.87 | 1.67 | 0.48 | -3.51 | 4.45E-04 | 6.59E-03 |
| *Atp2b2* | 34.13 | 1.67 | 0.53 | -3.16 | 1.55E-03 | 1.64E-02 |
| *Stra6* | 75.08 | 1.66 | 0.40 | -4.16 | 3.12E-05 | 8.04E-04 |
| *H2-Q10* | 7.85 | 1.66 | 0.61 | -2.71 | 6.78E-03 | 4.76E-02 |
| *Hey1* | 46.42 | 1.66 | 0.47 | -3.50 | 4.63E-04 | 6.77E-03 |
| *Celf3* | 82.48 | 1.65 | 0.31 | -5.24 | 1.57E-07 | 9.84E-06 |
| *Runx3* | 14.99 | 1.64 | 0.50 | -3.28 | 1.03E-03 | 1.23E-02 |
| *Sfta2* | 19.67 | 1.64 | 0.57 | -2.90 | 3.70E-03 | 3.09E-02 |
| *Arhgap9* | 15.08 | 1.64 | 0.48 | -3.39 | 6.92E-04 | 9.20E-03 |
| *H2-Q4* | 1355.62 | 1.64 | 0.19 | -8.46 | 2.60E-17 | 1.78E-14 |
| *Tcp11* | 15.15 | 1.64 | 0.49 | -3.34 | 8.48E-04 | 1.06E-02 |
| *Elfn1* | 89.39 | 1.64 | 0.33 | -4.96 | 7.08E-07 | 3.45E-05 |
| *Akr1b8* | 542.70 | 1.63 | 0.28 | -5.83 | 5.39E-09 | 5.20E-07 |
| *Sp100* | 1231.15 | 1.63 | 0.27 | -6.09 | 1.11E-09 | 1.34E-07 |
| *Cyyr1* | 16.31 | 1.63 | 0.55 | -2.95 | 3.16E-03 | 2.75E-02 |
| *Sox9* | 498.68 | 1.63 | 0.24 | -6.88 | 5.90E-12 | 1.33E-09 |
| *Nr2f1* | 21.37 | 1.63 | 0.50 | -3.27 | 1.07E-03 | 1.26E-02 |
| *Gpx2* | 19.99 | 1.63 | 0.51 | -3.20 | 1.35E-03 | 1.49E-02 |
| *Gpr183* | 13.28 | 1.62 | 0.47 | -3.44 | 5.89E-04 | 8.19E-03 |
| *Myo1g* | 23.97 | 1.62 | 0.36 | -4.55 | 5.43E-06 | 1.88E-04 |
| *Krt14* | 27272.05 | 1.62 | 0.28 | -5.86 | 4.51E-09 | 4.44E-07 |
| *Klf10* | 1004.81 | 1.61 | 0.33 | -4.92 | 8.80E-07 | 4.19E-05 |
| *Tnfrsf12a* | 1068.37 | 1.61 | 0.34 | -4.76 | 1.95E-06 | 8.18E-05 |
| *AC135964.2* | 53.56 | 1.61 | 0.27 | -5.94 | 2.92E-09 | 3.07E-07 |
| *Nim1k* | 37.04 | 1.61 | 0.43 | -3.73 | 1.95E-04 | 3.49E-03 |
| *Slc6a1* | 72.08 | 1.61 | 0.50 | -3.22 | 1.30E-03 | 1.44E-02 |
| *Dhx58* | 818.40 | 1.61 | 0.22 | -7.22 | 5.36E-13 | 1.63E-10 |
| *Pacsin1* | 74.54 | 1.60 | 0.53 | -3.05 | 2.27E-03 | 2.18E-02 |

| | | | | | | |
|---|---|---|---|---|---|---|
| *Gbp4* | 879.26 | 1.60 | 0.24 | -6.59 | 4.31E-11 | 7.97E-09 |
| *Necab2* | 26.55 | 1.60 | 0.55 | -2.93 | 3.40E-03 | 2.91E-02 |
| *Casp4* | 28.72 | 1.60 | 0.46 | -3.49 | 4.75E-04 | 6.90E-03 |
| *H2-Aa* | 99.25 | 1.59 | 0.27 | -5.91 | 3.37E-09 | 3.48E-07 |
| *Areg* | 164.25 | 1.59 | 0.30 | -5.38 | 7.53E-08 | 5.33E-06 |
| *Ifi207* | 22.78 | 1.59 | 0.48 | -3.30 | 9.53E-04 | 1.15E-02 |
| *Gabrb2* | 31.98 | 1.59 | 0.37 | -4.35 | 1.34E-05 | 4.07E-04 |
| *Gm11127* | 11.60 | 1.59 | 0.56 | -2.83 | 4.67E-03 | 3.64E-02 |
| *Rac2* | 52.93 | 1.59 | 0.37 | -4.32 | 1.59E-05 | 4.70E-04 |
| *Myrip* | 33.39 | 1.58 | 0.58 | -2.74 | 6.22E-03 | 4.49E-02 |
| *Ciart* | 313.40 | 1.58 | 0.31 | -5.07 | 3.95E-07 | 2.14E-05 |
| *Nrcam* | 20.88 | 1.58 | 0.49 | -3.22 | 1.26E-03 | 1.42E-02 |
| *Ifrd1* | 2138.63 | 1.58 | 0.27 | -5.78 | 7.27E-09 | 6.86E-07 |
| *Fscn1* | 2003.91 | 1.58 | 0.28 | -5.62 | 1.92E-08 | 1.63E-06 |
| *Itpripl1* | 101.84 | 1.58 | 0.24 | -6.69 | 2.24E-11 | 4.48E-09 |
| *Sla* | 33.88 | 1.57 | 0.34 | -4.68 | 2.86E-06 | 1.12E-04 |
| *H2-M3* | 103.50 | 1.57 | 0.26 | -6.04 | 1.53E-09 | 1.75E-07 |
| *Ccnd2* | 1773.09 | 1.57 | 0.25 | -6.27 | 3.50E-10 | 5.02E-08 |
| *Tubb3* | 43.41 | 1.57 | 0.48 | -3.30 | 9.74E-04 | 1.17E-02 |
| *Ifit3b* | 1260.95 | 1.57 | 0.26 | -6.08 | 1.22E-09 | 1.45E-07 |
| *Bmp2* | 50.03 | 1.56 | 0.45 | -3.49 | 4.80E-04 | 6.94E-03 |
| *Mndal* | 481.75 | 1.56 | 0.21 | -7.34 | 2.19E-13 | 7.34E-11 |
| *Ramp2* | 53.65 | 1.55 | 0.33 | -4.69 | 2.67E-06 | 1.06E-04 |
| *Pxdc1* | 2713.07 | 1.55 | 0.24 | -6.36 | 1.96E-10 | 3.07E-08 |
| *Impdh1* | 212.44 | 1.54 | 0.39 | -3.96 | 7.63E-05 | 1.67E-03 |
| *Myc* | 733.58 | 1.53 | 0.29 | -5.33 | 9.96E-08 | 6.80E-06 |
| *Fblim1* | 112.67 | 1.53 | 0.36 | -4.31 | 1.66E-05 | 4.87E-04 |
| *Psmb9* | 1132.48 | 1.53 | 0.24 | -6.48 | 9.48E-11 | 1.56E-08 |
| *Slc16a3* | 105.11 | 1.53 | 0.35 | -4.36 | 1.28E-05 | 3.91E-04 |
| *Socs1* | 122.19 | 1.53 | 0.34 | -4.55 | 5.46E-06 | 1.89E-04 |
| *Neurl1a* | 42.69 | 1.53 | 0.44 | -3.46 | 5.36E-04 | 7.61E-03 |
| *Dram1* | 22.93 | 1.52 | 0.48 | -3.20 | 1.39E-03 | 1.52E-02 |
| *Mlkl* | 263.65 | 1.52 | 0.18 | -8.47 | 2.38E-17 | 1.68E-14 |
| *Tm4sf1* | 283.23 | 1.52 | 0.40 | -3.80 | 1.43E-04 | 2.70E-03 |
| *Tbc1d9* | 64.78 | 1.52 | 0.44 | -3.44 | 5.86E-04 | 8.16E-03 |
| *Kcne1* | 16.83 | 1.52 | 0.43 | -3.51 | 4.54E-04 | 6.68E-03 |
| *Gimap8* | 14.37 | 1.51 | 0.50 | -3.05 | 2.31E-03 | 2.21E-02 |
| *Lcp1* | 158.88 | 1.51 | 0.27 | -5.70 | 1.20E-08 | 1.06E-06 |
| *Irf4* | 16.24 | 1.51 | 0.55 | -2.76 | 5.81E-03 | 4.26E-02 |

| | | | | | | |
|---|---|---|---|---|---|---|
| *Ampd1* | 24.05 | 1.51 | 0.44 | -3.46 | 5.46E-04 | 7.72E-03 |
| *Agap2* | 27.26 | 1.50 | 0.48 | -3.12 | 1.79E-03 | 1.83E-02 |
| *Clec7a* | 17.49 | 1.50 | 0.54 | -2.76 | 5.82E-03 | 4.26E-02 |
| *Nmnat2* | 16.87 | 1.50 | 0.49 | -3.04 | 2.38E-03 | 2.25E-02 |
| *Iqsec3* | 52.67 | 1.50 | 0.55 | -2.73 | 6.40E-03 | 4.58E-02 |
| *Coro1a* | 120.80 | 1.50 | 0.25 | -5.97 | 2.38E-09 | 2.56E-07 |
| *Trf* | 25.80 | 1.50 | 0.48 | -3.10 | 1.94E-03 | 1.94E-02 |
| *Camk1g* | 35.33 | 1.49 | 0.41 | -3.68 | 2.29E-04 | 3.94E-03 |
| *Ms4a6b* | 38.24 | 1.49 | 0.40 | -3.77 | 1.65E-04 | 3.06E-03 |
| *Angpt1* | 17.81 | 1.49 | 0.38 | -3.89 | 1.02E-04 | 2.09E-03 |
| *Ly6a* | 4797.42 | 1.49 | 0.22 | -6.87 | 6.22E-12 | 1.38E-09 |
| *Helz2* | 610.88 | 1.49 | 0.28 | -5.34 | 9.31E-08 | 6.38E-06 |
| *Mlf1* | 37.09 | 1.48 | 0.44 | -3.40 | 6.78E-04 | 9.09E-03 |
| *Cybb* | 82.04 | 1.47 | 0.29 | -5.15 | 2.65E-07 | 1.55E-05 |
| *Podxl* | 113.49 | 1.47 | 0.38 | -3.86 | 1.13E-04 | 2.28E-03 |
| *Lag3* | 15.43 | 1.47 | 0.40 | -3.71 | 2.08E-04 | 3.66E-03 |
| *Ly6c1* | 150.53 | 1.47 | 0.24 | -6.22 | 4.94E-10 | 6.71E-08 |
| *Csf2rb2* | 25.64 | 1.47 | 0.40 | -3.62 | 2.91E-04 | 4.76E-03 |
| *Oasl2* | 5155.39 | 1.47 | 0.21 | -6.98 | 2.92E-12 | 7.50E-10 |
| *Prr7* | 22.05 | 1.46 | 0.48 | -3.02 | 2.49E-03 | 2.32E-02 |
| *Baalc* | 13.05 | 1.46 | 0.46 | -3.19 | 1.41E-03 | 1.53E-02 |
| *Hs3st1* | 219.62 | 1.46 | 0.38 | -3.79 | 1.51E-04 | 2.82E-03 |
| *Rbm38* | 52.97 | 1.46 | 0.30 | -4.91 | 9.03E-07 | 4.29E-05 |
| *Kcnab2* | 96.22 | 1.46 | 0.35 | -4.19 | 2.75E-05 | 7.25E-04 |
| *Xaf1* | 693.58 | 1.45 | 0.21 | -6.87 | 6.28E-12 | 1.38E-09 |
| *Hbb-bs* | 211.12 | 1.44 | 0.29 | -5.02 | 5.16E-07 | 2.63E-05 |
| *Dock2* | 40.15 | 1.44 | 0.35 | -4.07 | 4.70E-05 | 1.13E-03 |
| *Sema3g* | 27.66 | 1.44 | 0.46 | -3.14 | 1.69E-03 | 1.75E-02 |
| *Ina* | 30.40 | 1.43 | 0.49 | -2.95 | 3.20E-03 | 2.78E-02 |
| *Tnfrsf4* | 18.41 | 1.43 | 0.46 | -3.12 | 1.83E-03 | 1.86E-02 |
| *Clec14a* | 25.04 | 1.43 | 0.53 | -2.70 | 6.99E-03 | 4.86E-02 |
| *Bhlhe41* | 441.16 | 1.43 | 0.31 | -4.66 | 3.12E-06 | 1.20E-04 |
| *Steap1* | 76.31 | 1.43 | 0.51 | -2.80 | 5.06E-03 | 3.86E-02 |
| *Gimap1* | 12.03 | 1.43 | 0.51 | -2.79 | 5.31E-03 | 4.00E-02 |
| *Hspb8* | 99.73 | 1.42 | 0.34 | -4.23 | 2.31E-05 | 6.30E-04 |
| *Gpr132* | 19.24 | 1.42 | 0.36 | -3.89 | 1.00E-04 | 2.06E-03 |
| *Ly75* | 201.93 | 1.42 | 0.30 | -4.67 | 3.08E-06 | 1.20E-04 |
| *Lcp2* | 14.42 | 1.42 | 0.48 | -2.95 | 3.22E-03 | 2.79E-02 |
| *Mir17hg* | 137.92 | 1.42 | 0.29 | -4.86 | 1.19E-06 | 5.45E-05 |

| | | | | | | |
|---|---|---|---|---|---|---|
| *Stat1* | 841.80 | 1.41 | 0.28 | -5.12 | 3.05E-07 | 1.74E-05 |
| *Dner* | 32.83 | 1.41 | 0.41 | -3.40 | 6.77E-04 | 9.08E-03 |
| *Cspg5* | 64.45 | 1.41 | 0.48 | -2.92 | 3.50E-03 | 2.97E-02 |
| *Kcnrg* | 34.11 | 1.41 | 0.33 | -4.30 | 1.72E-05 | 5.00E-04 |
| *Elovl4* | 169.99 | 1.40 | 0.43 | -3.24 | 1.21E-03 | 1.37E-02 |
| *Ms4a6c* | 17.40 | 1.40 | 0.49 | -2.84 | 4.49E-03 | 3.54E-02 |
| *Tub* | 96.59 | 1.40 | 0.47 | -2.98 | 2.90E-03 | 2.60E-02 |
| *1700017B05Rik* | 701.28 | 1.39 | 0.18 | -7.63 | 2.42E-14 | 9.97E-12 |
| *Hk2* | 1238.59 | 1.39 | 0.31 | -4.47 | 7.70E-06 | 2.52E-04 |
| *Gm8995* | 842.74 | 1.39 | 0.36 | -3.91 | 9.20E-05 | 1.95E-03 |
| *Apobec2* | 113.87 | 1.39 | 0.40 | -3.49 | 4.84E-04 | 6.98E-03 |
| *Acacb* | 45.55 | 1.39 | 0.45 | -3.09 | 2.00E-03 | 1.98E-02 |
| *Nat8l* | 384.52 | 1.39 | 0.38 | -3.66 | 2.49E-04 | 4.21E-03 |
| *Cplx1* | 46.21 | 1.39 | 0.44 | -3.15 | 1.62E-03 | 1.70E-02 |
| *3-Sep* | 32.94 | 1.39 | 0.45 | -3.08 | 2.04E-03 | 2.00E-02 |
| *Spink5* | 2366.94 | 1.39 | 0.27 | -5.12 | 3.06E-07 | 1.74E-05 |
| *Ube2l6* | 1191.09 | 1.38 | 0.21 | -6.60 | 4.16E-11 | 7.78E-09 |
| *Rnf224* | 86.51 | 1.38 | 0.30 | -4.66 | 3.18E-06 | 1.23E-04 |
| *Lman1l* | 20.59 | 1.38 | 0.49 | -2.79 | 5.32E-03 | 4.00E-02 |
| *Spn* | 45.29 | 1.37 | 0.30 | -4.64 | 3.46E-06 | 1.32E-04 |
| *Emilin2* | 45.11 | 1.37 | 0.43 | -3.22 | 1.30E-03 | 1.45E-02 |
| *Ifi205* | 14.55 | 1.37 | 0.49 | -2.83 | 4.64E-03 | 3.62E-02 |
| *S1pr1* | 33.45 | 1.37 | 0.39 | -3.52 | 4.25E-04 | 6.36E-03 |
| *Abcb1b* | 13.93 | 1.37 | 0.50 | -2.75 | 6.04E-03 | 4.38E-02 |
| *Il10ra* | 31.21 | 1.37 | 0.34 | -4.07 | 4.79E-05 | 1.15E-03 |
| *Spata20* | 18.91 | 1.37 | 0.44 | -3.11 | 1.89E-03 | 1.91E-02 |
| *Slc4a8* | 32.81 | 1.37 | 0.43 | -3.18 | 1.46E-03 | 1.56E-02 |
| *Arhgap30* | 75.86 | 1.36 | 0.34 | -3.96 | 7.64E-05 | 1.67E-03 |
| *Tmc7* | 142.29 | 1.36 | 0.37 | -3.68 | 2.33E-04 | 3.99E-03 |
| *Ccne1* | 227.60 | 1.36 | 0.27 | -5.00 | 5.65E-07 | 2.85E-05 |
| *Ifi203-ps* | 54.95 | 1.35 | 0.38 | -3.55 | 3.83E-04 | 5.86E-03 |
| *Nrsn1* | 28.35 | 1.35 | 0.43 | -3.14 | 1.67E-03 | 1.74E-02 |
| *Itgb2* | 45.57 | 1.35 | 0.32 | -4.23 | 2.37E-05 | 6.40E-04 |
| *Slc12a5* | 172.69 | 1.35 | 0.45 | -2.98 | 2.89E-03 | 2.59E-02 |
| *Apc2* | 33.72 | 1.34 | 0.47 | -2.84 | 4.57E-03 | 3.58E-02 |
| *Neurl3* | 26.54 | 1.34 | 0.31 | -4.35 | 1.38E-05 | 4.17E-04 |
| *Csf2rb* | 56.48 | 1.34 | 0.30 | -4.52 | 6.11E-06 | 2.09E-04 |
| *Otud1* | 1733.96 | 1.34 | 0.28 | -4.84 | 1.28E-06 | 5.76E-05 |
| *Cldn2* | 26.69 | 1.34 | 0.48 | -2.81 | 4.95E-03 | 3.80E-02 |

| | | | | | | |
|---|---|---|---|---|---|---|
| *Calm4* | 32.02 | 1.34 | 0.39 | -3.41 | 6.60E-04 | 8.90E-03 |
| *Zfp365* | 53.72 | 1.34 | 0.39 | -3.44 | 5.89E-04 | 8.19E-03 |
| *Tie1* | 40.97 | 1.33 | 0.49 | -2.75 | 6.02E-03 | 4.38E-02 |
| *Basp1* | 95.97 | 1.33 | 0.27 | -4.93 | 8.17E-07 | 3.95E-05 |
| *H2-Q1* | 39.83 | 1.33 | 0.30 | -4.47 | 7.81E-06 | 2.55E-04 |
| *Atp10a* | 33.78 | 1.33 | 0.40 | -3.35 | 8.15E-04 | 1.03E-02 |
| *Fyb* | 26.27 | 1.33 | 0.31 | -4.27 | 1.92E-05 | 5.48E-04 |
| *Rgcc* | 28.13 | 1.32 | 0.37 | -3.53 | 4.13E-04 | 6.20E-03 |
| *Zfp593* | 23.57 | 1.32 | 0.38 | -3.50 | 4.71E-04 | 6.86E-03 |
| *Unc119* | 441.33 | 1.32 | 0.39 | -3.39 | 6.96E-04 | 9.26E-03 |
| *Vav1* | 25.44 | 1.31 | 0.44 | -2.97 | 3.01E-03 | 2.67E-02 |
| *Sectm1b* | 45.53 | 1.31 | 0.43 | -3.04 | 2.37E-03 | 2.25E-02 |
| *Aldh1b1* | 13.35 | 1.31 | 0.42 | -3.09 | 2.03E-03 | 2.00E-02 |
| *Gm38312* | 17.93 | 1.31 | 0.48 | -2.72 | 6.60E-03 | 4.68E-02 |
| *Herc6* | 486.01 | 1.31 | 0.27 | -4.79 | 1.63E-06 | 7.08E-05 |
| *Cd93* | 39.41 | 1.31 | 0.46 | -2.83 | 4.60E-03 | 3.60E-02 |
| *Hap1* | 181.98 | 1.31 | 0.20 | -6.55 | 5.84E-11 | 1.02E-08 |
| *Nes* | 366.76 | 1.31 | 0.24 | -5.48 | 4.36E-08 | 3.37E-06 |
| *Fhl3* | 149.87 | 1.31 | 0.39 | -3.32 | 9.02E-04 | 1.10E-02 |
| *Ifi47* | 1224.07 | 1.30 | 0.22 | -6.00 | 2.00E-09 | 2.20E-07 |
| *Gprc5b* | 118.09 | 1.30 | 0.26 | -4.96 | 7.17E-07 | 3.49E-05 |
| *Acta2* | 246.55 | 1.30 | 0.34 | -3.84 | 1.21E-04 | 2.37E-03 |
| *Mmrn2* | 33.66 | 1.30 | 0.46 | -2.84 | 4.57E-03 | 3.58E-02 |
| *Dnajb4* | 1538.13 | 1.30 | 0.20 | -6.52 | 6.99E-11 | 1.19E-08 |
| *Vtcn1* | 37.50 | 1.30 | 0.39 | -3.37 | 7.57E-04 | 9.83E-03 |
| *Sptbn4* | 23.62 | 1.29 | 0.43 | -2.98 | 2.86E-03 | 2.57E-02 |
| *Zdhhc2* | 71.43 | 1.29 | 0.34 | -3.75 | 1.76E-04 | 3.21E-03 |
| *Fbxl22* | 24.45 | 1.29 | 0.48 | -2.71 | 6.81E-03 | 4.78E-02 |
| *Ptpn7* | 15.91 | 1.29 | 0.47 | -2.76 | 5.82E-03 | 4.26E-02 |
| *Epgn* | 706.19 | 1.29 | 0.28 | -4.57 | 4.83E-06 | 1.70E-04 |
| *Pde1b* | 32.96 | 1.28 | 0.37 | -3.43 | 6.08E-04 | 8.39E-03 |
| *Ly86* | 27.16 | 1.28 | 0.41 | -3.09 | 2.02E-03 | 1.99E-02 |
| *Tdh* | 15.19 | 1.28 | 0.41 | -3.11 | 1.85E-03 | 1.87E-02 |
| *Myo1f* | 25.73 | 1.28 | 0.41 | -3.09 | 2.00E-03 | 1.98E-02 |
| *Ifit1bl2* | 69.26 | 1.27 | 0.27 | -4.69 | 2.69E-06 | 1.07E-04 |
| *Cd38* | 17.39 | 1.27 | 0.42 | -3.02 | 2.50E-03 | 2.33E-02 |
| *Sptssb* | 66.12 | 1.27 | 0.42 | -2.99 | 2.76E-03 | 2.50E-02 |
| *Lrmp* | 19.51 | 1.27 | 0.44 | -2.90 | 3.68E-03 | 3.08E-02 |
| *Cspg4* | 82.19 | 1.26 | 0.31 | -4.05 | 5.13E-05 | 1.21E-03 |

| | | | | | | |
|---|---|---|---|---|---|---|
| *Ikzf1* | 29.15 | 1.26 | 0.42 | -2.98 | 2.91E-03 | 2.60E-02 |
| *Col9a3* | 74.39 | 1.26 | 0.31 | -4.07 | 4.74E-05 | 1.14E-03 |
| *Il1b* | 35.33 | 1.25 | 0.35 | -3.55 | 3.90E-04 | 5.96E-03 |
| *Tap1* | 2285.29 | 1.25 | 0.21 | -6.04 | 1.50E-09 | 1.74E-07 |
| *Scn8a* | 31.57 | 1.25 | 0.46 | -2.70 | 6.85E-03 | 4.79E-02 |
| *Hspa2* | 142.13 | 1.23 | 0.17 | -7.40 | 1.38E-13 | 5.06E-11 |
| *Hcls1* | 72.37 | 1.23 | 0.26 | -4.79 | 1.70E-06 | 7.35E-05 |
| *Rhof* | 84.60 | 1.23 | 0.39 | -3.19 | 1.41E-03 | 1.53E-02 |
| *Atp2a3* | 29.64 | 1.23 | 0.31 | -4.02 | 5.88E-05 | 1.35E-03 |
| *Plvap* | 108.55 | 1.23 | 0.30 | -4.15 | 3.26E-05 | 8.31E-04 |
| *Calb2* | 109.04 | 1.23 | 0.45 | -2.71 | 6.80E-03 | 4.77E-02 |
| *Scn1b* | 99.82 | 1.22 | 0.43 | -2.82 | 4.82E-03 | 3.72E-02 |
| *Gnai1* | 31.03 | 1.22 | 0.41 | -3.00 | 2.69E-03 | 2.45E-02 |
| *Stxbp1* | 342.88 | 1.21 | 0.38 | -3.19 | 1.44E-03 | 1.55E-02 |
| *Cdk5r1* | 39.69 | 1.21 | 0.36 | -3.37 | 7.64E-04 | 9.89E-03 |
| *Fbxl16* | 35.84 | 1.20 | 0.43 | -2.78 | 5.35E-03 | 4.02E-02 |
| *Cyth4* | 51.95 | 1.20 | 0.40 | -2.97 | 2.94E-03 | 2.62E-02 |
| *Tead4* | 195.81 | 1.20 | 0.44 | -2.70 | 6.96E-03 | 4.85E-02 |
| *Abl2* | 1823.26 | 1.20 | 0.33 | -3.58 | 3.46E-04 | 5.42E-03 |
| *Serpinb6b* | 77.92 | 1.19 | 0.29 | -4.07 | 4.68E-05 | 1.13E-03 |
| *Aph1c* | 30.67 | 1.19 | 0.39 | -3.08 | 2.08E-03 | 2.03E-02 |
| *Sema7a* | 81.84 | 1.19 | 0.44 | -2.73 | 6.34E-03 | 4.54E-02 |
| *Trib2* | 607.74 | 1.19 | 0.17 | -6.86 | 6.85E-12 | 1.47E-09 |
| *Il15* | 37.48 | 1.18 | 0.43 | -2.78 | 5.44E-03 | 4.06E-02 |
| *Ifi202b* | 9754.34 | 1.18 | 0.20 | -5.89 | 3.85E-09 | 3.89E-07 |
| *Creb5* | 54.09 | 1.18 | 0.39 | -3.07 | 2.16E-03 | 2.10E-02 |
| *Id3* | 816.52 | 1.18 | 0.25 | -4.75 | 2.03E-06 | 8.45E-05 |
| *Ier2* | 2963.03 | 1.18 | 0.30 | -3.90 | 9.80E-05 | 2.03E-03 |
| *Adgrf5* | 61.28 | 1.18 | 0.40 | -2.94 | 3.23E-03 | 2.80E-02 |
| *H2-T22* | 210.78 | 1.17 | 0.23 | -5.11 | 3.27E-07 | 1.83E-05 |
| *Lat* | 13.87 | 1.17 | 0.43 | -2.72 | 6.52E-03 | 4.64E-02 |
| *Phf11d* | 1369.89 | 1.17 | 0.18 | -6.40 | 1.53E-10 | 2.47E-08 |
| *H2-K2* | 51.92 | 1.17 | 0.33 | -3.50 | 4.60E-04 | 6.75E-03 |
| *Sult2b1* | 58.99 | 1.17 | 0.36 | -3.23 | 1.24E-03 | 1.39E-02 |
| *Mgl2* | 29.15 | 1.17 | 0.32 | -3.61 | 3.05E-04 | 4.92E-03 |
| *Lgals7* | 407.15 | 1.16 | 0.29 | -4.03 | 5.55E-05 | 1.30E-03 |
| *Smad7* | 512.82 | 1.16 | 0.20 | -5.75 | 9.16E-09 | 8.36E-07 |
| *Podxl2* | 83.86 | 1.16 | 0.37 | -3.15 | 1.65E-03 | 1.72E-02 |
| *Pld4* | 57.64 | 1.16 | 0.36 | -3.26 | 1.13E-03 | 1.31E-02 |

| | | | | | | |
|---|---|---|---|---|---|---|
| *Gbp2b* | 1535.39 | 1.15 | 0.23 | -5.12 | 3.13E-07 | 1.76E-05 |
| *A130010J15Rik* | 525.08 | 1.15 | 0.21 | -5.56 | 2.72E-08 | 2.20E-06 |
| *Nfam1* | 24.88 | 1.15 | 0.32 | -3.58 | 3.43E-04 | 5.38E-03 |
| *Epsti1* | 70.00 | 1.15 | 0.26 | -4.43 | 9.57E-06 | 3.04E-04 |
| *Cd5* | 32.94 | 1.15 | 0.29 | -3.97 | 7.05E-05 | 1.57E-03 |
| *Serpina3h* | 101.91 | 1.15 | 0.40 | -2.90 | 3.68E-03 | 3.08E-02 |
| *Glis3* | 120.20 | 1.14 | 0.22 | -5.10 | 3.42E-07 | 1.89E-05 |
| *Gbp3* | 557.69 | 1.14 | 0.29 | -3.99 | 6.69E-05 | 1.51E-03 |
| *C1qc* | 135.50 | 1.14 | 0.32 | -3.54 | 4.08E-04 | 6.15E-03 |
| *Hspa1b* | 14164.12 | 1.14 | 0.19 | -5.99 | 2.06E-09 | 2.24E-07 |
| *Kif5c* | 42.65 | 1.14 | 0.41 | -2.80 | 5.09E-03 | 3.87E-02 |
| *Coro2b* | 85.21 | 1.14 | 0.37 | -3.10 | 1.93E-03 | 1.94E-02 |
| *Taf4b* | 181.68 | 1.13 | 0.36 | -3.14 | 1.69E-03 | 1.75E-02 |
| *Pcdh7* | 1743.30 | 1.13 | 0.19 | -6.00 | 1.98E-09 | 2.19E-07 |
| *Tro* | 23.90 | 1.13 | 0.38 | -2.96 | 3.10E-03 | 2.73E-02 |
| *Gas7* | 80.56 | 1.13 | 0.39 | -2.89 | 3.84E-03 | 3.17E-02 |
| *Tslp* | 89.40 | 1.13 | 0.36 | -3.10 | 1.93E-03 | 1.94E-02 |
| *Cyp7b1* | 55.59 | 1.13 | 0.33 | -3.43 | 6.07E-04 | 8.38E-03 |
| *Tnfsf10* | 367.09 | 1.12 | 0.31 | -3.57 | 3.54E-04 | 5.51E-03 |
| *Lamb3* | 2451.41 | 1.12 | 0.18 | -6.09 | 1.13E-09 | 1.36E-07 |
| *S1pr3* | 18.40 | 1.12 | 0.40 | -2.77 | 5.56E-03 | 4.13E-02 |
| *Wnt3a* | 533.19 | 1.12 | 0.18 | -6.12 | 9.47E-10 | 1.16E-07 |
| *Nrg1* | 643.80 | 1.12 | 0.19 | -5.75 | 9.02E-09 | 8.31E-07 |
| *Gatm* | 59.34 | 1.12 | 0.30 | -3.78 | 1.54E-04 | 2.87E-03 |
| *Wnk4* | 3124.48 | 1.12 | 0.22 | -5.03 | 5.01E-07 | 2.60E-05 |
| *Vsir* | 53.18 | 1.11 | 0.29 | -3.89 | 1.00E-04 | 2.06E-03 |
| *Ercc6l* | 229.18 | 1.11 | 0.20 | -5.47 | 4.61E-08 | 3.54E-06 |
| *Me1* | 325.38 | 1.11 | 0.19 | -5.93 | 3.00E-09 | 3.14E-07 |
| *Gm6548* | 121.36 | 1.10 | 0.23 | -4.75 | 2.06E-06 | 8.50E-05 |
| *P2ry14* | 63.19 | 1.10 | 0.20 | -5.38 | 7.60E-08 | 5.36E-06 |
| *Laptm5* | 189.70 | 1.10 | 0.35 | -3.13 | 1.74E-03 | 1.79E-02 |
| *Crip2* | 470.27 | 1.10 | 0.36 | -3.05 | 2.27E-03 | 2.18E-02 |
| *Ppp1r18* | 591.66 | 1.10 | 0.16 | -6.97 | 3.14E-12 | 7.86E-10 |
| *Dbp* | 2996.15 | 1.10 | 0.30 | -3.70 | 2.13E-04 | 3.72E-03 |
| *Clcf1* | 369.74 | 1.09 | 0.28 | -3.91 | 9.20E-05 | 1.95E-03 |
| *Bdkrb2* | 45.23 | 1.09 | 0.34 | -3.20 | 1.36E-03 | 1.50E-02 |
| *AW011738* | 323.19 | 1.09 | 0.23 | -4.66 | 3.16E-06 | 1.22E-04 |
| *Tent5c* | 116.02 | 1.09 | 0.20 | -5.36 | 8.46E-08 | 5.85E-06 |
| *Btg2* | 1151.83 | 1.09 | 0.36 | -3.03 | 2.43E-03 | 2.28E-02 |

| | | | | | | |
|---|---|---|---|---|---|---|
| *Scamp5* | 185.47 | 1.09 | 0.35 | -3.12 | 1.83E-03 | 1.86E-02 |
| *Gm7592* | 151.14 | 1.09 | 0.23 | -4.77 | 1.84E-06 | 7.89E-05 |
| *Irf7* | 4265.06 | 1.09 | 0.20 | -5.34 | 9.29E-08 | 6.38E-06 |
| *Rrs1* | 566.96 | 1.09 | 0.19 | -5.85 | 4.93E-09 | 4.81E-07 |
| *Tes* | 264.37 | 1.09 | 0.18 | -6.00 | 2.01E-09 | 2.20E-07 |
| *Plk5* | 69.96 | 1.08 | 0.34 | -3.22 | 1.26E-03 | 1.42E-02 |
| *Ptn* | 354.73 | 1.08 | 0.22 | -4.86 | 1.20E-06 | 5.48E-05 |
| *Klf2* | 88.30 | 1.08 | 0.34 | -3.19 | 1.41E-03 | 1.53E-02 |
| *E2f7* | 890.30 | 1.08 | 0.15 | -7.27 | 3.70E-13 | 1.18E-10 |
| *Ncf1* | 33.59 | 1.08 | 0.33 | -3.23 | 1.26E-03 | 1.41E-02 |
| *Inpp5d* | 62.31 | 1.08 | 0.26 | -4.22 | 2.48E-05 | 6.64E-04 |
| *Fes* | 95.89 | 1.08 | 0.28 | -3.84 | 1.24E-04 | 2.42E-03 |
| *Tubb6* | 608.86 | 1.07 | 0.32 | -3.35 | 8.10E-04 | 1.03E-02 |
| *Zeb2* | 66.69 | 1.07 | 0.31 | -3.46 | 5.38E-04 | 7.64E-03 |
| *St3gal4* | 137.94 | 1.07 | 0.29 | -3.66 | 2.51E-04 | 4.24E-03 |
| *Hmx1* | 55.72 | 1.07 | 0.39 | -2.74 | 6.16E-03 | 4.45E-02 |
| *Msn* | 452.50 | 1.07 | 0.20 | -5.45 | 4.92E-08 | 3.76E-06 |
| *Csrnp1* | 1092.54 | 1.07 | 0.20 | -5.40 | 6.66E-08 | 4.83E-06 |
| *H2-T10* | 597.94 | 1.07 | 0.21 | -5.02 | 5.22E-07 | 2.65E-05 |
| *Tmem100* | 34.42 | 1.07 | 0.37 | -2.90 | 3.78E-03 | 3.13E-02 |
| *Tmem8* | 42.41 | 1.06 | 0.33 | -3.24 | 1.21E-03 | 1.37E-02 |
| *Nsg1* | 35.40 | 1.06 | 0.39 | -2.73 | 6.30E-03 | 4.52E-02 |
| *Serpinb10* | 55.16 | 1.06 | 0.34 | -3.08 | 2.10E-03 | 2.05E-02 |
| *Gm18194* | 150.91 | 1.06 | 0.29 | -3.65 | 2.64E-04 | 4.40E-03 |
| *Fgf22* | 29.60 | 1.06 | 0.33 | -3.24 | 1.21E-03 | 1.37E-02 |
| *Serpinb9* | 202.93 | 1.06 | 0.19 | -5.70 | 1.17E-08 | 1.04E-06 |
| *Ovol1* | 1667.75 | 1.06 | 0.21 | -5.11 | 3.14E-07 | 1.76E-05 |
| *Hal* | 66.49 | 1.05 | 0.33 | -3.17 | 1.53E-03 | 1.62E-02 |
| *Mfsd2a* | 165.02 | 1.05 | 0.30 | -3.52 | 4.29E-04 | 6.40E-03 |
| *Il18bp* | 100.25 | 1.05 | 0.30 | -3.50 | 4.69E-04 | 6.84E-03 |
| *Cadps2* | 81.28 | 1.05 | 0.35 | -3.02 | 2.56E-03 | 2.37E-02 |
| *C1qa* | 238.56 | 1.05 | 0.36 | -2.87 | 4.10E-03 | 3.31E-02 |
| *Bambi* | 185.87 | 1.05 | 0.19 | -5.58 | 2.46E-08 | 2.02E-06 |
| *Slc35e4* | 456.71 | 1.04 | 0.31 | -3.39 | 7.03E-04 | 9.30E-03 |
| *Ttc39c* | 55.49 | 1.04 | 0.34 | -3.05 | 2.32E-03 | 2.22E-02 |
| *Nckap5l* | 272.86 | 1.04 | 0.22 | -4.64 | 3.49E-06 | 1.32E-04 |
| *Gm20234* | 839.79 | 1.04 | 0.24 | -4.25 | 2.10E-05 | 5.85E-04 |
| *Cnnm1* | 127.80 | 1.04 | 0.30 | -3.42 | 6.18E-04 | 8.50E-03 |
| *Irf9* | 2295.63 | 1.04 | 0.17 | -6.11 | 9.77E-10 | 1.19E-07 |

| | | | | | | |
|---|---|---|---|---|---|---|
| *Osr2* | 363.67 | 1.03 | 0.36 | -2.84 | 4.54E-03 | 3.57E-02 |
| *Cdc42ep1* | 637.69 | 1.03 | 0.23 | -4.46 | 8.38E-06 | 2.72E-04 |
| *Ramp3* | 27.41 | 1.03 | 0.38 | -2.75 | 5.97E-03 | 4.35E-02 |
| *Celf4* | 96.94 | 1.03 | 0.34 | -3.00 | 2.66E-03 | 2.43E-02 |
| *Cd53* | 44.40 | 1.03 | 0.30 | -3.42 | 6.24E-04 | 8.54E-03 |
| *Mtfr2* | 61.58 | 1.03 | 0.21 | -5.00 | 5.88E-07 | 2.94E-05 |
| *Krt20* | 59.40 | 1.03 | 0.38 | -2.69 | 7.18E-03 | 4.95E-02 |
| *Igkc* | 127.90 | 1.02 | 0.32 | -3.18 | 1.47E-03 | 1.57E-02 |
| *Tfrc* | 643.52 | 1.02 | 0.17 | -5.91 | 3.39E-09 | 3.48E-07 |
| *Slc43a3* | 56.97 | 1.01 | 0.31 | -3.32 | 8.86E-04 | 1.09E-02 |
| *Nexmif* | 28.96 | 1.01 | 0.36 | -2.81 | 4.89E-03 | 3.76E-02 |
| *Egfl7* | 52.02 | 1.01 | 0.35 | -2.86 | 4.30E-03 | 3.43E-02 |
| *Slc25a25* | 659.28 | 1.01 | 0.18 | -5.67 | 1.45E-08 | 1.27E-06 |
| *Pdlim7* | 632.32 | 1.01 | 0.25 | -4.02 | 5.71E-05 | 1.33E-03 |
| *Ccdc88b* | 146.90 | 1.01 | 0.28 | -3.58 | 3.41E-04 | 5.38E-03 |
| *H2-D1* | 15264.45 | 1.01 | 0.18 | -5.73 | 9.84E-09 | 8.94E-07 |
| *Cebpb* | 993.95 | 1.00 | 0.26 | -3.82 | 1.33E-04 | 2.57E-03 |
| *6530402F18Rik* | 43.50 | 1.00 | 0.30 | -3.37 | 7.52E-04 | 9.78E-03 |
| *Bfsp1* | 452.69 | 1.00 | 0.24 | -4.24 | 2.20E-05 | 6.08E-04 |
| *Gm45767* | 86.99 | -1.00 | 0.28 | 3.58 | 3.45E-04 | 5.41E-03 |
| *1110025M09Rik* | 28.32 | -1.00 | 0.34 | 2.95 | 3.19E-03 | 2.78E-02 |
| *Zc3h6* | 317.69 | -1.01 | 0.20 | 5.14 | 2.75E-07 | 1.60E-05 |
| *D530033B14Rik* | 58.24 | -1.01 | 0.24 | 4.24 | 2.20E-05 | 6.08E-04 |
| *Adamtsl2* | 95.86 | -1.01 | 0.31 | 3.27 | 1.08E-03 | 1.27E-02 |
| *0610040B10Rik* | 36.29 | -1.01 | 0.31 | 3.20 | 1.35E-03 | 1.49E-02 |
| *Sorbs2* | 1957.01 | -1.01 | 0.20 | 5.04 | 4.74E-07 | 2.50E-05 |
| *Zfp874a* | 53.49 | -1.01 | 0.26 | 3.95 | 7.70E-05 | 1.68E-03 |
| *Gm15506* | 36.90 | -1.01 | 0.35 | 2.85 | 4.35E-03 | 3.47E-02 |
| *Zfp775* | 59.71 | -1.01 | 0.30 | 3.35 | 8.03E-04 | 1.02E-02 |
| *Tmco6* | 308.26 | -1.01 | 0.16 | 6.21 | 5.34E-10 | 7.14E-08 |
| *Ing4* | 37.67 | -1.02 | 0.28 | 3.58 | 3.42E-04 | 5.38E-03 |
| *Gm14322* | 44.58 | -1.02 | 0.36 | 2.80 | 5.13E-03 | 3.90E-02 |
| *Mroh6* | 752.83 | -1.02 | 0.26 | 3.97 | 7.04E-05 | 1.57E-03 |
| *Zfp874b* | 138.58 | -1.02 | 0.18 | 5.61 | 2.05E-08 | 1.71E-06 |
| *Gm15122* | 43.62 | -1.02 | 0.30 | 3.42 | 6.26E-04 | 8.56E-03 |
| *Col6a3* | 4149.53 | -1.02 | 0.34 | 3.00 | 2.72E-03 | 2.47E-02 |
| *Gxylt2* | 187.41 | -1.02 | 0.26 | 3.95 | 7.86E-05 | 1.70E-03 |
| *Gm16206* | 59.49 | -1.02 | 0.30 | 3.36 | 7.88E-04 | 1.01E-02 |
| *Ccdc9b* | 1204.34 | -1.02 | 0.19 | 5.31 | 1.07E-07 | 7.24E-06 |

| | | | | | | |
|---|---|---|---|---|---|---|
| *Kctd7* | 203.87 | -1.03 | 0.15 | 6.73 | 1.69E-11 | 3.41E-09 |
| *AI197445* | 48.79 | -1.03 | 0.31 | 3.37 | 7.48E-04 | 9.74E-03 |
| *Morn1* | 42.08 | -1.03 | 0.26 | 3.96 | 7.44E-05 | 1.63E-03 |
| *Slc46a2* | 301.49 | -1.03 | 0.24 | 4.25 | 2.16E-05 | 5.99E-04 |
| *Ccdc17* | 281.71 | -1.03 | 0.15 | 6.97 | 3.18E-12 | 7.87E-10 |
| *Cnbd2* | 23.57 | -1.03 | 0.35 | 2.93 | 3.40E-03 | 2.91E-02 |
| *Klhdc9* | 37.90 | -1.04 | 0.31 | 3.30 | 9.54E-04 | 1.15E-02 |
| *Hnmt* | 75.97 | -1.04 | 0.21 | 4.90 | 9.42E-07 | 4.44E-05 |
| *Gm42979* | 155.44 | -1.04 | 0.33 | 3.18 | 1.46E-03 | 1.57E-02 |
| *Gm13648* | 173.97 | -1.04 | 0.27 | 3.92 | 8.96E-05 | 1.90E-03 |
| *Wnt11* | 559.20 | -1.04 | 0.26 | 3.97 | 7.17E-05 | 1.59E-03 |
| *Gm26852* | 40.46 | -1.04 | 0.33 | 3.13 | 1.73E-03 | 1.78E-02 |
| *Slc28a3* | 2858.83 | -1.05 | 0.16 | 6.67 | 2.57E-11 | 5.00E-09 |
| *Mvd* | 1961.49 | -1.05 | 0.21 | 4.99 | 6.18E-07 | 3.08E-05 |
| *Clca4c-ps* | 4861.59 | -1.05 | 0.24 | 4.35 | 1.35E-05 | 4.10E-04 |
| *Ikzf3* | 189.47 | -1.05 | 0.26 | 4.10 | 4.18E-05 | 1.02E-03 |
| *Ttc36* | 295.51 | -1.05 | 0.25 | 4.23 | 2.30E-05 | 6.30E-04 |
| *Gm1976* | 51.24 | -1.05 | 0.32 | 3.25 | 1.15E-03 | 1.32E-02 |
| *Arrdc2* | 204.17 | -1.06 | 0.22 | 4.76 | 1.90E-06 | 8.04E-05 |
| *Gm10814* | 84.42 | -1.06 | 0.34 | 3.14 | 1.69E-03 | 1.75E-02 |
| *Dnajc28* | 95.60 | -1.06 | 0.33 | 3.21 | 1.32E-03 | 1.47E-02 |
| *Mfsd7a* | 299.21 | -1.06 | 0.19 | 5.50 | 3.72E-08 | 2.95E-06 |
| *Gm45606* | 17.51 | -1.06 | 0.38 | 2.82 | 4.85E-03 | 3.74E-02 |
| *Tsc22d3* | 3519.52 | -1.06 | 0.30 | 3.54 | 4.02E-04 | 6.09E-03 |
| *Glis1* | 57.84 | -1.07 | 0.29 | 3.66 | 2.51E-04 | 4.23E-03 |
| *Fam71e1* | 39.00 | -1.07 | 0.36 | 2.94 | 3.28E-03 | 2.83E-02 |
| *Plag1* | 76.34 | -1.07 | 0.26 | 4.13 | 3.69E-05 | 9.22E-04 |
| *2810454H06Rik* | 29.71 | -1.08 | 0.32 | 3.35 | 7.99E-04 | 1.02E-02 |
| *Gm15832* | 31.01 | -1.08 | 0.34 | 3.20 | 1.37E-03 | 1.50E-02 |
| *Rab17* | 315.05 | -1.08 | 0.25 | 4.26 | 2.01E-05 | 5.67E-04 |
| *Pag1* | 498.66 | -1.08 | 0.21 | 5.26 | 1.41E-07 | 8.95E-06 |
| *Lrrn2* | 50.62 | -1.08 | 0.31 | 3.51 | 4.42E-04 | 6.55E-03 |
| *Retreg1* | 524.07 | -1.08 | 0.17 | 6.38 | 1.79E-10 | 2.84E-08 |
| *Thsd1* | 105.33 | -1.09 | 0.27 | 4.05 | 5.07E-05 | 1.20E-03 |
| *Ceacam10* | 101.70 | -1.09 | 0.24 | 4.48 | 7.42E-06 | 2.45E-04 |
| *Tlr5* | 475.77 | -1.09 | 0.24 | 4.45 | 8.46E-06 | 2.73E-04 |
| *Slc25a27* | 249.77 | -1.09 | 0.27 | 3.98 | 6.78E-05 | 1.53E-03 |
| *Zfp239* | 57.65 | -1.10 | 0.35 | 3.13 | 1.73E-03 | 1.78E-02 |
| *Fsbp* | 27.14 | -1.10 | 0.37 | 2.94 | 3.27E-03 | 2.83E-02 |

| | | | | | | |
|---|---|---|---|---|---|---|
| *Gm26704* | 19.73 | -1.10 | 0.38 | 2.89 | 3.80E-03 | 3.14E-02 |
| *Acpp* | 2143.50 | -1.11 | 0.17 | 6.50 | 7.97E-11 | 1.34E-08 |
| *B130024G19Rik* | 72.59 | -1.11 | 0.23 | 4.88 | 1.05E-06 | 4.89E-05 |
| *1190028D05Rik* | 141.47 | -1.12 | 0.29 | 3.88 | 1.03E-04 | 2.11E-03 |
| *1700019D03Rik* | 32.57 | -1.12 | 0.36 | 3.13 | 1.75E-03 | 1.80E-02 |
| *Crct1* | 446.46 | -1.12 | 0.38 | 2.95 | 3.21E-03 | 2.79E-02 |
| *Gpr156* | 21.81 | -1.12 | 0.36 | 3.12 | 1.83E-03 | 1.87E-02 |
| *Gm5475* | 35.20 | -1.13 | 0.39 | 2.90 | 3.73E-03 | 3.10E-02 |
| *Ascl2* | 113.79 | -1.13 | 0.26 | 4.26 | 2.02E-05 | 5.71E-04 |
| *Fmo5* | 491.23 | -1.13 | 0.17 | 6.59 | 4.37E-11 | 8.01E-09 |
| *Gm10791* | 44.17 | -1.14 | 0.38 | 2.99 | 2.80E-03 | 2.53E-02 |
| *Selenbp2* | 20.34 | -1.14 | 0.35 | 3.28 | 1.04E-03 | 1.23E-02 |
| *Parp16* | 363.59 | -1.14 | 0.24 | 4.75 | 1.99E-06 | 8.32E-05 |
| *Gm4813* | 368.98 | -1.14 | 0.19 | 5.94 | 2.86E-09 | 3.03E-07 |
| *A430088P11Rik* | 26.74 | -1.14 | 0.40 | 2.84 | 4.54E-03 | 3.57E-02 |
| *Fcho1* | 422.42 | -1.14 | 0.17 | 6.68 | 2.36E-11 | 4.67E-09 |
| *Cbx7* | 391.64 | -1.14 | 0.16 | 7.02 | 2.29E-12 | 5.98E-10 |
| *Slc8a1* | 25.24 | -1.14 | 0.36 | 3.17 | 1.50E-03 | 1.59E-02 |
| *A4galt* | 200.79 | -1.14 | 0.22 | 5.10 | 3.38E-07 | 1.89E-05 |
| *Acot4* | 37.90 | -1.15 | 0.27 | 4.26 | 2.06E-05 | 5.78E-04 |
| *Izumo4* | 172.25 | -1.15 | 0.21 | 5.53 | 3.13E-08 | 2.51E-06 |
| *Nrxn1* | 178.93 | -1.15 | 0.23 | 5.09 | 3.52E-07 | 1.93E-05 |
| *Zfp169* | 422.97 | -1.16 | 0.25 | 4.70 | 2.59E-06 | 1.04E-04 |
| *Lrrc31* | 79.27 | -1.16 | 0.30 | 3.81 | 1.38E-04 | 2.63E-03 |
| *Myo5c* | 100.66 | -1.16 | 0.29 | 4.06 | 4.89E-05 | 1.16E-03 |
| *Zfp133-ps* | 27.46 | -1.17 | 0.33 | 3.52 | 4.36E-04 | 6.47E-03 |
| *4930550C14Rik* | 47.88 | -1.17 | 0.41 | 2.84 | 4.45E-03 | 3.53E-02 |
| *Abca5* | 672.33 | -1.17 | 0.23 | 5.09 | 3.50E-07 | 1.93E-05 |
| *Tha1* | 109.49 | -1.17 | 0.19 | 6.22 | 4.94E-10 | 6.71E-08 |
| *Gm14508* | 38.95 | -1.18 | 0.33 | 3.60 | 3.18E-04 | 5.08E-03 |
| *Nkapl* | 15.22 | -1.18 | 0.42 | 2.79 | 5.33E-03 | 4.00E-02 |
| *Acrbp* | 51.37 | -1.18 | 0.39 | 3.00 | 2.71E-03 | 2.47E-02 |
| *B230319C09Rik* | 20.80 | -1.19 | 0.43 | 2.76 | 5.77E-03 | 4.24E-02 |
| *Hcrtr1* | 20.02 | -1.19 | 0.40 | 3.00 | 2.69E-03 | 2.46E-02 |
| *Zfp970* | 54.55 | -1.19 | 0.30 | 3.92 | 8.86E-05 | 1.89E-03 |
| *2900005J15Rik* | 34.46 | -1.19 | 0.36 | 3.27 | 1.07E-03 | 1.25E-02 |
| *Gm11423* | 25.68 | -1.19 | 0.39 | 3.06 | 2.23E-03 | 2.15E-02 |
| *Trp53inp1* | 1026.12 | -1.19 | 0.20 | 5.84 | 5.37E-09 | 5.20E-07 |
| *Gm20045* | 59.21 | -1.20 | 0.34 | 3.47 | 5.26E-04 | 7.49E-03 |

| | | | | | | |
|---|---|---|---|---|---|---|
| *Gm17112* | 62.41 | -1.20 | 0.31 | 3.87 | 1.11E-04 | 2.23E-03 |
| *Gm37893* | 74.95 | -1.20 | 0.43 | 2.82 | 4.84E-03 | 3.74E-02 |
| *Gm13204* | 37.20 | -1.20 | 0.42 | 2.88 | 3.97E-03 | 3.24E-02 |
| *Mlph* | 646.82 | -1.20 | 0.25 | 4.91 | 9.20E-07 | 4.36E-05 |
| *Gm15545* | 61.53 | -1.21 | 0.28 | 4.37 | 1.22E-05 | 3.74E-04 |
| *2900076A07Rik* | 40.38 | -1.21 | 0.42 | 2.86 | 4.25E-03 | 3.40E-02 |
| *Gm28497* | 77.85 | -1.21 | 0.26 | 4.65 | 3.30E-06 | 1.27E-04 |
| *4933407K13Rik* | 68.67 | -1.21 | 0.38 | 3.19 | 1.40E-03 | 1.52E-02 |
| *Gm26792* | 31.23 | -1.21 | 0.40 | 3.04 | 2.40E-03 | 2.27E-02 |
| *Kcnk4* | 203.18 | -1.21 | 0.24 | 4.97 | 6.76E-07 | 3.33E-05 |
| *Slc15a1* | 44.99 | -1.21 | 0.31 | 3.95 | 7.83E-05 | 1.70E-03 |
| *Enpep* | 898.84 | -1.22 | 0.22 | 5.56 | 2.63E-08 | 2.15E-06 |
| *Gm43039* | 18.97 | -1.22 | 0.41 | 3.00 | 2.73E-03 | 2.48E-02 |
| *0610031O16Rik* | 93.64 | -1.22 | 0.29 | 4.28 | 1.91E-05 | 5.47E-04 |
| *2810410L24Rik* | 74.76 | -1.23 | 0.25 | 4.88 | 1.06E-06 | 4.90E-05 |
| *Gm17275* | 144.63 | -1.23 | 0.34 | 3.64 | 2.73E-04 | 4.51E-03 |
| *Ugt1a6a* | 70.51 | -1.23 | 0.24 | 5.20 | 1.96E-07 | 1.19E-05 |
| *Zkscan4* | 43.08 | -1.23 | 0.36 | 3.37 | 7.43E-04 | 9.72E-03 |
| *Gm10602* | 25.58 | -1.23 | 0.36 | 3.43 | 6.08E-04 | 8.39E-03 |
| *Best1* | 30.60 | -1.23 | 0.38 | 3.24 | 1.21E-03 | 1.37E-02 |
| *Pou2f3* | 92.25 | -1.24 | 0.20 | 6.12 | 9.18E-10 | 1.14E-07 |
| *Stra6l* | 61.27 | -1.24 | 0.39 | 3.20 | 1.38E-03 | 1.51E-02 |
| *Rnaset2b* | 37.35 | -1.24 | 0.29 | 4.31 | 1.62E-05 | 4.77E-04 |
| *Nap1l3* | 70.39 | -1.25 | 0.22 | 5.78 | 7.27E-09 | 6.86E-07 |
| *Dach1* | 495.49 | -1.25 | 0.35 | 3.54 | 4.01E-04 | 6.08E-03 |
| *Colca2* | 20.46 | -1.25 | 0.35 | 3.60 | 3.13E-04 | 5.01E-03 |
| *Hif3a* | 71.74 | -1.25 | 0.28 | 4.42 | 9.67E-06 | 3.06E-04 |
| *Fdps* | 571.62 | -1.26 | 0.24 | 5.21 | 1.84E-07 | 1.13E-05 |
| *Rbm20* | 195.42 | -1.26 | 0.16 | 8.13 | 4.15E-16 | 2.42E-13 |
| *A830008E24Rik* | 170.97 | -1.26 | 0.22 | 5.67 | 1.46E-08 | 1.27E-06 |
| *Tbx6* | 28.70 | -1.26 | 0.34 | 3.70 | 2.12E-04 | 3.71E-03 |
| *Nckap5* | 39.08 | -1.27 | 0.39 | 3.22 | 1.30E-03 | 1.45E-02 |
| *Gm35339* | 52.45 | -1.27 | 0.33 | 3.81 | 1.37E-04 | 2.63E-03 |
| *Egfl6* | 122.11 | -1.28 | 0.23 | 5.65 | 1.57E-08 | 1.36E-06 |
| *Pigr* | 272.57 | -1.28 | 0.35 | 3.65 | 2.62E-04 | 4.38E-03 |
| *Gm47754* | 95.31 | -1.28 | 0.22 | 5.95 | 2.73E-09 | 2.91E-07 |
| *Gm43071* | 39.88 | -1.29 | 0.43 | 2.98 | 2.91E-03 | 2.60E-02 |
| *Gm14027* | 16.37 | -1.29 | 0.46 | 2.80 | 5.04E-03 | 3.85E-02 |
| *Gm12202* | 100.64 | -1.29 | 0.35 | 3.67 | 2.42E-04 | 4.12E-03 |

| | | | | | | |
|---|---|---|---|---|---|---|
| *Gm30238* | 36.27 | -1.30 | 0.35 | 3.67 | 2.41E-04 | 4.10E-03 |
| *9530034E10Rik* | 26.50 | -1.30 | 0.39 | 3.31 | 9.18E-04 | 1.12E-02 |
| *Zfp879* | 26.63 | -1.30 | 0.43 | 3.06 | 2.20E-03 | 2.13E-02 |
| *Mertk* | 177.50 | -1.31 | 0.25 | 5.30 | 1.18E-07 | 7.90E-06 |
| *Hoxd8* | 31.49 | -1.31 | 0.38 | 3.46 | 5.40E-04 | 7.65E-03 |
| *Gm44065* | 15.65 | -1.31 | 0.49 | 2.70 | 6.92E-03 | 4.83E-02 |
| *Gm43445* | 42.44 | -1.32 | 0.40 | 3.26 | 1.10E-03 | 1.28E-02 |
| *Gm43268* | 14.42 | -1.32 | 0.46 | 2.86 | 4.18E-03 | 3.36E-02 |
| *Cd177* | 38.43 | -1.33 | 0.44 | 2.99 | 2.75E-03 | 2.49E-02 |
| *Gm47827* | 14.92 | -1.34 | 0.46 | 2.91 | 3.61E-03 | 3.03E-02 |
| *Gpr157* | 116.83 | -1.34 | 0.28 | 4.73 | 2.26E-06 | 9.24E-05 |
| *AV356131* | 13.78 | -1.34 | 0.50 | 2.72 | 6.61E-03 | 4.68E-02 |
| *Gm49494* | 12.07 | -1.35 | 0.50 | 2.71 | 6.65E-03 | 4.70E-02 |
| *Lpin1* | 565.68 | -1.35 | 0.19 | 7.26 | 3.78E-13 | 1.19E-10 |
| *Gm17705* | 12.43 | -1.36 | 0.48 | 2.81 | 5.02E-03 | 3.84E-02 |
| *Gm37233* | 34.07 | -1.36 | 0.47 | 2.88 | 3.98E-03 | 3.24E-02 |
| *Grin1os* | 48.15 | -1.36 | 0.27 | 5.10 | 3.41E-07 | 1.89E-05 |
| *Vstm2a* | 35.31 | -1.36 | 0.41 | 3.34 | 8.28E-04 | 1.04E-02 |
| *Rxfp1* | 22.88 | -1.36 | 0.50 | 2.71 | 6.70E-03 | 4.73E-02 |
| *Zpbp2* | 14.51 | -1.37 | 0.46 | 2.97 | 2.95E-03 | 2.62E-02 |
| *Tmem253* | 30.30 | -1.37 | 0.37 | 3.71 | 2.05E-04 | 3.63E-03 |
| *Zfp457* | 22.40 | -1.37 | 0.37 | 3.72 | 2.00E-04 | 3.56E-03 |
| *Gm44321* | 32.51 | -1.37 | 0.47 | 2.95 | 3.20E-03 | 2.78E-02 |
| *4932411K12Rik* | 18.20 | -1.38 | 0.45 | 3.05 | 2.28E-03 | 2.18E-02 |
| *Steap4* | 517.79 | -1.38 | 0.35 | 3.89 | 9.89E-05 | 2.05E-03 |
| *Zfp112* | 39.37 | -1.39 | 0.35 | 3.98 | 6.95E-05 | 1.56E-03 |
| *Gm12258* | 135.74 | -1.40 | 0.31 | 4.47 | 7.76E-06 | 2.54E-04 |
| *Gm16170* | 65.29 | -1.41 | 0.36 | 3.88 | 1.03E-04 | 2.11E-03 |
| *St6galnac1* | 2141.35 | -1.41 | 0.18 | 7.85 | 4.14E-15 | 1.95E-12 |
| *2900052L18Rik* | 56.34 | -1.43 | 0.25 | 5.70 | 1.18E-08 | 1.05E-06 |
| *Gria1* | 69.55 | -1.43 | 0.48 | 2.98 | 2.92E-03 | 2.61E-02 |
| *Gm10069* | 9.33 | -1.44 | 0.52 | 2.76 | 5.79E-03 | 4.25E-02 |
| *Gm37581* | 17.69 | -1.45 | 0.52 | 2.76 | 5.81E-03 | 4.26E-02 |
| *Gm15987* | 305.70 | -1.45 | 0.23 | 6.21 | 5.38E-10 | 7.15E-08 |
| *Gm49502* | 37.98 | -1.46 | 0.53 | 2.76 | 5.86E-03 | 4.28E-02 |
| *Cacna2d3* | 69.42 | -1.46 | 0.51 | 2.83 | 4.61E-03 | 3.60E-02 |
| *Gm45250* | 107.09 | -1.46 | 0.37 | 3.91 | 9.15E-05 | 1.94E-03 |
| *Chkb* | 27.16 | -1.46 | 0.45 | 3.27 | 1.06E-03 | 1.25E-02 |
| *Gm15787* | 55.76 | -1.46 | 0.25 | 5.92 | 3.22E-09 | 3.34E-07 |

| | | | | | | |
|---|---|---|---|---|---|---|
| *Gpc2* | 173.66 | -1.47 | 0.23 | 6.27 | 3.71E-10 | 5.28E-08 |
| *Abhd16b* | 11.24 | -1.47 | 0.52 | 2.85 | 4.36E-03 | 3.47E-02 |
| *Gabrp* | 14.88 | -1.47 | 0.50 | 2.98 | 2.93E-03 | 2.61E-02 |
| *Gm26668* | 27.14 | -1.47 | 0.44 | 3.34 | 8.27E-04 | 1.04E-02 |
| *AC151284.1* | 10.64 | -1.48 | 0.53 | 2.79 | 5.28E-03 | 3.98E-02 |
| *Lce3a* | 883.03 | -1.48 | 0.32 | 4.59 | 4.40E-06 | 1.58E-04 |
| *Upk3b* | 209.94 | -1.49 | 0.27 | 5.48 | 4.17E-08 | 3.25E-06 |
| *D130040H23Rik* | 33.46 | -1.49 | 0.38 | 3.91 | 9.32E-05 | 1.97E-03 |
| *Gm36723* | 13.67 | -1.49 | 0.49 | 3.06 | 2.23E-03 | 2.15E-02 |
| *Gm43609* | 46.75 | -1.49 | 0.39 | 3.82 | 1.35E-04 | 2.60E-03 |
| *Ptprv* | 52.52 | -1.49 | 0.35 | 4.24 | 2.22E-05 | 6.12E-04 |
| *Apoe* | 4582.92 | -1.50 | 0.32 | 4.61 | 3.98E-06 | 1.47E-04 |
| *Cyp4f15* | 385.93 | -1.50 | 0.26 | 5.80 | 6.64E-09 | 6.32E-07 |
| *Gm17276* | 13.08 | -1.50 | 0.52 | 2.88 | 3.96E-03 | 3.24E-02 |
| *Gm38102* | 69.83 | -1.51 | 0.37 | 4.07 | 4.71E-05 | 1.13E-03 |
| *Gm31036* | 10.02 | -1.52 | 0.55 | 2.77 | 5.55E-03 | 4.13E-02 |
| *1700012D14Rik* | 52.60 | -1.52 | 0.23 | 6.55 | 5.67E-11 | 1.00E-08 |
| *Gm6297* | 12.20 | -1.53 | 0.53 | 2.90 | 3.71E-03 | 3.09E-02 |
| *Gm15612* | 13.52 | -1.53 | 0.54 | 2.82 | 4.77E-03 | 3.69E-02 |
| *Fam13a* | 183.61 | -1.55 | 0.23 | 6.86 | 6.79E-12 | 1.47E-09 |
| *Sytl3* | 157.39 | -1.55 | 0.22 | 7.07 | 1.57E-12 | 4.45E-10 |
| *Gm9419* | 8.57 | -1.55 | 0.57 | 2.72 | 6.61E-03 | 4.68E-02 |
| *Zfp108* | 48.96 | -1.56 | 0.29 | 5.36 | 8.17E-08 | 5.70E-06 |
| *Gm11735* | 710.54 | -1.56 | 0.18 | 8.73 | 2.45E-18 | 2.02E-15 |
| *Icam5* | 11.07 | -1.57 | 0.54 | 2.89 | 3.81E-03 | 3.15E-02 |
| *Galnt5* | 32.30 | -1.57 | 0.52 | 3.03 | 2.47E-03 | 2.31E-02 |
| *Kndc1* | 102.71 | -1.57 | 0.29 | 5.44 | 5.22E-08 | 3.96E-06 |
| *Tnf* | 207.09 | -1.57 | 0.34 | 4.57 | 4.77E-06 | 1.69E-04 |
| *Rbpjl* | 83.13 | -1.57 | 0.51 | 3.11 | 1.90E-03 | 1.91E-02 |
| *Fign* | 8.21 | -1.57 | 0.58 | 2.72 | 6.54E-03 | 4.65E-02 |
| *A730015C16Rik* | 16.81 | -1.58 | 0.46 | 3.46 | 5.37E-04 | 7.62E-03 |
| *Ppil6* | 41.89 | -1.58 | 0.45 | 3.53 | 4.08E-04 | 6.15E-03 |
| *G630016G05Rik* | 15.97 | -1.58 | 0.51 | 3.10 | 1.92E-03 | 1.93E-02 |
| *AI182371* | 16.60 | -1.58 | 0.48 | 3.33 | 8.80E-04 | 1.08E-02 |
| *Ccl28* | 77.52 | -1.60 | 0.41 | 3.90 | 9.74E-05 | 2.03E-03 |
| *Bmf* | 314.57 | -1.60 | 0.21 | 7.64 | 2.16E-14 | 9.12E-12 |
| *Dnase1l2* | 99.07 | -1.60 | 0.28 | 5.64 | 1.74E-08 | 1.49E-06 |
| *Ccdc129* | 13.49 | -1.60 | 0.53 | 3.04 | 2.35E-03 | 2.24E-02 |
| *Gm15853* | 10.46 | -1.61 | 0.48 | 3.35 | 8.22E-04 | 1.04E-02 |

| | | | | | | |
|---|---|---|---|---|---|---|
| *Rdh16f2* | 10.98 | -1.61 | 0.52 | 3.10 | 1.91E-03 | 1.92E-02 |
| *Mobp* | 47.38 | -1.62 | 0.35 | 4.62 | 3.76E-06 | 1.41E-04 |
| *Eif4ebp3* | 13.97 | -1.62 | 0.45 | 3.57 | 3.56E-04 | 5.52E-03 |
| *Gm42549* | 43.13 | -1.62 | 0.48 | 3.38 | 7.25E-04 | 9.52E-03 |
| *Gm15947* | 29.00 | -1.62 | 0.33 | 4.92 | 8.55E-07 | 4.08E-05 |
| *Ankdd1a* | 24.01 | -1.62 | 0.48 | 3.39 | 7.00E-04 | 9.27E-03 |
| *Gm5628* | 18.32 | -1.64 | 0.47 | 3.52 | 4.26E-04 | 6.38E-03 |
| *E330011O21Rik* | 10.23 | -1.67 | 0.61 | 2.71 | 6.77E-03 | 4.76E-02 |
| *Mamdc2* | 325.37 | -1.67 | 0.60 | 2.76 | 5.84E-03 | 4.27E-02 |
| *Defb14* | 43.92 | -1.67 | 0.32 | 5.26 | 1.45E-07 | 9.12E-06 |
| *Noxred1* | 10.87 | -1.68 | 0.59 | 2.85 | 4.32E-03 | 3.44E-02 |
| *Gm2670* | 10.49 | -1.68 | 0.53 | 3.18 | 1.46E-03 | 1.56E-02 |
| *Gm43912* | 29.88 | -1.69 | 0.61 | 2.78 | 5.51E-03 | 4.10E-02 |
| *Gm28856* | 18.15 | -1.69 | 0.63 | 2.70 | 6.98E-03 | 4.85E-02 |
| *Ccr10* | 20.26 | -1.69 | 0.40 | 4.19 | 2.74E-05 | 7.23E-04 |
| *Gm13421* | 12.99 | -1.69 | 0.53 | 3.20 | 1.39E-03 | 1.52E-02 |
| *Pabpc4l* | 15.04 | -1.70 | 0.43 | 3.96 | 7.61E-05 | 1.66E-03 |
| *Nkx2-6* | 9.26 | -1.70 | 0.54 | 3.17 | 1.52E-03 | 1.61E-02 |
| *Pantr2* | 32.37 | -1.72 | 0.41 | 4.15 | 3.30E-05 | 8.39E-04 |
| *Gm8250* | 21.55 | -1.73 | 0.40 | 4.29 | 1.80E-05 | 5.19E-04 |
| *Il1f5* | 341.75 | -1.74 | 0.21 | 8.28 | 1.21E-16 | 7.48E-14 |
| *Gm31579* | 11.69 | -1.75 | 0.58 | 3.02 | 2.53E-03 | 2.35E-02 |
| *Col13a1* | 72.59 | -1.75 | 0.57 | 3.07 | 2.16E-03 | 2.09E-02 |
| *Kprp* | 673.24 | -1.76 | 0.33 | 5.40 | 6.79E-08 | 4.87E-06 |
| *Dchs2* | 37.52 | -1.77 | 0.40 | 4.44 | 9.02E-06 | 2.90E-04 |
| *Gm6967* | 9.48 | -1.77 | 0.57 | 3.10 | 1.95E-03 | 1.95E-02 |
| *Gm17249* | 30.65 | -1.77 | 0.47 | 3.74 | 1.86E-04 | 3.36E-03 |
| *Zfp583* | 12.85 | -1.79 | 0.48 | 3.76 | 1.69E-04 | 3.11E-03 |
| *Gm5373* | 9.44 | -1.80 | 0.62 | 2.90 | 3.73E-03 | 3.10E-02 |
| *Unc5c* | 91.89 | -1.80 | 0.35 | 5.18 | 2.18E-07 | 1.30E-05 |
| *5033403F01Rik* | 14.92 | -1.81 | 0.47 | 3.83 | 1.27E-04 | 2.47E-03 |
| *Mfsd6l* | 11.66 | -1.81 | 0.54 | 3.36 | 7.88E-04 | 1.01E-02 |
| *Gm31024* | 33.67 | -1.82 | 0.35 | 5.13 | 2.90E-07 | 1.67E-05 |
| *Gm7967* | 15.99 | -1.82 | 0.59 | 3.09 | 2.03E-03 | 1.99E-02 |
| *Alox5* | 57.41 | -1.82 | 0.36 | 5.03 | 4.87E-07 | 2.55E-05 |
| *AC183097.2* | 43.82 | -1.82 | 0.45 | 4.09 | 4.25E-05 | 1.04E-03 |
| *Fxyd4* | 208.62 | -1.83 | 0.25 | 7.23 | 4.78E-13 | 1.48E-10 |
| *Il1rapl1* | 17.46 | -1.83 | 0.44 | 4.12 | 3.86E-05 | 9.55E-04 |
| *Gm36936* | 59.86 | -1.83 | 0.48 | 3.81 | 1.37E-04 | 2.63E-03 |

| | | | | | | |
|---|---|---|---|---|---|---|
| *Gm36899* | 6.66 | -1.83 | 0.62 | 2.97 | 3.02E-03 | 2.67E-02 |
| *BB218582* | 19.30 | -1.84 | 0.45 | 4.07 | 4.69E-05 | 1.13E-03 |
| *Adrb3* | 52.69 | -1.84 | 0.38 | 4.89 | 1.02E-06 | 4.76E-05 |
| *2010007H06Rik* | 11.20 | -1.84 | 0.61 | 3.03 | 2.46E-03 | 2.31E-02 |
| *Zbtb16* | 533.32 | -1.84 | 0.28 | 6.52 | 6.91E-11 | 1.19E-08 |
| *Platr14* | 15.63 | -1.84 | 0.56 | 3.32 | 8.94E-04 | 1.09E-02 |
| *Tmem132c* | 212.27 | -1.86 | 0.56 | 3.33 | 8.76E-04 | 1.08E-02 |
| *Mir3076* | 12.03 | -1.87 | 0.61 | 3.05 | 2.27E-03 | 2.18E-02 |
| *C730034F03Rik* | 8.60 | -1.88 | 0.62 | 3.04 | 2.38E-03 | 2.25E-02 |
| *4631405J19Rik* | 48.27 | -1.88 | 0.30 | 6.22 | 5.10E-10 | 6.87E-08 |
| *Necab1* | 166.30 | -1.88 | 0.30 | 6.23 | 4.66E-10 | 6.40E-08 |
| *Gm28286* | 61.68 | -1.89 | 0.42 | 4.51 | 6.47E-06 | 2.18E-04 |
| *Gm16046* | 15.47 | -1.89 | 0.67 | 2.85 | 4.40E-03 | 3.49E-02 |
| *Gm15774* | 21.28 | -1.90 | 0.61 | 3.13 | 1.73E-03 | 1.78E-02 |
| *A530001N23Rik* | 65.93 | -1.90 | 0.30 | 6.31 | 2.79E-10 | 4.07E-08 |
| *Gm42572* | 22.58 | -1.92 | 0.62 | 3.10 | 1.97E-03 | 1.96E-02 |
| *Gm18095* | 8.42 | -1.93 | 0.65 | 2.98 | 2.84E-03 | 2.56E-02 |
| *Gm20383* | 91.27 | -1.94 | 0.36 | 5.40 | 6.49E-08 | 4.74E-06 |
| *Gm47753* | 48.67 | -1.95 | 0.49 | 4.01 | 5.97E-05 | 1.37E-03 |
| *Gm20756* | 6.50 | -1.95 | 0.69 | 2.83 | 4.67E-03 | 3.64E-02 |
| *Ntn5* | 6.09 | -1.96 | 0.67 | 2.93 | 3.38E-03 | 2.90E-02 |
| *Gm37274* | 17.78 | -1.96 | 0.68 | 2.86 | 4.23E-03 | 3.39E-02 |
| *Mesp1* | 14.15 | -1.96 | 0.56 | 3.53 | 4.08E-04 | 6.15E-03 |
| *Gm26931* | 12.32 | -1.97 | 0.69 | 2.85 | 4.34E-03 | 3.46E-02 |
| *Mir7001* | 11.59 | -1.97 | 0.65 | 3.02 | 2.51E-03 | 2.33E-02 |
| *Gm11766* | 14.65 | -1.98 | 0.48 | 4.09 | 4.31E-05 | 1.05E-03 |
| *Gm15462* | 10.31 | -1.99 | 0.71 | 2.79 | 5.29E-03 | 3.99E-02 |
| *Fat4* | 595.54 | -1.99 | 0.34 | 5.78 | 7.41E-09 | 6.95E-07 |
| *Il17c* | 17.01 | -2.00 | 0.57 | 3.48 | 4.96E-04 | 7.12E-03 |
| *Spag8* | 10.76 | -2.00 | 0.54 | 3.67 | 2.39E-04 | 4.09E-03 |
| *Hes2* | 9.68 | -2.03 | 0.68 | 2.98 | 2.89E-03 | 2.59E-02 |
| *Lce3f* | 49.47 | -2.05 | 0.49 | 4.15 | 3.31E-05 | 8.41E-04 |
| *Gm38123* | 23.06 | -2.06 | 0.52 | 3.93 | 8.56E-05 | 1.83E-03 |
| *Alkbh3os1* | 10.77 | -2.06 | 0.65 | 3.17 | 1.51E-03 | 1.60E-02 |
| *Gm17029* | 18.97 | -2.07 | 0.57 | 3.63 | 2.79E-04 | 4.60E-03 |
| *Ccdc40* | 5.32 | -2.08 | 0.77 | 2.70 | 7.03E-03 | 4.88E-02 |
| *Gm16106* | 9.79 | -2.09 | 0.63 | 3.30 | 9.61E-04 | 1.16E-02 |
| *Gm17552* | 7.59 | -2.09 | 0.62 | 3.39 | 6.99E-04 | 9.27E-03 |
| *Gm48053* | 11.46 | -2.10 | 0.59 | 3.58 | 3.48E-04 | 5.45E-03 |

| | | | | | | |
|---|---|---|---|---|---|---|
| *Gm29427* | 8.35 | -2.13 | 0.79 | 2.70 | 6.86E-03 | 4.80E-02 |
| *Gm266* | 14.77 | -2.16 | 0.72 | 2.99 | 2.83E-03 | 2.55E-02 |
| *Spats1* | 10.78 | -2.17 | 0.51 | 4.28 | 1.83E-05 | 5.26E-04 |
| *D830024N08Rik* | 13.54 | -2.17 | 0.66 | 3.28 | 1.05E-03 | 1.24E-02 |
| *Gm7162* | 8.57 | -2.17 | 0.78 | 2.80 | 5.14E-03 | 3.90E-02 |
| *Apoc1* | 170.89 | -2.18 | 0.34 | 6.45 | 1.15E-10 | 1.89E-08 |
| *Cyp4a12a* | 583.51 | -2.18 | 0.32 | 6.90 | 5.11E-12 | 1.16E-09 |
| *Gm10101* | 15.31 | -2.18 | 0.47 | 4.67 | 3.04E-06 | 1.18E-04 |
| *Gm32699* | 18.45 | -2.20 | 0.49 | 4.46 | 8.20E-06 | 2.66E-04 |
| *A730091E23Rik* | 20.76 | -2.27 | 0.56 | 4.08 | 4.58E-05 | 1.11E-03 |
| *Gm37249* | 10.23 | -2.27 | 0.75 | 3.01 | 2.59E-03 | 2.39E-02 |
| *Lce3b* | 30.30 | -2.29 | 0.43 | 5.35 | 8.75E-08 | 6.03E-06 |
| *Gm7241* | 5.55 | -2.31 | 0.79 | 2.92 | 3.54E-03 | 3.00E-02 |
| *Cyp2b19* | 275.34 | -2.31 | 0.30 | 7.72 | 1.18E-14 | 5.18E-12 |
| *Gm34121* | 13.04 | -2.32 | 0.55 | 4.19 | 2.84E-05 | 7.43E-04 |
| *Gm13444* | 25.38 | -2.32 | 0.37 | 6.25 | 4.03E-10 | 5.67E-08 |
| *Gm22935* | 19.69 | -2.33 | 0.71 | 3.29 | 9.94E-04 | 1.19E-02 |
| *Odaph* | 5.09 | -2.33 | 0.79 | 2.94 | 3.28E-03 | 2.83E-02 |
| *Fhad1os1* | 11.90 | -2.34 | 0.63 | 3.74 | 1.86E-04 | 3.36E-03 |
| *Gm45470* | 53.73 | -2.39 | 0.47 | 5.08 | 3.75E-07 | 2.04E-05 |
| *Gm45104* | 83.48 | -2.42 | 0.41 | 5.89 | 3.94E-09 | 3.96E-07 |
| *4933437G19Rik* | 5.26 | -2.43 | 0.79 | 3.08 | 2.08E-03 | 2.03E-02 |
| *Gm15512* | 4.77 | -2.45 | 0.85 | 2.87 | 4.14E-03 | 3.34E-02 |
| *Gm39078* | 4.65 | -2.45 | 0.80 | 3.04 | 2.33E-03 | 2.22E-02 |
| *Gm10653* | 6.23 | -2.51 | 0.79 | 3.19 | 1.43E-03 | 1.55E-02 |
| *Gm25219* | 11.07 | -2.51 | 0.73 | 3.43 | 5.99E-04 | 8.31E-03 |
| *Gm5949* | 4.14 | -2.51 | 0.86 | 2.90 | 3.72E-03 | 3.10E-02 |
| *4930511E03Rik* | 12.48 | -2.51 | 0.82 | 3.05 | 2.32E-03 | 2.21E-02 |
| *Gm37226* | 4.80 | -2.51 | 0.91 | 2.78 | 5.49E-03 | 4.09E-02 |
| *Ict1os* | 7.06 | -2.55 | 0.79 | 3.24 | 1.19E-03 | 1.35E-02 |
| *Gm45457* | 15.27 | -2.59 | 0.51 | 5.04 | 4.65E-07 | 2.45E-05 |
| *Gm34294* | 16.65 | -2.60 | 0.59 | 4.39 | 1.13E-05 | 3.51E-04 |
| *Gm43800* | 8.00 | -2.60 | 0.75 | 3.45 | 5.53E-04 | 7.81E-03 |
| *Adam1b* | 8.19 | -2.65 | 0.78 | 3.41 | 6.53E-04 | 8.82E-03 |
| *Gm44202* | 59.03 | -2.66 | 0.47 | 5.61 | 2.04E-08 | 1.71E-06 |
| *Gm48684* | 3.82 | -2.67 | 0.92 | 2.90 | 3.72E-03 | 3.10E-02 |
| *1700008A23Rik* | 9.99 | -2.69 | 0.84 | 3.21 | 1.32E-03 | 1.46E-02 |
| *Ces1h* | 11.44 | -2.69 | 0.62 | 4.32 | 1.54E-05 | 4.56E-04 |
| *Gm10647* | 3.90 | -2.69 | 0.98 | 2.74 | 6.20E-03 | 4.48E-02 |

| | | | | | | |
|---|---|---|---|---|---|---|
| *Gm17182* | 7.07 | -2.70 | 0.81 | 3.35 | 8.14E-04 | 1.03E-02 |
| *Rangrf* | 4.08 | -2.77 | 1.03 | 2.70 | 7.02E-03 | 4.88E-02 |
| *4921534A09Rik* | 6.60 | -2.79 | 0.83 | 3.38 | 7.24E-04 | 9.51E-03 |
| *Gm10244* | 6.72 | -2.81 | 0.87 | 3.23 | 1.24E-03 | 1.40E-02 |
| *Gm35392* | 5.12 | -2.82 | 0.89 | 3.17 | 1.52E-03 | 1.61E-02 |
| *Cxcl2* | 60.82 | -2.83 | 0.53 | 5.37 | 7.77E-08 | 5.45E-06 |
| *Gm10563* | 4.17 | -2.84 | 0.98 | 2.88 | 3.96E-03 | 3.24E-02 |
| *Gm14444* | 3.42 | -2.85 | 1.04 | 2.73 | 6.28E-03 | 4.52E-02 |
| *Plscr5* | 7.73 | -2.85 | 0.75 | 3.81 | 1.37E-04 | 2.63E-03 |
| *AC154667.1* | 21.42 | -2.86 | 0.42 | 6.83 | 8.44E-12 | 1.78E-09 |
| *Gm12996* | 4.38 | -2.89 | 0.95 | 3.04 | 2.39E-03 | 2.26E-02 |
| *Gm45792* | 7.08 | -2.94 | 0.84 | 3.49 | 4.77E-04 | 6.91E-03 |
| *Cnpy1* | 25.29 | -2.96 | 0.49 | 6.02 | 1.72E-09 | 1.93E-07 |
| *3830406C13Rik* | 83.80 | -2.98 | 0.85 | 3.50 | 4.66E-04 | 6.80E-03 |
| *Gm16201* | 3.81 | -3.00 | 0.97 | 3.08 | 2.07E-03 | 2.03E-02 |
| *Gm46495* | 5.74 | -3.05 | 0.90 | 3.39 | 7.11E-04 | 9.38E-03 |
| *Gm31717* | 2.98 | -3.10 | 1.09 | 2.83 | 4.63E-03 | 3.61E-02 |
| *Gp2* | 41.84 | -3.17 | 0.67 | 4.72 | 2.30E-06 | 9.39E-05 |
| *Gm12883* | 10.49 | -3.19 | 0.67 | 4.80 | 1.62E-06 | 7.03E-05 |
| *Gm37818* | 15.17 | -3.23 | 0.67 | 4.82 | 1.41E-06 | 6.20E-05 |
| *Gm25193* | 5.72 | -3.31 | 0.82 | 4.04 | 5.27E-05 | 1.24E-03 |
| *Muc5ac* | 60.32 | -3.34 | 0.87 | 3.83 | 1.27E-04 | 2.46E-03 |
| *Spag6* | 5.05 | -3.46 | 1.07 | 3.22 | 1.29E-03 | 1.44E-02 |
| *Gm10687* | 2.57 | -3.48 | 1.23 | 2.83 | 4.64E-03 | 3.62E-02 |
| *Gm45754* | 3.73 | -3.48 | 1.25 | 2.78 | 5.48E-03 | 4.09E-02 |
| *Gm42628* | 3.80 | -3.49 | 1.19 | 2.93 | 3.41E-03 | 2.91E-02 |
| *Gm14439* | 2.66 | -3.53 | 1.28 | 2.76 | 5.75E-03 | 4.23E-02 |
| *Gm43592* | 2.72 | -3.57 | 1.32 | 2.70 | 6.98E-03 | 4.85E-02 |
| *Bcas1os2* | 4.92 | -3.90 | 1.23 | 3.17 | 1.50E-03 | 1.60E-02 |
| *Gm48591* | 2.24 | -3.91 | 1.44 | 2.71 | 6.77E-03 | 4.76E-02 |
| *Gm43273* | 2.27 | -3.93 | 1.45 | 2.71 | 6.72E-03 | 4.74E-02 |
| *4930412M03Rik* | 2.58 | -4.11 | 1.30 | 3.17 | 1.51E-03 | 1.60E-02 |
| *Gm35572* | 4.21 | -4.25 | 1.27 | 3.35 | 8.00E-04 | 1.02E-02 |
| *Gm23613* | 2.88 | -4.30 | 1.38 | 3.11 | 1.90E-03 | 1.91E-02 |
| *Gm12886* | 4.44 | -4.34 | 1.21 | 3.59 | 3.34E-04 | 5.29E-03 |
| *Gm6592* | 3.23 | -4.44 | 1.33 | 3.34 | 8.53E-04 | 1.06E-02 |
| *Gm20511* | 4.83 | -4.45 | 1.13 | 3.95 | 7.93E-05 | 1.71E-03 |
| *Gm20535* | 3.89 | -4.74 | 1.13 | 4.19 | 2.74E-05 | 7.23E-04 |
| *Mcpt4* | 2.56 | -4.76 | 1.37 | 3.47 | 5.24E-04 | 7.46E-03 |

| | | | | | | |
|---|---|---|---|---|---|---|
| *Gm8520* | 5.97 | -4.76 | 1.29 | 3.70 | 2.19E-04 | 3.80E-03 |
| *Slc25a31* | 6.04 | -4.77 | 1.42 | 3.37 | 7.46E-04 | 9.73E-03 |
| *Tpsab1* | 2.75 | -4.86 | 1.57 | 3.09 | 2.02E-03 | 1.99E-02 |
| *Zfp264* | 8.82 | -6.53 | 2.23 | 2.93 | 3.40E-03 | 2.91E-02 |

**Table S2. Differential gene expression analysis of lacripep- vs PBS-treated *Aire KO* cornea.**

| *Gene* | baseMean | log2FoldChange | lfcSE | stat | pvalue | padj |
|---|---|---|---|---|---|---|
| *Myh3* | 125.52 | 6.45 | 1.53 | -4.23 | 2.38E-05 | 1.19E-03 |
| *Myh7* | 4.37 | 5.87 | 1.56 | -3.76 | 1.67E-04 | 5.64E-03 |
| *Myh4* | 6.63 | 5.87 | 1.48 | -3.96 | 7.52E-05 | 2.93E-03 |
| *Prss56* | 6.34 | 5.80 | 1.17 | -4.96 | 7.00E-07 | 7.99E-05 |
| *Hfe2* | 5.50 | 5.59 | 1.75 | -3.20 | 1.36E-03 | 2.89E-02 |
| *Csrp3* | 10.89 | 5.01 | 1.16 | -4.32 | 1.58E-05 | 8.44E-04 |
| *4930579C12Rik* | 3.22 | 4.80 | 1.37 | -3.49 | 4.74E-04 | 1.28E-02 |
| *Xirp1* | 13.81 | 4.53 | 1.08 | -4.21 | 2.55E-05 | 1.25E-03 |
| *Klhl40* | 6.23 | 4.50 | 1.37 | -3.28 | 1.05E-03 | 2.37E-02 |
| *Glra2* | 9.08 | 4.10 | 0.86 | -4.76 | 1.90E-06 | 1.57E-04 |
| *Gabrr3* | 5.95 | 4.07 | 1.24 | -3.29 | 1.01E-03 | 2.30E-02 |
| *Glb1l3* | 5.88 | 4.06 | 1.06 | -3.83 | 1.27E-04 | 4.55E-03 |
| *Lsmem1* | 12.86 | 3.89 | 1.04 | -3.73 | 1.94E-04 | 6.30E-03 |
| *Hist1h2ah* | 6.17 | 3.81 | 1.18 | -3.23 | 1.23E-03 | 2.66E-02 |
| *Gm26709* | 3.73 | 3.79 | 1.25 | -3.04 | 2.40E-03 | 4.43E-02 |
| *Obscn* | 23.35 | 3.75 | 0.85 | -4.39 | 1.13E-05 | 6.41E-04 |
| *A930041C12Rik* | 5.83 | 3.71 | 1.00 | -3.70 | 2.15E-04 | 6.91E-03 |
| *Ttn* | 58.51 | 3.63 | 0.81 | -4.49 | 6.98E-06 | 4.47E-04 |
| *Scgn* | 35.85 | 3.60 | 0.55 | -6.53 | 6.41E-11 | 5.86E-08 |
| *Gpr75* | 5.32 | 3.54 | 0.99 | -3.58 | 3.40E-04 | 9.99E-03 |
| *Nr2e1* | 14.15 | 3.48 | 0.73 | -4.76 | 1.96E-06 | 1.60E-04 |
| *Cacna1s* | 26.23 | 3.45 | 0.65 | -5.31 | 1.11E-07 | 2.01E-05 |
| *Grm5* | 9.75 | 3.43 | 0.83 | -4.12 | 3.79E-05 | 1.69E-03 |
| *Gp5* | 8.96 | 3.28 | 1.01 | -3.25 | 1.17E-03 | 2.56E-02 |
| *Crxos* | 17.77 | 3.25 | 0.70 | -4.67 | 3.02E-06 | 2.29E-04 |
| *Gm16262* | 15.85 | 3.24 | 0.66 | -4.87 | 1.09E-06 | 1.08E-04 |
| *Hrc* | 14.78 | 3.23 | 0.96 | -3.37 | 7.57E-04 | 1.85E-02 |
| *Fam163a* | 13.73 | 3.21 | 0.65 | -4.91 | 9.18E-07 | 9.55E-05 |
| *RP23-185G9.3* | 15.65 | 3.17 | 0.82 | -3.87 | 1.09E-04 | 4.03E-03 |
| *Gm10742* | 21.03 | 3.16 | 0.71 | -4.44 | 8.94E-06 | 5.37E-04 |
| *Gabra2* | 10.01 | 3.12 | 0.75 | -4.19 | 2.84E-05 | 1.36E-03 |
| *Pnmal1* | 11.26 | 3.11 | 0.73 | -4.25 | 2.15E-05 | 1.08E-03 |
| *Gabrr1* | 54.57 | 3.10 | 0.49 | -6.35 | 2.13E-10 | 1.48E-07 |
| *Ccdc177* | 40.49 | 3.09 | 0.56 | -5.48 | 4.21E-08 | 9.97E-06 |
| *Pde6h* | 50.38 | 3.06 | 0.72 | -4.26 | 2.01E-05 | 1.02E-03 |
| *Cabp2* | 6.33 | 3.06 | 0.89 | -3.43 | 6.00E-04 | 1.53E-02 |
| *Mmp10* | 131.56 | 3.06 | 1.02 | -2.99 | 2.79E-03 | 4.91E-02 |

| | | | | | | |
|---|---|---|---|---|---|---|
| *Galnt13* | 26.60 | 3.04 | 0.73 | -4.19 | 2.79E-05 | 1.35E-03 |
| *Impg1* | 183.37 | 3.04 | 0.45 | -6.75 | 1.49E-11 | 1.72E-08 |
| *Hkdc1* | 16.49 | 3.03 | 0.64 | -4.72 | 2.33E-06 | 1.84E-04 |
| *Gm11961* | 5.46 | 3.03 | 0.97 | -3.13 | 1.75E-03 | 3.44E-02 |
| *Lrit3* | 22.32 | 3.02 | 0.55 | -5.47 | 4.42E-08 | 1.02E-05 |
| *Nlgn1* | 13.63 | 3.00 | 0.67 | -4.48 | 7.55E-06 | 4.75E-04 |
| *Sebox* | 52.25 | 3.00 | 0.75 | -4.01 | 6.02E-05 | 2.45E-03 |
| *Alpk2* | 16.42 | 3.00 | 0.54 | -5.52 | 3.44E-08 | 8.54E-06 |
| *Gngt1* | 510.09 | 2.99 | 0.90 | -3.32 | 9.14E-04 | 2.13E-02 |
| *Rgs9* | 176.77 | 2.99 | 0.53 | -5.66 | 1.49E-08 | 4.71E-06 |
| *Fam163b* | 6.79 | 2.99 | 0.80 | -3.72 | 1.99E-04 | 6.44E-03 |
| *5330413P13Rik* | 9.07 | 2.98 | 0.86 | -3.47 | 5.15E-04 | 1.36E-02 |
| *Adarb2* | 7.45 | 2.96 | 0.87 | -3.40 | 6.66E-04 | 1.66E-02 |
| *A830039N20Ri* | 40.23 | 2.95 | 0.61 | -4.81 | 1.54E-06 | 1.36E-04 |
| *Astn1* | 24.89 | 2.94 | 0.76 | -3.85 | 1.16E-04 | 4.23E-03 |
| *Lrrtm4* | 5.15 | 2.94 | 0.98 | -3.01 | 2.61E-03 | 4.69E-02 |
| *Nrg3* | 5.93 | 2.94 | 0.92 | -3.19 | 1.40E-03 | 2.93E-02 |
| *Cmya5* | 15.98 | 2.93 | 0.85 | -3.44 | 5.92E-04 | 1.52E-02 |
| *Lrit2* | 119.21 | 2.93 | 0.66 | -4.41 | 1.02E-05 | 5.90E-04 |
| *Trpc4* | 5.14 | 2.93 | 0.97 | -3.01 | 2.65E-03 | 4.74E-02 |
| *Ahrr* | 42.21 | 2.92 | 0.55 | -5.29 | 1.25E-07 | 2.20E-05 |
| *Fabp12* | 66.56 | 2.92 | 0.66 | -4.43 | 9.47E-06 | 5.55E-04 |
| *Rtkn2* | 40.52 | 2.92 | 0.47 | -6.20 | 5.57E-10 | 3.12E-07 |
| *Hist1h2ac* | 9.04 | 2.92 | 0.74 | -3.93 | 8.61E-05 | 3.25E-03 |
| *Sall3* | 7.17 | 2.91 | 0.93 | -3.12 | 1.83E-03 | 3.58E-02 |
| *Rd3* | 198.83 | 2.91 | 0.60 | -4.83 | 1.38E-06 | 1.25E-04 |
| *A330050F15Rik* | 10.81 | 2.90 | 0.74 | -3.94 | 8.15E-05 | 3.13E-03 |
| *Zdhhc22* | 10.12 | 2.89 | 0.95 | -3.03 | 2.43E-03 | 4.47E-02 |
| *Grik1* | 45.87 | 2.89 | 0.60 | -4.83 | 1.39E-06 | 1.25E-04 |
| *Pcdh15* | 54.14 | 2.88 | 0.64 | -4.49 | 7.14E-06 | 4.53E-04 |
| *Cartpt* | 72.30 | 2.88 | 0.63 | -4.54 | 5.60E-06 | 3.82E-04 |
| *4930529M08Rik* | 18.94 | 2.87 | 0.57 | -5.08 | 3.79E-07 | 5.14E-05 |
| *Slc4a10* | 62.69 | 2.87 | 0.51 | -5.64 | 1.70E-08 | 5.19E-06 |
| *Rcvrn* | 675.89 | 2.84 | 0.85 | -3.34 | 8.25E-04 | 1.98E-02 |
| *Fam184b* | 12.06 | 2.83 | 0.85 | -3.32 | 9.16E-04 | 2.13E-02 |
| *Nrl* | 1299.55 | 2.82 | 0.77 | -3.67 | 2.40E-04 | 7.53E-03 |
| *Sh2d1a* | 11.50 | 2.82 | 0.82 | -3.44 | 5.73E-04 | 1.49E-02 |
| *Gm17057* | 7.01 | 2.81 | 0.79 | -3.55 | 3.82E-04 | 1.10E-02 |
| *Vgf* | 45.49 | 2.80 | 0.52 | -5.36 | 8.17E-08 | 1.59E-05 |

| | | | | | | |
|---|---|---|---|---|---|---|
| *Chgb* | 216.80 | 2.79 | 0.53 | -5.27 | 1.38E-07 | 2.38E-05 |
| *Ube4bos1* | 12.78 | 2.79 | 0.61 | -4.54 | 5.52E-06 | 3.79E-04 |
| *A530058N18Rik* | 6.11 | 2.78 | 0.90 | -3.09 | 2.00E-03 | 3.83E-02 |
| *Nxnl1* | 185.59 | 2.78 | 0.61 | -4.53 | 5.92E-06 | 3.99E-04 |
| *Slitrk3* | 11.96 | 2.78 | 0.74 | -3.77 | 1.65E-04 | 5.61E-03 |
| *BC033916* | 101.48 | 2.77 | 0.51 | -5.39 | 7.19E-08 | 1.45E-05 |
| *Stk32b* | 5.97 | 2.77 | 0.92 | -2.99 | 2.75E-03 | 4.87E-02 |
| *RP24-95P4.2* | 25.78 | 2.76 | 0.89 | -3.10 | 1.92E-03 | 3.72E-02 |
| *Impg2* | 239.02 | 2.76 | 0.53 | -5.22 | 1.83E-07 | 3.04E-05 |
| *Gabrr2* | 54.19 | 2.76 | 0.42 | -6.54 | 6.33E-11 | 5.86E-08 |
| *Rplp2-ps1* | 24.92 | 2.76 | 0.79 | -3.50 | 4.70E-04 | 1.28E-02 |
| *Igsf21* | 47.77 | 2.76 | 0.56 | -4.95 | 7.45E-07 | 8.35E-05 |
| *Gm11744* | 136.48 | 2.75 | 0.68 | -4.06 | 4.90E-05 | 2.09E-03 |
| *Rp1l1* | 352.39 | 2.75 | 0.54 | -5.12 | 3.06E-07 | 4.40E-05 |
| *A930003A15Rik* | 50.14 | 2.74 | 0.73 | -3.74 | 1.81E-04 | 6.00E-03 |
| *Sox30* | 12.99 | 2.73 | 0.78 | -3.50 | 4.61E-04 | 1.27E-02 |
| *Cplx4* | 304.11 | 2.73 | 0.83 | -3.29 | 1.00E-03 | 2.29E-02 |
| *Rs1* | 1223.17 | 2.73 | 0.56 | -4.87 | 1.11E-06 | 1.08E-04 |
| *Prph2* | 2870.09 | 2.73 | 0.56 | -4.84 | 1.28E-06 | 1.20E-04 |
| *Gpr37* | 124.40 | 2.73 | 0.60 | -4.53 | 5.93E-06 | 3.99E-04 |
| *Hdac9* | 47.25 | 2.72 | 0.45 | -6.00 | 2.03E-09 | 1.01E-06 |
| *Gm13111* | 9.96 | 2.72 | 0.66 | -4.15 | 3.34E-05 | 1.54E-03 |
| *Rp1* | 750.17 | 2.72 | 0.55 | -4.94 | 7.98E-07 | 8.66E-05 |
| *Pde6a* | 1325.16 | 2.72 | 0.51 | -5.34 | 9.29E-08 | 1.75E-05 |
| *Unc80* | 92.70 | 2.72 | 0.53 | -5.16 | 2.48E-07 | 3.82E-05 |
| *Lhfpl4* | 72.47 | 2.72 | 0.61 | -4.43 | 9.45E-06 | 5.55E-04 |
| *Cabp4* | 163.50 | 2.71 | 0.61 | -4.45 | 8.44E-06 | 5.20E-04 |
| *Zfp804a* | 14.70 | 2.71 | 0.67 | -4.07 | 4.76E-05 | 2.04E-03 |
| *A330094K24Rik* | 37.29 | 2.70 | 0.43 | -6.25 | 4.04E-10 | 2.42E-07 |
| *4833424O15Rik* | 12.35 | 2.70 | 0.71 | -3.80 | 1.47E-04 | 5.10E-03 |
| *Pcp2* | 35.27 | 2.70 | 0.59 | -4.54 | 5.61E-06 | 3.82E-04 |
| *Slc24a1* | 785.60 | 2.68 | 0.46 | -5.79 | 7.23E-09 | 2.79E-06 |
| *Scg3* | 81.74 | 2.68 | 0.65 | -4.12 | 3.87E-05 | 1.72E-03 |
| *Sag* | 7505.71 | 2.68 | 0.78 | -3.45 | 5.55E-04 | 1.45E-02 |
| *Isl1* | 48.54 | 2.68 | 0.65 | -4.11 | 3.91E-05 | 1.73E-03 |
| *Cnga1* | 534.67 | 2.68 | 0.58 | -4.59 | 4.47E-06 | 3.19E-04 |
| *Neurod2* | 16.61 | 2.68 | 0.53 | -5.01 | 5.35E-07 | 6.63E-05 |
| *Rho* | 16411.79 | 2.67 | 0.53 | -5.02 | 5.26E-07 | 6.57E-05 |
| *Cngb1* | 578.39 | 2.67 | 0.78 | -3.44 | 5.83E-04 | 1.51E-02 |

| | | | | | | |
|---|---|---|---|---|---|---|
| *Pcdh17* | 18.04 | 2.67 | 0.66 | -4.04 | 5.37E-05 | 2.24E-03 |
| *Tulp1* | 1006.45 | 2.66 | 0.78 | -3.43 | 6.11E-04 | 1.55E-02 |
| *Lin7a* | 103.47 | 2.66 | 0.50 | -5.33 | 1.01E-07 | 1.88E-05 |
| *Cdh8* | 9.33 | 2.66 | 0.74 | -3.57 | 3.56E-04 | 1.04E-02 |
| *Mak* | 120.79 | 2.66 | 0.54 | -4.89 | 9.83E-07 | 9.87E-05 |
| *Nrxn3* | 91.04 | 2.66 | 0.48 | -5.57 | 2.54E-08 | 6.88E-06 |
| *Kcne2* | 31.46 | 2.65 | 0.88 | -3.02 | 2.54E-03 | 4.62E-02 |
| *Tmem26* | 9.85 | 2.65 | 0.72 | -3.69 | 2.26E-04 | 7.18E-03 |
| *Mpp4* | 314.73 | 2.64 | 0.55 | -4.79 | 1.66E-06 | 1.42E-04 |
| *Gnat1* | 6972.68 | 2.64 | 0.55 | -4.80 | 1.59E-06 | 1.39E-04 |
| *Mylpf* | 63.06 | 2.64 | 0.76 | -3.45 | 5.53E-04 | 1.45E-02 |
| *Pde6g* | 1219.85 | 2.64 | 0.80 | -3.31 | 9.41E-04 | 2.18E-02 |
| *Gm7546* | 9.18 | 2.63 | 0.86 | -3.06 | 2.21E-03 | 4.16E-02 |
| *2610034M16Rik* | 214.91 | 2.63 | 0.58 | -4.57 | 4.93E-06 | 3.46E-04 |
| *Hcn1* | 157.16 | 2.63 | 0.55 | -4.79 | 1.70E-06 | 1.43E-04 |
| *Crx* | 741.64 | 2.63 | 0.50 | -5.22 | 1.84E-07 | 3.04E-05 |
| *Gucy2f* | 97.00 | 2.63 | 0.54 | -4.87 | 1.12E-06 | 1.08E-04 |
| *Slco1c1* | 12.77 | 2.63 | 0.70 | -3.77 | 1.62E-04 | 5.54E-03 |
| *Gucy2e* | 511.24 | 2.62 | 0.48 | -5.43 | 5.72E-08 | 1.21E-05 |
| *Gjd2* | 54.29 | 2.62 | 0.61 | -4.29 | 1.80E-05 | 9.34E-04 |
| *Susd3* | 109.59 | 2.62 | 0.41 | -6.36 | 1.97E-10 | 1.48E-07 |
| *Frrs1l* | 23.71 | 2.61 | 0.54 | -4.81 | 1.52E-06 | 1.35E-04 |
| *Pdc* | 1000.62 | 2.61 | 0.77 | -3.39 | 7.03E-04 | 1.74E-02 |
| *Zfp488* | 12.38 | 2.60 | 0.79 | -3.28 | 1.04E-03 | 2.36E-02 |
| *Neb* | 34.37 | 2.60 | 0.80 | -3.26 | 1.11E-03 | 2.48E-02 |
| *Osbp2* | 529.57 | 2.60 | 0.50 | -5.18 | 2.23E-07 | 3.51E-05 |
| *Kcnv2* | 359.95 | 2.59 | 0.52 | -4.95 | 7.53E-07 | 8.38E-05 |
| *5430419D17Rik* | 19.96 | 2.59 | 0.86 | -3.02 | 2.50E-03 | 4.58E-02 |
| *Frmd5* | 64.30 | 2.58 | 0.58 | -4.47 | 7.97E-06 | 4.96E-04 |
| *Lrrn3* | 30.33 | 2.58 | 0.55 | -4.67 | 2.97E-06 | 2.26E-04 |
| *Pde6b* | 1570.79 | 2.58 | 0.54 | -4.75 | 2.08E-06 | 1.68E-04 |
| *Pde6c* | 9.00 | 2.58 | 0.81 | -3.19 | 1.40E-03 | 2.93E-02 |
| *Esrrb* | 357.16 | 2.58 | 0.47 | -5.45 | 5.17E-08 | 1.16E-05 |
| *Prox1* | 125.92 | 2.58 | 0.35 | -7.26 | 3.76E-13 | 9.78E-10 |
| *Vtn* | 543.59 | 2.57 | 0.57 | -4.52 | 6.31E-06 | 4.15E-04 |
| *Bhlhe23* | 21.43 | 2.57 | 0.65 | -3.98 | 6.85E-05 | 2.74E-03 |
| *Rims2* | 97.25 | 2.57 | 0.54 | -4.79 | 1.70E-06 | 1.43E-04 |
| *Elovl4* | 444.66 | 2.56 | 0.34 | -7.57 | 3.70E-14 | 1.61E-10 |
| *Wdr17* | 171.31 | 2.56 | 0.54 | -4.75 | 2.08E-06 | 1.68E-04 |

| | | | | | | |
|---|---|---|---|---|---|---|
| *Gabra1* | 81.44 | 2.56 | 0.57 | -4.52 | 6.25E-06 | 4.13E-04 |
| *Cdk5r2* | 188.53 | 2.56 | 0.56 | -4.61 | 4.06E-06 | 2.93E-04 |
| *Grk1* | 988.75 | 2.56 | 0.51 | -5.04 | 4.57E-07 | 5.92E-05 |
| *Kcnj14* | 356.63 | 2.56 | 0.47 | -5.41 | 6.44E-08 | 1.34E-05 |
| *Dpp6* | 74.33 | 2.56 | 0.53 | -4.85 | 1.25E-06 | 1.18E-04 |
| *Slc1a7* | 118.44 | 2.55 | 0.62 | -4.13 | 3.60E-05 | 1.62E-03 |
| *Dpf3* | 80.45 | 2.55 | 0.52 | -4.87 | 1.13E-06 | 1.08E-04 |
| *Lrit1* | 194.72 | 2.54 | 0.63 | -4.03 | 5.65E-05 | 2.33E-03 |
| *Chst3* | 46.67 | 2.54 | 0.55 | -4.65 | 3.40E-06 | 2.54E-04 |
| *Rtbdn* | 351.57 | 2.54 | 0.55 | -4.64 | 3.56E-06 | 2.63E-04 |
| *Slc17a7* | 1277.11 | 2.54 | 0.76 | -3.33 | 8.72E-04 | 2.06E-02 |
| *Rbp3* | 2985.74 | 2.54 | 0.55 | -4.62 | 3.92E-06 | 2.86E-04 |
| *Akap6* | 59.65 | 2.54 | 0.35 | -7.26 | 3.94E-13 | 9.78E-10 |
| *Csn3* | 136.60 | 2.54 | 0.60 | -4.26 | 2.04E-05 | 1.04E-03 |
| *Lrrc2* | 24.52 | 2.53 | 0.51 | -4.98 | 6.23E-07 | 7.36E-05 |
| *Nrap* | 25.57 | 2.53 | 0.78 | -3.24 | 1.21E-03 | 2.63E-02 |
| *Phactr3* | 23.52 | 2.53 | 0.65 | -3.87 | 1.10E-04 | 4.04E-03 |
| *Car2* | 311.74 | 2.52 | 0.49 | -5.10 | 3.37E-07 | 4.72E-05 |
| *Asic4* | 58.81 | 2.52 | 0.74 | -3.39 | 7.04E-04 | 1.74E-02 |
| *Glp2r* | 31.02 | 2.52 | 0.57 | -4.43 | 9.34E-06 | 5.55E-04 |
| *Aipl1* | 581.36 | 2.52 | 0.56 | -4.53 | 5.95E-06 | 3.99E-04 |
| *Rgs9bp* | 531.70 | 2.52 | 0.51 | -4.91 | 9.24E-07 | 9.55E-05 |
| *Olfm3* | 13.85 | 2.52 | 0.74 | -3.38 | 7.22E-04 | 1.78E-02 |
| *Cacng5* | 27.27 | 2.51 | 0.57 | -4.44 | 9.04E-06 | 5.41E-04 |
| *Lactbl1* | 53.88 | 2.51 | 0.63 | -3.99 | 6.70E-05 | 2.69E-03 |
| *Car8* | 58.43 | 2.51 | 0.52 | -4.84 | 1.27E-06 | 1.20E-04 |
| *Igsf11* | 53.21 | 2.51 | 0.72 | -3.49 | 4.91E-04 | 1.32E-02 |
| *Ankrd33* | 117.69 | 2.50 | 0.61 | -4.12 | 3.76E-05 | 1.68E-03 |
| *Rpe65* | 312.79 | 2.50 | 0.36 | -6.93 | 4.20E-12 | 6.63E-09 |
| *Rlbp1* | 584.96 | 2.50 | 0.38 | -6.56 | 5.43E-11 | 5.54E-08 |
| *Bai3* | 24.32 | 2.50 | 0.58 | -4.31 | 1.62E-05 | 8.55E-04 |
| *Tnfrsf8* | 5.25 | 2.50 | 0.84 | -2.99 | 2.77E-03 | 4.89E-02 |
| *Pcsk1n* | 217.36 | 2.50 | 0.54 | -4.59 | 4.45E-06 | 3.19E-04 |
| *Zfyve28* | 90.02 | 2.50 | 0.51 | -4.89 | 1.01E-06 | 1.01E-04 |
| *AI118078* | 27.26 | 2.50 | 0.70 | -3.59 | 3.35E-04 | 9.92E-03 |
| *Lingo3* | 65.07 | 2.49 | 0.60 | -4.17 | 3.00E-05 | 1.43E-03 |
| *Lamp5* | 36.78 | 2.49 | 0.72 | -3.45 | 5.64E-04 | 1.47E-02 |
| *Paqr9* | 51.73 | 2.48 | 0.52 | -4.77 | 1.81E-06 | 1.51E-04 |
| *Rom1* | 1566.34 | 2.48 | 0.48 | -5.17 | 2.30E-07 | 3.57E-05 |

| | | | | | | |
|---|---|---|---|---|---|---|
| *Hcn4* | 6.32 | 2.47 | 0.80 | -3.10 | 1.91E-03 | 3.71E-02 |
| *Crb1* | 209.34 | 2.47 | 0.56 | -4.38 | 1.18E-05 | 6.62E-04 |
| *Pitpnm2os2* | 37.08 | 2.47 | 0.74 | -3.35 | 8.12E-04 | 1.96E-02 |
| *RP24-247B20.1* | 40.20 | 2.47 | 0.51 | -4.86 | 1.15E-06 | 1.10E-04 |
| *Sox2* | 26.54 | 2.46 | 0.46 | -5.36 | 8.49E-08 | 1.63E-05 |
| *Kcnma1* | 40.20 | 2.46 | 0.48 | -5.10 | 3.40E-07 | 4.72E-05 |
| *Tnnt1* | 26.23 | 2.46 | 0.59 | -4.18 | 2.94E-05 | 1.41E-03 |
| *Prdm8* | 69.16 | 2.45 | 0.49 | -4.98 | 6.39E-07 | 7.50E-05 |
| *Htr3a* | 23.22 | 2.45 | 0.52 | -4.70 | 2.64E-06 | 2.04E-04 |
| *Mmp24* | 15.93 | 2.45 | 0.72 | -3.43 | 6.10E-04 | 1.55E-02 |
| *BC027072* | 348.84 | 2.44 | 0.57 | -4.29 | 1.83E-05 | 9.38E-04 |
| *Rdh12* | 341.32 | 2.44 | 0.49 | -5.01 | 5.46E-07 | 6.73E-05 |
| *Plch1* | 87.11 | 2.43 | 0.51 | -4.81 | 1.47E-06 | 1.32E-04 |
| *Cpne6* | 35.31 | 2.43 | 0.46 | -5.23 | 1.70E-07 | 2.86E-05 |
| *Slc6a11* | 142.32 | 2.43 | 0.50 | -4.85 | 1.23E-06 | 1.17E-04 |
| *Svop* | 58.79 | 2.43 | 0.64 | -3.81 | 1.40E-04 | 4.92E-03 |
| *Cerkl* | 39.61 | 2.43 | 0.49 | -5.00 | 5.84E-07 | 7.15E-05 |
| *Brinp2* | 16.78 | 2.43 | 0.50 | -4.83 | 1.39E-06 | 1.25E-04 |
| *Atp1b2* | 1125.01 | 2.42 | 0.44 | -5.50 | 3.83E-08 | 9.37E-06 |
| *Hist2h4* | 20.51 | 2.42 | 0.61 | -3.98 | 7.01E-05 | 2.77E-03 |
| *St8sia1* | 97.55 | 2.42 | 0.45 | -5.44 | 5.38E-08 | 1.17E-05 |
| *Cacna2d4* | 271.58 | 2.42 | 0.45 | -5.39 | 6.98E-08 | 1.43E-05 |
| *1700009P17Rik* | 20.36 | 2.42 | 0.69 | -3.49 | 4.82E-04 | 1.30E-02 |
| *Abca4* | 924.28 | 2.41 | 0.43 | -5.60 | 2.14E-08 | 5.90E-06 |
| *Gm26954* | 40.99 | 2.41 | 0.59 | -4.08 | 4.43E-05 | 1.93E-03 |
| *Klhl3* | 26.38 | 2.41 | 0.50 | -4.77 | 1.81E-06 | 1.51E-04 |
| *Rorb* | 126.32 | 2.41 | 0.50 | -4.79 | 1.65E-06 | 1.42E-04 |
| *Ankrd33b* | 343.79 | 2.40 | 0.42 | -5.77 | 7.82E-09 | 2.92E-06 |
| *Gabrg2* | 49.23 | 2.40 | 0.51 | -4.72 | 2.39E-06 | 1.87E-04 |
| *Nxnl2* | 62.10 | 2.40 | 0.50 | -4.77 | 1.85E-06 | 1.53E-04 |
| *Guca1b* | 815.15 | 2.40 | 0.56 | -4.31 | 1.65E-05 | 8.65E-04 |
| *Slitrk1* | 8.02 | 2.40 | 0.78 | -3.05 | 2.27E-03 | 4.24E-02 |
| *St6galnac5* | 18.84 | 2.39 | 0.68 | -3.52 | 4.28E-04 | 1.21E-02 |
| *Dusp8* | 217.40 | 2.39 | 0.45 | -5.29 | 1.24E-07 | 2.20E-05 |
| *Wfdc18* | 68.86 | 2.38 | 0.72 | -3.33 | 8.60E-04 | 2.04E-02 |
| *Kcnh6* | 22.24 | 2.38 | 0.69 | -3.45 | 5.63E-04 | 1.47E-02 |
| *Syt1* | 580.44 | 2.38 | 0.46 | -5.13 | 2.89E-07 | 4.26E-05 |
| *Rd3l* | 66.11 | 2.38 | 0.62 | -3.85 | 1.21E-04 | 4.32E-03 |
| *Lrat* | 100.83 | 2.38 | 0.37 | -6.50 | 7.92E-11 | 6.88E-08 |

| | | | | | | |
|---|---|---|---|---|---|---|
| *Dscaml1* | 56.37 | 2.38 | 0.60 | -3.98 | 6.83E-05 | 2.73E-03 |
| *Gabrb3* | 78.97 | 2.38 | 0.52 | -4.56 | 5.14E-06 | 3.59E-04 |
| *Syp* | 345.92 | 2.38 | 0.40 | -5.96 | 2.50E-09 | 1.17E-06 |
| *Mypn* | 13.80 | 2.37 | 0.68 | -3.51 | 4.42E-04 | 1.23E-02 |
| *Bbc3* | 61.36 | 2.37 | 0.63 | -3.75 | 1.77E-04 | 5.88E-03 |
| *Trim67* | 22.91 | 2.37 | 0.59 | -4.01 | 6.00E-05 | 2.45E-03 |
| *Pak7* | 25.13 | 2.37 | 0.71 | -3.33 | 8.61E-04 | 2.04E-02 |
| *Pmaip1* | 55.47 | 2.37 | 0.65 | -3.67 | 2.41E-04 | 7.54E-03 |
| *Map7d2* | 28.85 | 2.37 | 0.74 | -3.21 | 1.34E-03 | 2.85E-02 |
| *Ppef2* | 133.91 | 2.36 | 0.59 | -4.03 | 5.52E-05 | 2.28E-03 |
| *Cacng2* | 15.92 | 2.36 | 0.65 | -3.65 | 2.64E-04 | 8.11E-03 |
| *Tmem178b* | 20.31 | 2.36 | 0.66 | -3.60 | 3.14E-04 | 9.40E-03 |
| *Slc1a2* | 211.64 | 2.36 | 0.27 | -8.87 | 7.63E-19 | 1.33E-14 |
| *Mmd2* | 40.91 | 2.36 | 0.61 | -3.85 | 1.16E-04 | 4.23E-03 |
| *Rgr* | 624.33 | 2.36 | 0.40 | -5.83 | 5.69E-09 | 2.25E-06 |
| *Lrfn2* | 14.04 | 2.35 | 0.62 | -3.80 | 1.47E-04 | 5.10E-03 |
| *Vsx2* | 258.14 | 2.35 | 0.46 | -5.09 | 3.57E-07 | 4.88E-05 |
| *BC030499* | 48.55 | 2.35 | 0.70 | -3.37 | 7.54E-04 | 1.85E-02 |
| *Pcdh10* | 33.72 | 2.35 | 0.51 | -4.64 | 3.48E-06 | 2.60E-04 |
| *Hpcal4* | 62.68 | 2.34 | 0.48 | -4.90 | 9.42E-07 | 9.68E-05 |
| *Tshr* | 22.23 | 2.34 | 0.41 | -5.67 | 1.42E-08 | 4.58E-06 |
| *Cecr2* | 49.46 | 2.34 | 0.40 | -5.84 | 5.38E-09 | 2.17E-06 |
| *Fstl5* | 31.46 | 2.34 | 0.56 | -4.15 | 3.39E-05 | 1.56E-03 |
| *Rdh8* | 65.72 | 2.34 | 0.46 | -5.09 | 3.56E-07 | 4.88E-05 |
| *Gm14290* | 211.54 | 2.34 | 0.46 | -5.13 | 2.89E-07 | 4.26E-05 |
| *Cacna1e* | 7.44 | 2.34 | 0.77 | -3.02 | 2.56E-03 | 4.64E-02 |
| *Ppp2r2b* | 169.43 | 2.34 | 0.52 | -4.50 | 6.95E-06 | 4.47E-04 |
| *Slc38a3* | 455.63 | 2.33 | 0.57 | -4.10 | 4.17E-05 | 1.83E-03 |
| *Plaur* | 203.96 | 2.33 | 0.56 | -4.19 | 2.82E-05 | 1.35E-03 |
| *Nhlh2* | 37.67 | 2.33 | 0.56 | -4.14 | 3.48E-05 | 1.58E-03 |
| *Myt1* | 43.41 | 2.33 | 0.50 | -4.63 | 3.62E-06 | 2.66E-04 |
| *Car10* | 42.24 | 2.32 | 0.49 | -4.70 | 2.62E-06 | 2.03E-04 |
| *Otx2* | 267.28 | 2.32 | 0.46 | -5.06 | 4.26E-07 | 5.65E-05 |
| *Slitrk2* | 36.92 | 2.32 | 0.55 | -4.24 | 2.21E-05 | 1.11E-03 |
| *Skida1* | 111.15 | 2.32 | 0.36 | -6.47 | 1.01E-10 | 8.31E-08 |
| *Elovl2* | 36.04 | 2.32 | 0.65 | -3.56 | 3.68E-04 | 1.07E-02 |
| *Gpr152* | 60.92 | 2.32 | 0.52 | -4.46 | 8.14E-06 | 5.05E-04 |
| *Insm1* | 53.71 | 2.31 | 0.66 | -3.52 | 4.34E-04 | 1.22E-02 |
| *Ppm1e* | 101.71 | 2.31 | 0.47 | -4.91 | 9.10E-07 | 9.55E-05 |

| | | | | | | |
|---|---|---|---|---|---|---|
| *Ackr1* | 53.35 | 2.31 | 0.31 | -7.40 | 1.31E-13 | 4.56E-10 |
| *Gnb3* | 313.81 | 2.30 | 0.58 | -3.93 | 8.43E-05 | 3.20E-03 |
| *Amer2* | 54.46 | 2.30 | 0.41 | -5.64 | 1.69E-08 | 5.19E-06 |
| *Srrm4* | 51.92 | 2.29 | 0.48 | -4.79 | 1.66E-06 | 1.42E-04 |
| *Kcnc4* | 28.30 | 2.29 | 0.67 | -3.43 | 5.97E-04 | 1.52E-02 |
| *Epha7* | 20.73 | 2.29 | 0.60 | -3.80 | 1.47E-04 | 5.10E-03 |
| *Gm15706* | 48.83 | 2.29 | 0.42 | -5.43 | 5.63E-08 | 1.21E-05 |
| *Msi1* | 453.57 | 2.29 | 0.53 | -4.34 | 1.39E-05 | 7.65E-04 |
| *Ttr* | 703.98 | 2.29 | 0.36 | -6.28 | 3.45E-10 | 2.14E-07 |
| *Slc6a13* | 102.13 | 2.29 | 0.72 | -3.15 | 1.62E-03 | 3.24E-02 |
| *Esrrg* | 67.78 | 2.28 | 0.42 | -5.48 | 4.20E-08 | 9.97E-06 |
| *Asphd1* | 11.30 | 2.28 | 0.65 | -3.50 | 4.57E-04 | 1.26E-02 |
| *Aqp4* | 74.99 | 2.28 | 0.51 | -4.45 | 8.73E-06 | 5.30E-04 |
| *Tmem132d* | 27.99 | 2.27 | 0.47 | -4.83 | 1.34E-06 | 1.23E-04 |
| *Rasgrf2* | 96.16 | 2.27 | 0.44 | -5.11 | 3.21E-07 | 4.54E-05 |
| *Kcnj10* | 74.61 | 2.27 | 0.33 | -6.80 | 1.02E-11 | 1.26E-08 |
| *Kcnc1* | 89.52 | 2.26 | 0.52 | -4.36 | 1.29E-05 | 7.13E-04 |
| *Hmga2* | 21.86 | 2.26 | 0.69 | -3.26 | 1.13E-03 | 2.51E-02 |
| *Zfp365* | 186.37 | 2.25 | 0.41 | -5.54 | 2.97E-08 | 7.81E-06 |
| *Slc12a5* | 600.80 | 2.25 | 0.49 | -4.58 | 4.67E-06 | 3.31E-04 |
| *Drd2* | 22.19 | 2.25 | 0.70 | -3.21 | 1.32E-03 | 2.81E-02 |
| *Tmem72* | 21.67 | 2.24 | 0.57 | -3.97 | 7.30E-05 | 2.87E-03 |
| *Unc119* | 1432.67 | 2.24 | 0.40 | -5.63 | 1.78E-08 | 5.29E-06 |
| *Slc6a1* | 321.61 | 2.24 | 0.52 | -4.30 | 1.73E-05 | 8.99E-04 |
| *Rab39b* | 21.45 | 2.24 | 0.67 | -3.35 | 8.19E-04 | 1.97E-02 |
| *Myrip* | 93.28 | 2.24 | 0.26 | -8.71 | 3.00E-18 | 2.60E-14 |
| *Gad1* | 165.76 | 2.23 | 0.57 | -3.89 | 9.91E-05 | 3.67E-03 |
| *Atp1a3* | 1844.09 | 2.23 | 0.45 | -4.94 | 7.98E-07 | 8.66E-05 |
| *Hist1h4j* | 17.76 | 2.22 | 0.62 | -3.59 | 3.27E-04 | 9.75E-03 |
| *Hecw1* | 22.50 | 2.22 | 0.43 | -5.12 | 2.99E-07 | 4.36E-05 |
| *Nyap2* | 30.65 | 2.22 | 0.68 | -3.25 | 1.17E-03 | 2.57E-02 |
| *Snap91* | 255.45 | 2.22 | 0.54 | -4.13 | 3.62E-05 | 1.63E-03 |
| *Sntg2* | 32.34 | 2.21 | 0.45 | -4.89 | 9.84E-07 | 9.87E-05 |
| *Acsl6* | 177.76 | 2.20 | 0.37 | -5.89 | 3.80E-09 | 1.63E-06 |
| *Gria4* | 47.59 | 2.20 | 0.54 | -4.07 | 4.78E-05 | 2.04E-03 |
| *Gm15983* | 15.97 | 2.20 | 0.69 | -3.20 | 1.40E-03 | 2.93E-02 |
| *AI847159* | 87.46 | 2.20 | 0.62 | -3.54 | 4.06E-04 | 1.16E-02 |
| *RP23-52N17.1* | 161.72 | 2.20 | 0.62 | -3.57 | 3.58E-04 | 1.04E-02 |
| *Cbln2* | 31.32 | 2.20 | 0.51 | -4.33 | 1.51E-05 | 8.14E-04 |

| | | | | | | |
|---|---|---|---|---|---|---|
| *Kcnb1* | 230.24 | 2.19 | 0.58 | -3.75 | 1.77E-04 | 5.88E-03 |
| *Zfp385b* | 51.11 | 2.19 | 0.44 | -4.94 | 7.89E-07 | 8.66E-05 |
| *Fam161a* | 152.79 | 2.19 | 0.56 | -3.94 | 8.13E-05 | 3.13E-03 |
| *Mmp17* | 30.73 | 2.19 | 0.44 | -4.99 | 5.91E-07 | 7.18E-05 |
| *Sv2b* | 406.82 | 2.18 | 0.43 | -5.11 | 3.15E-07 | 4.49E-05 |
| *Zfp641* | 23.34 | 2.18 | 0.59 | -3.68 | 2.34E-04 | 7.40E-03 |
| *Samd7* | 103.92 | 2.18 | 0.62 | -3.52 | 4.30E-04 | 1.21E-02 |
| *Sez6* | 76.10 | 2.18 | 0.42 | -5.21 | 1.87E-07 | 3.07E-05 |
| *Ano2* | 68.06 | 2.18 | 0.40 | -5.40 | 6.50E-08 | 1.34E-05 |
| *Prrt3* | 16.85 | 2.18 | 0.47 | -4.66 | 3.23E-06 | 2.43E-04 |
| *Sarm1* | 92.96 | 2.18 | 0.56 | -3.91 | 9.27E-05 | 3.47E-03 |
| *Tmem35* | 45.81 | 2.18 | 0.60 | -3.63 | 2.88E-04 | 8.73E-03 |
| *Tunar* | 17.89 | 2.17 | 0.70 | -3.11 | 1.89E-03 | 3.68E-02 |
| *Cntn4* | 37.22 | 2.17 | 0.61 | -3.56 | 3.67E-04 | 1.07E-02 |
| *Tub* | 279.22 | 2.17 | 0.57 | -3.80 | 1.42E-04 | 4.97E-03 |
| *Wscd1* | 92.12 | 2.17 | 0.44 | -4.93 | 8.31E-07 | 8.85E-05 |
| *1500009C09Rik* | 19.88 | 2.17 | 0.63 | -3.48 | 5.10E-04 | 1.35E-02 |
| *C1ql3* | 37.69 | 2.17 | 0.54 | -4.00 | 6.42E-05 | 2.59E-03 |
| *Rnf165* | 26.03 | 2.17 | 0.58 | -3.71 | 2.05E-04 | 6.61E-03 |
| *Slc16a8* | 187.75 | 2.16 | 0.30 | -7.20 | 5.97E-13 | 1.30E-09 |
| *Scg5* | 191.55 | 2.16 | 0.53 | -4.06 | 4.81E-05 | 2.05E-03 |
| *Kcnj12* | 13.06 | 2.16 | 0.66 | -3.29 | 1.02E-03 | 2.31E-02 |
| *St8sia3* | 20.15 | 2.16 | 0.62 | -3.47 | 5.19E-04 | 1.37E-02 |
| *Tmem59l* | 58.14 | 2.16 | 0.49 | -4.39 | 1.13E-05 | 6.41E-04 |
| *Cacna1h* | 84.60 | 2.15 | 0.55 | -3.95 | 7.89E-05 | 3.05E-03 |
| *Wasf1* | 32.84 | 2.15 | 0.50 | -4.32 | 1.55E-05 | 8.31E-04 |
| *Cspg5* | 187.73 | 2.15 | 0.26 | -8.13 | 4.24E-16 | 2.45E-12 |
| *Mapk8ip2* | 130.53 | 2.15 | 0.53 | -4.06 | 4.93E-05 | 2.09E-03 |
| *Gdap1* | 39.59 | 2.14 | 0.45 | -4.72 | 2.33E-06 | 1.84E-04 |
| *C1ql2* | 60.30 | 2.14 | 0.57 | -3.75 | 1.76E-04 | 5.88E-03 |
| *Dusp26* | 45.97 | 2.14 | 0.54 | -3.93 | 8.32E-05 | 3.17E-03 |
| *Vsx1* | 23.85 | 2.14 | 0.70 | -3.07 | 2.14E-03 | 4.05E-02 |
| *Drd4* | 113.17 | 2.13 | 0.47 | -4.51 | 6.47E-06 | 4.22E-04 |
| *Negr1* | 69.27 | 2.13 | 0.56 | -3.78 | 1.57E-04 | 5.39E-03 |
| *Cadps* | 169.49 | 2.13 | 0.51 | -4.17 | 3.02E-05 | 1.43E-03 |
| *Rpgrip1* | 202.15 | 2.13 | 0.47 | -4.49 | 7.13E-06 | 4.53E-04 |
| *Hist3h2ba* | 31.70 | 2.12 | 0.55 | -3.83 | 1.29E-04 | 4.60E-03 |
| *Pacsin1* | 315.74 | 2.12 | 0.43 | -4.89 | 1.02E-06 | 1.01E-04 |
| *Trpc1* | 137.56 | 2.12 | 0.34 | -6.29 | 3.25E-10 | 2.10E-07 |

| | | | | | | |
|---|---|---|---|---|---|---|
| *Fgf13* | 28.93 | 2.12 | 0.59 | -3.58 | 3.40E-04 | 9.99E-03 |
| *Cacnb2* | 141.93 | 2.12 | 0.50 | -4.21 | 2.51E-05 | 1.23E-03 |
| *Chga* | 141.79 | 2.11 | 0.43 | -4.90 | 9.54E-07 | 9.75E-05 |
| *Gckr* | 88.93 | 2.11 | 0.57 | -3.68 | 2.36E-04 | 7.43E-03 |
| *Ryr2* | 35.82 | 2.11 | 0.47 | -4.51 | 6.46E-06 | 4.22E-04 |
| *Sncb* | 320.09 | 2.11 | 0.42 | -4.99 | 6.13E-07 | 7.33E-05 |
| *Sbsn* | 260.79 | 2.10 | 0.56 | -3.75 | 1.76E-04 | 5.88E-03 |
| *Ptprr* | 11.98 | 2.10 | 0.63 | -3.34 | 8.51E-04 | 2.03E-02 |
| *Cdk5r1* | 77.13 | 2.10 | 0.41 | -5.14 | 2.79E-07 | 4.17E-05 |
| *Caly* | 29.49 | 2.10 | 0.64 | -3.27 | 1.09E-03 | 2.45E-02 |
| *Tmem63c* | 18.60 | 2.10 | 0.69 | -3.03 | 2.44E-03 | 4.49E-02 |
| *Scg2* | 213.95 | 2.10 | 0.43 | -4.93 | 8.14E-07 | 8.78E-05 |
| *Tox2* | 48.36 | 2.09 | 0.33 | -6.29 | 3.26E-10 | 2.10E-07 |
| *Cabp1* | 110.40 | 2.09 | 0.50 | -4.20 | 2.71E-05 | 1.32E-03 |
| *Ildr2* | 68.99 | 2.09 | 0.47 | -4.49 | 7.02E-06 | 4.48E-04 |
| *Slc32a1* | 57.59 | 2.08 | 0.65 | -3.21 | 1.35E-03 | 2.87E-02 |
| *Gpr137c* | 18.60 | 2.08 | 0.50 | -4.20 | 2.67E-05 | 1.31E-03 |
| *Doc2b* | 74.24 | 2.08 | 0.61 | -3.40 | 6.72E-04 | 1.67E-02 |
| *Kctd8* | 20.83 | 2.08 | 0.58 | -3.56 | 3.64E-04 | 1.06E-02 |
| *Impdh1* | 637.35 | 2.08 | 0.40 | -5.20 | 1.98E-07 | 3.18E-05 |
| *Maneal* | 77.72 | 2.07 | 0.49 | -4.22 | 2.41E-05 | 1.19E-03 |
| *Rnf150* | 89.33 | 2.07 | 0.34 | -6.06 | 1.40E-09 | 7.35E-07 |
| *Epha8* | 37.21 | 2.07 | 0.52 | -3.96 | 7.49E-05 | 2.93E-03 |
| *Zfp385c* | 19.43 | 2.06 | 0.64 | -3.22 | 1.27E-03 | 2.73E-02 |
| *Mcf2l* | 184.20 | 2.06 | 0.41 | -5.04 | 4.56E-07 | 5.92E-05 |
| *Mlc1* | 81.77 | 2.06 | 0.46 | -4.45 | 8.67E-06 | 5.30E-04 |
| *B3galt1* | 21.16 | 2.06 | 0.53 | -3.91 | 9.24E-05 | 3.47E-03 |
| *Tbx3* | 23.37 | 2.06 | 0.64 | -3.23 | 1.22E-03 | 2.65E-02 |
| *Gucy1a3* | 135.49 | 2.06 | 0.46 | -4.47 | 7.82E-06 | 4.90E-04 |
| *Krt16* | 3435.08 | 2.06 | 0.46 | -4.47 | 7.93E-06 | 4.95E-04 |
| *Slc4a5* | 479.67 | 2.06 | 0.55 | -3.71 | 2.07E-04 | 6.65E-03 |
| *D3Bwg0562e* | 66.92 | 2.06 | 0.47 | -4.41 | 1.05E-05 | 6.07E-04 |
| *Gnat2* | 83.84 | 2.05 | 0.56 | -3.64 | 2.73E-04 | 8.35E-03 |
| *Fam171b* | 184.43 | 2.05 | 0.36 | -5.72 | 1.06E-08 | 3.60E-06 |
| *Pon1* | 38.90 | 2.05 | 0.35 | -5.77 | 7.91E-09 | 2.92E-06 |
| *Rax* | 92.89 | 2.05 | 0.51 | -3.98 | 6.89E-05 | 2.75E-03 |
| *Amph* | 185.23 | 2.05 | 0.40 | -5.14 | 2.77E-07 | 4.17E-05 |
| *C1ql1* | 88.36 | 2.05 | 0.49 | -4.20 | 2.71E-05 | 1.32E-03 |
| *Nr2e3* | 471.15 | 2.04 | 0.46 | -4.43 | 9.43E-06 | 5.55E-04 |

| | | | | | | |
|---|---|---|---|---|---|---|
| *Faim2* | 177.90 | 2.04 | 0.44 | -4.68 | 2.87E-06 | 2.19E-04 |
| *Srrm3* | 81.50 | 2.04 | 0.56 | -3.68 | 2.36E-04 | 7.42E-03 |
| *Hey2* | 15.34 | 2.04 | 0.57 | -3.61 | 3.08E-04 | 9.26E-03 |
| *Elmod1* | 58.98 | 2.04 | 0.49 | -4.13 | 3.60E-05 | 1.62E-03 |
| *Stxbp1* | 1065.58 | 2.04 | 0.36 | -5.62 | 1.91E-08 | 5.53E-06 |
| *Slc8a3* | 45.54 | 2.04 | 0.37 | -5.52 | 3.30E-08 | 8.31E-06 |
| *Tmem56* | 127.05 | 2.03 | 0.43 | -4.72 | 2.34E-06 | 1.84E-04 |
| *Sgip1* | 155.09 | 2.03 | 0.54 | -3.75 | 1.74E-04 | 5.84E-03 |
| *Fam57b* | 831.76 | 2.03 | 0.47 | -4.31 | 1.60E-05 | 8.49E-04 |
| *Mmp12* | 46.92 | 2.03 | 0.45 | -4.52 | 6.07E-06 | 4.02E-04 |
| *Hapln4* | 14.52 | 2.03 | 0.63 | -3.21 | 1.32E-03 | 2.82E-02 |
| *Tox3* | 26.99 | 2.03 | 0.57 | -3.53 | 4.20E-04 | 1.19E-02 |
| *Celsr3* | 109.71 | 2.02 | 0.36 | -5.63 | 1.80E-08 | 5.29E-06 |
| *Ptprn* | 116.57 | 2.02 | 0.46 | -4.37 | 1.27E-05 | 7.05E-04 |
| *Six3* | 180.84 | 2.02 | 0.54 | -3.72 | 1.97E-04 | 6.40E-03 |
| *Hist2h2bb* | 160.45 | 2.02 | 0.61 | -3.29 | 9.93E-04 | 2.28E-02 |
| *Gramd1b* | 404.46 | 2.02 | 0.35 | -5.76 | 8.39E-09 | 2.97E-06 |
| *Sox8* | 25.88 | 2.02 | 0.42 | -4.81 | 1.54E-06 | 1.36E-04 |
| *Cacng4* | 112.26 | 2.01 | 0.59 | -3.44 | 5.89E-04 | 1.51E-02 |
| *Vat1l* | 216.55 | 2.01 | 0.64 | -3.12 | 1.82E-03 | 3.58E-02 |
| *Ppm1n* | 36.85 | 2.01 | 0.63 | -3.18 | 1.49E-03 | 3.07E-02 |
| *Lrrc4c* | 23.51 | 2.01 | 0.53 | -3.76 | 1.70E-04 | 5.74E-03 |
| *Ap3b2* | 118.01 | 2.01 | 0.39 | -5.18 | 2.24E-07 | 3.51E-05 |
| *Tdrd9* | 18.19 | 2.00 | 0.61 | -3.28 | 1.05E-03 | 2.37E-02 |
| *Strip2* | 54.50 | 2.00 | 0.36 | -5.56 | 2.65E-08 | 7.08E-06 |
| *Rragb* | 55.52 | 2.00 | 0.51 | -3.96 | 7.60E-05 | 2.96E-03 |
| *Myh7b* | 32.18 | 2.00 | 0.48 | -4.17 | 2.99E-05 | 1.43E-03 |
| *Tmem136* | 192.35 | 2.00 | 0.31 | -6.36 | 2.06E-10 | 1.48E-07 |
| *Brsk2* | 132.62 | 2.00 | 0.47 | -4.29 | 1.81E-05 | 9.36E-04 |
| *4930447C04Rik* | 17.82 | 1.99 | 0.56 | -3.55 | 3.88E-04 | 1.11E-02 |
| *Cdh22* | 26.22 | 1.99 | 0.39 | -5.12 | 3.03E-07 | 4.38E-05 |
| *Prmt8* | 18.61 | 1.99 | 0.52 | -3.79 | 1.49E-04 | 5.16E-03 |
| *Cdhr1* | 941.17 | 1.99 | 0.30 | -6.68 | 2.38E-11 | 2.59E-08 |
| *Gucy1b3* | 99.70 | 1.99 | 0.49 | -4.10 | 4.22E-05 | 1.85E-03 |
| *Tmem130* | 84.47 | 1.99 | 0.48 | -4.16 | 3.14E-05 | 1.47E-03 |
| *Ptprt* | 26.04 | 1.98 | 0.49 | -4.06 | 5.00E-05 | 2.12E-03 |
| *St8sia2* | 16.96 | 1.98 | 0.56 | -3.51 | 4.50E-04 | 1.24E-02 |
| *Rasl10b* | 97.18 | 1.97 | 0.39 | -5.05 | 4.44E-07 | 5.85E-05 |
| *Ly6h* | 24.12 | 1.97 | 0.66 | -2.99 | 2.79E-03 | 4.91E-02 |

| | | | | | | |
|---|---|---|---|---|---|---|
| *4-Mar* | 26.04 | 1.97 | 0.49 | -4.05 | 5.17E-05 | 2.17E-03 |
| *Lhx4* | 22.22 | 1.97 | 0.64 | -3.06 | 2.20E-03 | 4.15E-02 |
| *Asic2* | 27.75 | 1.97 | 0.56 | -3.52 | 4.26E-04 | 1.20E-02 |
| *AI593442* | 41.35 | 1.96 | 0.45 | -4.35 | 1.37E-05 | 7.53E-04 |
| *Arhgef26* | 146.89 | 1.96 | 0.37 | -5.28 | 1.29E-07 | 2.24E-05 |
| *Gpr179* | 151.21 | 1.95 | 0.43 | -4.50 | 6.80E-06 | 4.41E-04 |
| *Ptprn2* | 41.92 | 1.95 | 0.49 | -3.95 | 7.89E-05 | 3.05E-03 |
| *Slc22a8* | 41.26 | 1.95 | 0.45 | -4.32 | 1.54E-05 | 8.31E-04 |
| *Ttc25* | 10.14 | 1.95 | 0.63 | -3.10 | 1.94E-03 | 3.74E-02 |
| *Thsd7a* | 55.93 | 1.95 | 0.48 | -4.01 | 5.95E-05 | 2.44E-03 |
| *Ermn* | 24.95 | 1.94 | 0.45 | -4.34 | 1.40E-05 | 7.66E-04 |
| *Mycn* | 10.18 | 1.94 | 0.60 | -3.25 | 1.17E-03 | 2.56E-02 |
| *Eef1a2* | 349.20 | 1.94 | 0.48 | -4.07 | 4.61E-05 | 1.99E-03 |
| *Mfsd7c* | 48.52 | 1.93 | 0.34 | -5.77 | 8.14E-09 | 2.94E-06 |
| *Hist2h3c2* | 17.62 | 1.93 | 0.53 | -3.61 | 3.02E-04 | 9.15E-03 |
| *Abcc8* | 24.53 | 1.93 | 0.56 | -3.43 | 5.93E-04 | 1.52E-02 |
| *Ctnna2* | 30.88 | 1.93 | 0.39 | -4.97 | 6.66E-07 | 7.71E-05 |
| *Rdh5* | 139.73 | 1.93 | 0.41 | -4.72 | 2.33E-06 | 1.84E-04 |
| *Cacng3* | 28.28 | 1.92 | 0.46 | -4.15 | 3.34E-05 | 1.54E-03 |
| *Coro2b* | 244.02 | 1.92 | 0.41 | -4.69 | 2.76E-06 | 2.12E-04 |
| *Ptgs2* | 431.59 | 1.92 | 0.61 | -3.14 | 1.69E-03 | 3.36E-02 |
| *Col2a1* | 83.31 | 1.91 | 0.50 | -3.86 | 1.16E-04 | 4.22E-03 |
| *Epha5* | 22.45 | 1.91 | 0.51 | -3.76 | 1.73E-04 | 5.84E-03 |
| *Hspb6* | 195.22 | 1.91 | 0.35 | -5.44 | 5.21E-08 | 1.16E-05 |
| *Rit2* | 72.22 | 1.91 | 0.57 | -3.33 | 8.80E-04 | 2.08E-02 |
| *Fhod3* | 84.65 | 1.91 | 0.43 | -4.44 | 8.92E-06 | 5.37E-04 |
| *Gpr85* | 17.86 | 1.91 | 0.56 | -3.42 | 6.25E-04 | 1.58E-02 |
| *5330417C22Rik* | 17.36 | 1.90 | 0.54 | -3.54 | 4.06E-04 | 1.16E-02 |
| *Myrf* | 81.54 | 1.90 | 0.30 | -6.42 | 1.39E-10 | 1.09E-07 |
| *Camk2b* | 148.75 | 1.90 | 0.53 | -3.59 | 3.36E-04 | 9.94E-03 |
| *Fam222a* | 51.35 | 1.89 | 0.54 | -3.51 | 4.52E-04 | 1.24E-02 |
| *Atcay* | 65.68 | 1.89 | 0.45 | -4.24 | 2.25E-05 | 1.13E-03 |
| *Abcd2* | 28.47 | 1.89 | 0.58 | -3.27 | 1.07E-03 | 2.40E-02 |
| *Trnp1* | 269.13 | 1.89 | 0.39 | -4.83 | 1.37E-06 | 1.25E-04 |
| *Bex2* | 78.10 | 1.89 | 0.54 | -3.51 | 4.49E-04 | 1.24E-02 |
| *Ppargc1b* | 130.15 | 1.89 | 0.26 | -7.14 | 9.41E-13 | 1.82E-09 |
| *Bex1* | 21.63 | 1.88 | 0.58 | -3.23 | 1.24E-03 | 2.67E-02 |
| *Ubap1l* | 101.17 | 1.88 | 0.41 | -4.55 | 5.25E-06 | 3.65E-04 |
| *Ttyh1* | 305.41 | 1.88 | 0.39 | -4.79 | 1.66E-06 | 1.42E-04 |

| | | | | | | |
|---|---|---|---|---|---|---|
| *Mogat1* | 16.10 | 1.88 | 0.58 | -3.22 | 1.28E-03 | 2.75E-02 |
| *Ndnf* | 75.54 | 1.88 | 0.47 | -4.01 | 6.07E-05 | 2.46E-03 |
| *Fam19a3* | 58.39 | 1.88 | 0.52 | -3.61 | 3.04E-04 | 9.19E-03 |
| *Shank2* | 53.94 | 1.88 | 0.37 | -5.02 | 5.04E-07 | 6.34E-05 |
| *Map1b* | 558.65 | 1.87 | 0.33 | -5.61 | 2.06E-08 | 5.76E-06 |
| *Inmt* | 39.60 | 1.87 | 0.59 | -3.18 | 1.49E-03 | 3.07E-02 |
| *Nrxn2* | 149.07 | 1.87 | 0.52 | -3.59 | 3.28E-04 | 9.75E-03 |
| *Ccdc126* | 240.30 | 1.86 | 0.30 | -6.18 | 6.34E-10 | 3.44E-07 |
| *Snap25* | 815.06 | 1.86 | 0.48 | -3.89 | 9.94E-05 | 3.67E-03 |
| *Rgs8* | 101.90 | 1.86 | 0.53 | -3.51 | 4.51E-04 | 1.24E-02 |
| *Dlg4* | 463.86 | 1.85 | 0.41 | -4.53 | 5.97E-06 | 3.99E-04 |
| *6530402F18Rik* | 97.86 | 1.85 | 0.30 | -6.24 | 4.51E-10 | 2.61E-07 |
| *Neurl1a* | 125.07 | 1.85 | 0.38 | -4.87 | 1.12E-06 | 1.08E-04 |
| *Mef2c* | 97.83 | 1.85 | 0.46 | -4.04 | 5.37E-05 | 2.24E-03 |
| *Hist1h2be* | 82.56 | 1.85 | 0.44 | -4.15 | 3.27E-05 | 1.52E-03 |
| *Bhlhe22* | 42.70 | 1.84 | 0.37 | -5.03 | 4.99E-07 | 6.33E-05 |
| *Abcg4* | 144.88 | 1.84 | 0.43 | -4.31 | 1.61E-05 | 8.51E-04 |
| *Areg* | 105.23 | 1.84 | 0.61 | -3.02 | 2.53E-03 | 4.61E-02 |
| *Cadm3* | 209.39 | 1.84 | 0.51 | -3.61 | 3.08E-04 | 9.26E-03 |
| *Bai1* | 151.06 | 1.84 | 0.31 | -5.89 | 3.85E-09 | 1.63E-06 |
| *Dnajc12* | 44.99 | 1.84 | 0.45 | -4.05 | 5.16E-05 | 2.17E-03 |
| *Scamp5* | 536.22 | 1.83 | 0.45 | -4.09 | 4.38E-05 | 1.91E-03 |
| *Nap1l5* | 251.43 | 1.83 | 0.50 | -3.66 | 2.51E-04 | 7.79E-03 |
| *Cntn1* | 124.60 | 1.83 | 0.43 | -4.26 | 2.02E-05 | 1.03E-03 |
| *Rps6kl1* | 17.76 | 1.83 | 0.52 | -3.49 | 4.78E-04 | 1.29E-02 |
| *Tbc1d9* | 136.56 | 1.83 | 0.41 | -4.44 | 9.11E-06 | 5.44E-04 |
| *Tmem27* | 120.03 | 1.82 | 0.56 | -3.27 | 1.09E-03 | 2.45E-02 |
| *Dusp10* | 312.59 | 1.82 | 0.42 | -4.33 | 1.48E-05 | 8.06E-04 |
| *Dpysl4* | 85.18 | 1.82 | 0.52 | -3.48 | 5.00E-04 | 1.34E-02 |
| *Soga3* | 21.83 | 1.82 | 0.57 | -3.16 | 1.56E-03 | 3.17E-02 |
| *Kcnh2* | 110.87 | 1.82 | 0.47 | -3.91 | 9.31E-05 | 3.47E-03 |
| *Scn1a* | 33.83 | 1.82 | 0.57 | -3.19 | 1.40E-03 | 2.93E-02 |
| *Celf3* | 138.98 | 1.81 | 0.33 | -5.47 | 4.52E-08 | 1.03E-05 |
| *Gabbr2* | 43.67 | 1.81 | 0.44 | -4.14 | 3.53E-05 | 1.60E-03 |
| *Lrp2* | 34.14 | 1.81 | 0.42 | -4.32 | 1.56E-05 | 8.32E-04 |
| *St18* | 17.11 | 1.81 | 0.49 | -3.68 | 2.33E-04 | 7.38E-03 |
| *Chrna4* | 29.41 | 1.81 | 0.49 | -3.68 | 2.34E-04 | 7.40E-03 |
| *Crmp1* | 99.40 | 1.80 | 0.44 | -4.14 | 3.42E-05 | 1.56E-03 |
| *Gprin1* | 44.69 | 1.80 | 0.43 | -4.22 | 2.43E-05 | 1.20E-03 |

| | | | | | | |
|---|---|---|---|---|---|---|
| *Reln* | 289.77 | 1.80 | 0.60 | -3.01 | 2.57E-03 | 4.65E-02 |
| *D630045J12Rik* | 78.69 | 1.79 | 0.33 | -5.38 | 7.41E-08 | 1.48E-05 |
| *Rgs16* | 58.10 | 1.79 | 0.37 | -4.80 | 1.57E-06 | 1.37E-04 |
| *Ccdc85a* | 26.57 | 1.79 | 0.47 | -3.82 | 1.36E-04 | 4.80E-03 |
| *Pde1a* | 37.35 | 1.79 | 0.47 | -3.79 | 1.52E-04 | 5.25E-03 |
| *Samsn1* | 21.83 | 1.79 | 0.47 | -3.81 | 1.40E-04 | 4.92E-03 |
| *Pak3* | 71.90 | 1.79 | 0.38 | -4.75 | 2.03E-06 | 1.66E-04 |
| *Abhd3* | 22.87 | 1.79 | 0.52 | -3.41 | 6.55E-04 | 1.64E-02 |
| *Atp2b2* | 84.64 | 1.78 | 0.31 | -5.74 | 9.62E-09 | 3.34E-06 |
| *Hpca* | 80.87 | 1.78 | 0.54 | -3.30 | 9.68E-04 | 2.23E-02 |
| *Faxc* | 30.97 | 1.78 | 0.45 | -3.93 | 8.47E-05 | 3.21E-03 |
| *Samd11* | 136.21 | 1.78 | 0.42 | -4.24 | 2.22E-05 | 1.11E-03 |
| *Bin1* | 455.69 | 1.77 | 0.32 | -5.48 | 4.25E-08 | 9.97E-06 |
| *Pcbp4* | 795.27 | 1.77 | 0.41 | -4.34 | 1.44E-05 | 7.87E-04 |
| *Trim9* | 106.55 | 1.77 | 0.30 | -5.88 | 4.12E-09 | 1.70E-06 |
| *Napb* | 300.48 | 1.76 | 0.41 | -4.30 | 1.69E-05 | 8.86E-04 |
| *Smpd3* | 27.25 | 1.76 | 0.53 | -3.34 | 8.27E-04 | 1.98E-02 |
| *Nptx1* | 73.96 | 1.76 | 0.40 | -4.43 | 9.41E-06 | 5.55E-04 |
| *Cnr1* | 18.41 | 1.75 | 0.54 | -3.27 | 1.09E-03 | 2.45E-02 |
| *Sez6l* | 28.82 | 1.75 | 0.57 | -3.07 | 2.15E-03 | 4.07E-02 |
| *Pygm* | 266.12 | 1.75 | 0.37 | -4.77 | 1.85E-06 | 1.53E-04 |
| *BC034090* | 208.35 | 1.74 | 0.48 | -3.64 | 2.75E-04 | 8.39E-03 |
| *Arxes1* | 29.17 | 1.74 | 0.42 | -4.20 | 2.72E-05 | 1.32E-03 |
| *Dner* | 74.65 | 1.74 | 0.44 | -3.96 | 7.51E-05 | 2.93E-03 |
| *Sult4a1* | 85.68 | 1.74 | 0.40 | -4.36 | 1.28E-05 | 7.09E-04 |
| *Iqsec3* | 196.98 | 1.74 | 0.44 | -3.95 | 7.87E-05 | 3.05E-03 |
| *Arc* | 223.28 | 1.74 | 0.57 | -3.06 | 2.22E-03 | 4.18E-02 |
| *Zdhhc2* | 184.79 | 1.73 | 0.25 | -7.03 | 2.12E-12 | 3.69E-09 |
| *Plekhb1* | 592.01 | 1.73 | 0.37 | -4.66 | 3.18E-06 | 2.40E-04 |
| *Rundc3b* | 15.16 | 1.73 | 0.58 | -3.01 | 2.62E-03 | 4.70E-02 |
| *Nmnat2* | 72.06 | 1.72 | 0.39 | -4.44 | 8.92E-06 | 5.37E-04 |
| *Pde1b* | 129.04 | 1.72 | 0.53 | -3.25 | 1.15E-03 | 2.54E-02 |
| *Tspyl4* | 229.57 | 1.72 | 0.38 | -4.48 | 7.46E-06 | 4.71E-04 |
| *RP23-90A8.2* | 19.52 | 1.72 | 0.55 | -3.15 | 1.65E-03 | 3.30E-02 |
| *Disp2* | 92.42 | 1.72 | 0.51 | -3.36 | 7.88E-04 | 1.92E-02 |
| *Gnaz* | 86.99 | 1.72 | 0.50 | -3.41 | 6.53E-04 | 1.64E-02 |
| *Nrsn1* | 69.06 | 1.71 | 0.44 | -3.93 | 8.50E-05 | 3.22E-03 |
| *Sez6l2* | 110.04 | 1.71 | 0.42 | -4.08 | 4.49E-05 | 1.95E-03 |
| *Nfasc* | 255.87 | 1.71 | 0.32 | -5.36 | 8.54E-08 | 1.63E-05 |

| | | | | | | |
|---|---|---|---|---|---|---|
| *Lgi1* | 27.75 | 1.71 | 0.56 | -3.06 | 2.25E-03 | 4.21E-02 |
| *Nsg2* | 107.02 | 1.71 | 0.50 | -3.40 | 6.86E-04 | 1.70E-02 |
| *Ntng2* | 121.74 | 1.71 | 0.38 | -4.51 | 6.61E-06 | 4.30E-04 |
| *Tnfrsf23* | 79.42 | 1.71 | 0.54 | -3.16 | 1.57E-03 | 3.17E-02 |
| *Fscn2* | 138.13 | 1.70 | 0.42 | -4.07 | 4.77E-05 | 2.04E-03 |
| *Pcdha10* | 85.34 | 1.70 | 0.41 | -4.16 | 3.18E-05 | 1.48E-03 |
| *Fam196a* | 30.78 | 1.70 | 0.49 | -3.48 | 4.96E-04 | 1.33E-02 |
| *Edil3* | 69.37 | 1.70 | 0.45 | -3.81 | 1.36E-04 | 4.81E-03 |
| *Hspa12a* | 169.23 | 1.70 | 0.40 | -4.22 | 2.45E-05 | 1.21E-03 |
| *Pou3f1* | 16.56 | 1.70 | 0.56 | -3.04 | 2.34E-03 | 4.35E-02 |
| *Vstm2b* | 55.47 | 1.70 | 0.44 | -3.84 | 1.24E-04 | 4.45E-03 |
| *Adcy1* | 323.53 | 1.70 | 0.40 | -4.19 | 2.78E-05 | 1.34E-03 |
| *Syt13* | 73.42 | 1.69 | 0.41 | -4.16 | 3.18E-05 | 1.48E-03 |
| *Rrh* | 45.25 | 1.69 | 0.42 | -4.03 | 5.65E-05 | 2.33E-03 |
| *Amigo2* | 128.44 | 1.69 | 0.31 | -5.53 | 3.19E-08 | 8.25E-06 |
| *Ush2a* | 111.04 | 1.69 | 0.37 | -4.58 | 4.60E-06 | 3.28E-04 |
| *Serpine3* | 31.91 | 1.69 | 0.55 | -3.08 | 2.07E-03 | 3.94E-02 |
| *Gabra3* | 24.05 | 1.69 | 0.55 | -3.07 | 2.12E-03 | 4.03E-02 |
| *Kif1a* | 581.87 | 1.69 | 0.34 | -5.03 | 4.79E-07 | 6.12E-05 |
| *Cyp1b1* | 78.31 | 1.69 | 0.47 | -3.59 | 3.27E-04 | 9.75E-03 |
| *Zswim5* | 41.13 | 1.69 | 0.38 | -4.40 | 1.07E-05 | 6.10E-04 |
| *Ank1* | 45.78 | 1.69 | 0.47 | -3.56 | 3.69E-04 | 1.07E-02 |
| *Rtn4rl2* | 31.21 | 1.69 | 0.49 | -3.42 | 6.28E-04 | 1.59E-02 |
| *Ckb* | 1418.23 | 1.69 | 0.38 | -4.45 | 8.59E-06 | 5.27E-04 |
| *Kcnc3* | 92.55 | 1.68 | 0.34 | -4.95 | 7.42E-07 | 8.35E-05 |
| *Tagln3* | 75.19 | 1.68 | 0.50 | -3.35 | 8.13E-04 | 1.96E-02 |
| *Nim1k* | 78.90 | 1.68 | 0.38 | -4.46 | 8.17E-06 | 5.05E-04 |
| *Reep6* | 1037.61 | 1.68 | 0.36 | -4.72 | 2.33E-06 | 1.84E-04 |
| *Dusp4* | 92.63 | 1.68 | 0.34 | -4.91 | 9.10E-07 | 9.55E-05 |
| *Gng4* | 32.14 | 1.68 | 0.47 | -3.57 | 3.51E-04 | 1.03E-02 |
| *Gnb5* | 573.23 | 1.67 | 0.33 | -5.04 | 4.63E-07 | 5.96E-05 |
| *Mmd* | 197.51 | 1.66 | 0.28 | -5.97 | 2.43E-09 | 1.17E-06 |
| *Lonrf2* | 79.74 | 1.66 | 0.53 | -3.13 | 1.74E-03 | 3.44E-02 |
| *3-Sep* | 101.72 | 1.66 | 0.39 | -4.30 | 1.70E-05 | 8.87E-04 |
| *Efr3b* | 103.55 | 1.66 | 0.50 | -3.34 | 8.52E-04 | 2.03E-02 |
| *Zfp536* | 28.32 | 1.66 | 0.43 | -3.81 | 1.38E-04 | 4.84E-03 |
| *1-Mar* | 53.49 | 1.66 | 0.32 | -5.21 | 1.90E-07 | 3.08E-05 |
| *Adssl1* | 44.22 | 1.66 | 0.49 | -3.41 | 6.48E-04 | 1.63E-02 |
| *Cntnap2* | 71.87 | 1.65 | 0.41 | -3.99 | 6.49E-05 | 2.61E-03 |

| | | | | | | |
|---|---|---|---|---|---|---|
| *Cxxc4* | 69.24 | 1.65 | 0.42 | -3.94 | 8.26E-05 | 3.15E-03 |
| *Kcnab2* | 331.93 | 1.65 | 0.35 | -4.73 | 2.26E-06 | 1.82E-04 |
| *Nefh* | 60.91 | 1.65 | 0.47 | -3.50 | 4.63E-04 | 1.27E-02 |
| *Gm21743* | 222.21 | 1.64 | 0.24 | -6.89 | 5.75E-12 | 8.33E-09 |
| *Synpr* | 197.43 | 1.64 | 0.54 | -3.06 | 2.23E-03 | 4.18E-02 |
| *Diras1* | 58.97 | 1.64 | 0.48 | -3.40 | 6.63E-04 | 1.66E-02 |
| *Phf21b* | 68.32 | 1.64 | 0.41 | -3.97 | 7.07E-05 | 2.79E-03 |
| *Tmod2* | 108.70 | 1.64 | 0.49 | -3.33 | 8.84E-04 | 2.08E-02 |
| *Syngr3* | 37.72 | 1.63 | 0.48 | -3.40 | 6.77E-04 | 1.68E-02 |
| *Podxl* | 295.50 | 1.63 | 0.48 | -3.40 | 6.66E-04 | 1.66E-02 |
| *Fam155a* | 30.88 | 1.63 | 0.47 | -3.48 | 5.10E-04 | 1.35E-02 |
| *Phyhipl* | 145.71 | 1.63 | 0.38 | -4.25 | 2.11E-05 | 1.07E-03 |
| *Ptpro* | 43.21 | 1.63 | 0.36 | -4.53 | 5.84E-06 | 3.96E-04 |
| *Fsd1l* | 84.91 | 1.63 | 0.46 | -3.52 | 4.37E-04 | 1.22E-02 |
| *Cers4* | 248.55 | 1.63 | 0.38 | -4.27 | 1.97E-05 | 1.01E-03 |
| *Eepd1* | 58.18 | 1.63 | 0.33 | -4.93 | 8.21E-07 | 8.80E-05 |
| *Gpr123* | 28.01 | 1.63 | 0.49 | -3.30 | 9.83E-04 | 2.26E-02 |
| *Cdh23* | 19.70 | 1.62 | 0.47 | -3.45 | 5.50E-04 | 1.44E-02 |
| *2900011O08Rik* | 52.55 | 1.62 | 0.52 | -3.09 | 2.02E-03 | 3.87E-02 |
| *Stra6* | 194.25 | 1.62 | 0.31 | -5.30 | 1.13E-07 | 2.03E-05 |
| *Ptp4a3* | 500.61 | 1.62 | 0.27 | -5.96 | 2.57E-09 | 1.17E-06 |
| *Ntm* | 133.14 | 1.61 | 0.52 | -3.11 | 1.90E-03 | 3.68E-02 |
| *Rtn4rl1* | 112.75 | 1.61 | 0.36 | -4.45 | 8.71E-06 | 5.30E-04 |
| *Bsn* | 100.29 | 1.61 | 0.42 | -3.85 | 1.18E-04 | 4.28E-03 |
| *Kif5c* | 160.60 | 1.61 | 0.35 | -4.55 | 5.33E-06 | 3.69E-04 |
| *4-Sep* | 166.61 | 1.60 | 0.32 | -5.07 | 3.96E-07 | 5.31E-05 |
| *Atp7b* | 32.09 | 1.60 | 0.46 | -3.50 | 4.63E-04 | 1.27E-02 |
| *Nap1l2* | 34.35 | 1.60 | 0.39 | -4.05 | 5.03E-05 | 2.12E-03 |
| *Tmem108* | 67.56 | 1.59 | 0.32 | -4.91 | 9.19E-07 | 9.55E-05 |
| *B4galnt4* | 105.20 | 1.59 | 0.47 | -3.39 | 6.96E-04 | 1.72E-02 |
| *Slc16a6* | 169.09 | 1.59 | 0.32 | -4.95 | 7.24E-07 | 8.22E-05 |
| *Fam78b* | 46.83 | 1.59 | 0.41 | -3.91 | 9.11E-05 | 3.43E-03 |
| *Rtn1* | 247.48 | 1.59 | 0.35 | -4.55 | 5.35E-06 | 3.69E-04 |
| *Rph3a* | 143.53 | 1.58 | 0.42 | -3.80 | 1.47E-04 | 5.10E-03 |
| *Atp8a2* | 43.63 | 1.58 | 0.45 | -3.50 | 4.66E-04 | 1.27E-02 |
| *Gria2* | 85.51 | 1.58 | 0.49 | -3.22 | 1.28E-03 | 2.75E-02 |
| *Zcchc18* | 101.09 | 1.58 | 0.40 | -3.94 | 8.14E-05 | 3.13E-03 |
| *Bcat1* | 179.81 | 1.58 | 0.50 | -3.18 | 1.45E-03 | 3.01E-02 |
| *Nacad* | 62.54 | 1.58 | 0.45 | -3.55 | 3.92E-04 | 1.12E-02 |

| | | | | | | |
|---|---|---|---|---|---|---|
| *Ppp1r9a* | 103.16 | 1.58 | 0.48 | -3.29 | 9.96E-04 | 2.28E-02 |
| *Fam167a* | 35.05 | 1.58 | 0.48 | -3.31 | 9.49E-04 | 2.20E-02 |
| *Mamld1* | 49.67 | 1.58 | 0.49 | -3.19 | 1.42E-03 | 2.96E-02 |
| *Gng3* | 86.42 | 1.58 | 0.50 | -3.16 | 1.57E-03 | 3.18E-02 |
| *Nptx2* | 34.32 | 1.58 | 0.45 | -3.51 | 4.47E-04 | 1.24E-02 |
| *Zmat4* | 33.09 | 1.58 | 0.43 | -3.66 | 2.51E-04 | 7.79E-03 |
| *Sec14l2* | 68.38 | 1.58 | 0.33 | -4.71 | 2.50E-06 | 1.95E-04 |
| *Slc25a33* | 80.07 | 1.58 | 0.36 | -4.42 | 9.98E-06 | 5.82E-04 |
| *Elfn2* | 29.36 | 1.57 | 0.49 | -3.20 | 1.39E-03 | 2.92E-02 |
| *Ankrd34a* | 26.38 | 1.57 | 0.52 | -3.00 | 2.72E-03 | 4.84E-02 |
| *Tdrkh* | 36.82 | 1.57 | 0.48 | -3.25 | 1.15E-03 | 2.54E-02 |
| *Sema7a* | 133.42 | 1.57 | 0.43 | -3.65 | 2.61E-04 | 8.02E-03 |
| *Gfap* | 36.63 | 1.57 | 0.45 | -3.46 | 5.47E-04 | 1.43E-02 |
| *Add2* | 19.15 | 1.57 | 0.48 | -3.26 | 1.12E-03 | 2.50E-02 |
| *Slc7a11* | 83.27 | 1.56 | 0.45 | -3.44 | 5.85E-04 | 1.51E-02 |
| *Ebpl* | 180.47 | 1.56 | 0.32 | -4.83 | 1.35E-06 | 1.24E-04 |
| *Ttc39c* | 83.48 | 1.56 | 0.35 | -4.42 | 9.73E-06 | 5.69E-04 |
| *Dlgap1* | 65.61 | 1.56 | 0.40 | -3.94 | 8.17E-05 | 3.13E-03 |
| *Eml5* | 161.35 | 1.56 | 0.31 | -4.99 | 6.16E-07 | 7.33E-05 |
| *Itga4* | 111.51 | 1.55 | 0.37 | -4.23 | 2.29E-05 | 1.14E-03 |
| *Diras2* | 235.80 | 1.55 | 0.35 | -4.49 | 6.96E-06 | 4.47E-04 |
| *Usp13* | 29.68 | 1.55 | 0.44 | -3.50 | 4.73E-04 | 1.28E-02 |
| *Gas7* | 240.36 | 1.55 | 0.41 | -3.75 | 1.75E-04 | 5.88E-03 |
| *Jph1* | 17.10 | 1.55 | 0.48 | -3.19 | 1.40E-03 | 2.93E-02 |
| *Pde10a* | 43.20 | 1.54 | 0.41 | -3.74 | 1.83E-04 | 6.04E-03 |
| *Dpysl5* | 48.59 | 1.54 | 0.47 | -3.25 | 1.17E-03 | 2.56E-02 |
| *Reep2* | 85.28 | 1.53 | 0.42 | -3.65 | 2.58E-04 | 7.97E-03 |
| *Kcnq2* | 74.82 | 1.53 | 0.43 | -3.60 | 3.22E-04 | 9.62E-03 |
| *Mtmr7* | 59.78 | 1.53 | 0.49 | -3.11 | 1.85E-03 | 3.61E-02 |
| *Psd* | 462.98 | 1.53 | 0.30 | -5.14 | 2.76E-07 | 4.17E-05 |
| *Pde8b* | 75.81 | 1.53 | 0.37 | -4.12 | 3.84E-05 | 1.71E-03 |
| *Nrcam* | 68.22 | 1.53 | 0.42 | -3.63 | 2.86E-04 | 8.68E-03 |
| *Ccl6* | 62.24 | 1.52 | 0.47 | -3.23 | 1.23E-03 | 2.66E-02 |
| *Rnf157* | 179.77 | 1.52 | 0.31 | -4.87 | 1.11E-06 | 1.08E-04 |
| *Kif21b* | 313.86 | 1.52 | 0.28 | -5.38 | 7.48E-08 | 1.48E-05 |
| *Mpp3* | 45.91 | 1.52 | 0.46 | -3.32 | 8.86E-04 | 2.08E-02 |
| *Plcxd2* | 177.15 | 1.51 | 0.29 | -5.31 | 1.11E-07 | 2.01E-05 |
| *Nol4l* | 118.73 | 1.51 | 0.27 | -5.67 | 1.39E-08 | 4.55E-06 |
| *Zfp385a* | 1168.47 | 1.51 | 0.27 | -5.62 | 1.94E-08 | 5.53E-06 |

| | | | | | | |
|---|---|---|---|---|---|---|
| *Sema6a* | 53.62 | 1.51 | 0.44 | -3.45 | 5.55E-04 | 1.45E-02 |
| *Astn2* | 38.65 | 1.51 | 0.48 | -3.13 | 1.72E-03 | 3.41E-02 |
| *Pcdh9* | 20.53 | 1.50 | 0.43 | -3.52 | 4.27E-04 | 1.20E-02 |
| *Nol4* | 24.46 | 1.50 | 0.48 | -3.16 | 1.60E-03 | 3.22E-02 |
| *Slc35g2* | 26.70 | 1.50 | 0.50 | -3.01 | 2.61E-03 | 4.69E-02 |
| *Kcnab3* | 45.02 | 1.50 | 0.48 | -3.13 | 1.74E-03 | 3.44E-02 |
| *Atp6v1g2* | 96.65 | 1.50 | 0.48 | -3.12 | 1.84E-03 | 3.60E-02 |
| *Elfn1* | 142.99 | 1.50 | 0.41 | -3.69 | 2.22E-04 | 7.08E-03 |
| *Rims1* | 60.69 | 1.50 | 0.41 | -3.64 | 2.76E-04 | 8.42E-03 |
| *Tmem151b* | 17.73 | 1.49 | 0.49 | -3.06 | 2.20E-03 | 4.15E-02 |
| *Lrp11* | 116.79 | 1.49 | 0.30 | -4.99 | 6.16E-07 | 7.33E-05 |
| *Ppp2r2c* | 67.81 | 1.49 | 0.36 | -4.15 | 3.31E-05 | 1.53E-03 |
| *Gpm6a* | 454.17 | 1.49 | 0.36 | -4.14 | 3.41E-05 | 1.56E-03 |
| *Gpr162* | 164.22 | 1.49 | 0.41 | -3.64 | 2.70E-04 | 8.29E-03 |
| *Celf4* | 202.21 | 1.49 | 0.29 | -5.07 | 3.97E-07 | 5.31E-05 |
| *Nsg1* | 80.51 | 1.49 | 0.37 | -4.08 | 4.55E-05 | 1.97E-03 |
| *Cacna1b* | 19.85 | 1.49 | 0.49 | -3.02 | 2.55E-03 | 4.63E-02 |
| *Gulp1* | 57.65 | 1.49 | 0.46 | -3.26 | 1.13E-03 | 2.51E-02 |
| *Rgcc* | 47.98 | 1.48 | 0.48 | -3.10 | 1.91E-03 | 3.70E-02 |
| *Eno2* | 725.01 | 1.48 | 0.46 | -3.23 | 1.22E-03 | 2.65E-02 |
| *Zfp92* | 72.03 | 1.48 | 0.49 | -3.00 | 2.72E-03 | 4.84E-02 |
| *Ggt7* | 92.16 | 1.47 | 0.32 | -4.64 | 3.50E-06 | 2.60E-04 |
| *Prrt1* | 97.31 | 1.47 | 0.40 | -3.70 | 2.16E-04 | 6.93E-03 |
| *Pvrl3* | 161.81 | 1.47 | 0.48 | -3.04 | 2.40E-03 | 4.43E-02 |
| *Lyve1* | 37.09 | 1.46 | 0.41 | -3.53 | 4.18E-04 | 1.19E-02 |
| *Dzank1* | 90.88 | 1.46 | 0.38 | -3.87 | 1.08E-04 | 3.97E-03 |
| *Dusp1* | 2281.44 | 1.46 | 0.41 | -3.56 | 3.71E-04 | 1.07E-02 |
| *Gnao1* | 451.69 | 1.45 | 0.39 | -3.77 | 1.61E-04 | 5.51E-03 |
| *Pgbd5* | 56.21 | 1.45 | 0.46 | -3.15 | 1.66E-03 | 3.31E-02 |
| *Frmd3* | 29.57 | 1.45 | 0.46 | -3.17 | 1.54E-03 | 3.14E-02 |
| *Kcna5* | 82.33 | 1.45 | 0.31 | -4.62 | 3.78E-06 | 2.77E-04 |
| *Tceal5* | 39.01 | 1.45 | 0.44 | -3.27 | 1.09E-03 | 2.44E-02 |
| *Ptprz1* | 90.42 | 1.45 | 0.48 | -3.02 | 2.55E-03 | 4.63E-02 |
| *Apc2* | 100.82 | 1.45 | 0.48 | -3.02 | 2.52E-03 | 4.60E-02 |
| *Sall2* | 111.89 | 1.44 | 0.30 | -4.84 | 1.30E-06 | 1.20E-04 |
| *Sv2a* | 468.26 | 1.44 | 0.41 | -3.53 | 4.12E-04 | 1.17E-02 |
| *Scrt1* | 100.62 | 1.44 | 0.47 | -3.06 | 2.18E-03 | 4.12E-02 |
| *Arid3b* | 54.95 | 1.44 | 0.36 | -3.98 | 6.92E-05 | 2.75E-03 |
| *Slc24a3* | 170.42 | 1.43 | 0.38 | -3.77 | 1.66E-04 | 5.62E-03 |

| | | | | | | |
|---|---|---|---|---|---|---|
| *Rgs6* | 72.95 | 1.42 | 0.34 | -4.17 | 3.10E-05 | 1.46E-03 |
| *Slc24a2* | 42.75 | 1.42 | 0.42 | -3.35 | 8.15E-04 | 1.96E-02 |
| *Krt18* | 481.48 | 1.42 | 0.27 | -5.32 | 1.06E-07 | 1.95E-05 |
| *Clic6* | 233.02 | 1.42 | 0.23 | -6.05 | 1.46E-09 | 7.47E-07 |
| *Stxbp6* | 34.23 | 1.42 | 0.41 | -3.42 | 6.29E-04 | 1.59E-02 |
| *Mgat5b* | 20.14 | 1.41 | 0.45 | -3.13 | 1.74E-03 | 3.43E-02 |
| *Tubb4a* | 208.30 | 1.41 | 0.43 | -3.26 | 1.13E-03 | 2.51E-02 |
| *Agap2* | 97.51 | 1.41 | 0.26 | -5.53 | 3.23E-08 | 8.25E-06 |
| *Mdm1* | 168.37 | 1.41 | 0.35 | -4.02 | 5.93E-05 | 2.44E-03 |
| *Lrrc16b* | 84.59 | 1.41 | 0.47 | -3.00 | 2.67E-03 | 4.76E-02 |
| *Caskin1* | 110.83 | 1.39 | 0.35 | -4.04 | 5.41E-05 | 2.24E-03 |
| *Hid1* | 86.86 | 1.39 | 0.32 | -4.39 | 1.11E-05 | 6.32E-04 |
| *Tmem151a* | 45.06 | 1.39 | 0.37 | -3.73 | 1.93E-04 | 6.30E-03 |
| *Chd3os* | 82.58 | 1.39 | 0.44 | -3.13 | 1.73E-03 | 3.43E-02 |
| *Tceal3* | 90.24 | 1.39 | 0.40 | -3.48 | 5.05E-04 | 1.35E-02 |
| *Wfikkn2* | 71.02 | 1.38 | 0.30 | -4.58 | 4.74E-06 | 3.35E-04 |
| *Gm17024* | 35.16 | 1.38 | 0.44 | -3.17 | 1.53E-03 | 3.12E-02 |
| *Apba1* | 101.25 | 1.38 | 0.37 | -3.71 | 2.04E-04 | 6.61E-03 |
| *Rhpn1* | 75.22 | 1.38 | 0.38 | -3.67 | 2.47E-04 | 7.69E-03 |
| *Pcbp3* | 460.81 | 1.38 | 0.28 | -4.96 | 6.96E-07 | 7.99E-05 |
| *St3gal1* | 419.43 | 1.38 | 0.30 | -4.56 | 5.15E-06 | 3.59E-04 |
| *Spock1* | 208.85 | 1.37 | 0.33 | -4.17 | 3.04E-05 | 1.43E-03 |
| *P4htm* | 108.53 | 1.37 | 0.41 | -3.32 | 9.05E-04 | 2.11E-02 |
| *Grtp1* | 112.06 | 1.36 | 0.35 | -3.89 | 9.91E-05 | 3.67E-03 |
| *Klhl23* | 160.04 | 1.35 | 0.31 | -4.33 | 1.49E-05 | 8.08E-04 |
| *Sybu* | 57.23 | 1.35 | 0.32 | -4.17 | 3.03E-05 | 1.43E-03 |
| *Chd7* | 381.15 | 1.35 | 0.20 | -6.87 | 6.57E-12 | 8.78E-09 |
| *Efnb3* | 38.45 | 1.34 | 0.40 | -3.32 | 9.16E-04 | 2.13E-02 |
| *Traf3ip3* | 39.71 | 1.34 | 0.32 | -4.11 | 3.89E-05 | 1.72E-03 |
| *Ppp3cc* | 135.93 | 1.33 | 0.27 | -4.94 | 7.63E-07 | 8.44E-05 |
| *Ncald* | 129.17 | 1.33 | 0.27 | -4.90 | 9.66E-07 | 9.81E-05 |
| *Qpct* | 119.41 | 1.33 | 0.35 | -3.75 | 1.77E-04 | 5.88E-03 |
| *Camk2n2* | 58.43 | 1.33 | 0.32 | -4.16 | 3.13E-05 | 1.47E-03 |
| *Fam3c* | 417.29 | 1.32 | 0.42 | -3.16 | 1.56E-03 | 3.17E-02 |
| *Gem* | 61.42 | 1.32 | 0.33 | -3.94 | 8.20E-05 | 3.14E-03 |
| *Cttnbp2* | 58.40 | 1.32 | 0.41 | -3.20 | 1.39E-03 | 2.92E-02 |
| *Wasf3* | 172.91 | 1.31 | 0.27 | -4.79 | 1.64E-06 | 1.42E-04 |
| *Dmxl2* | 280.41 | 1.31 | 0.36 | -3.61 | 3.09E-04 | 9.27E-03 |
| *Dmd* | 140.63 | 1.31 | 0.36 | -3.63 | 2.82E-04 | 8.60E-03 |

| | | | | | | |
|---|---|---|---|---|---|---|
| *Crocc* | 573.56 | 1.31 | 0.34 | -3.83 | 1.30E-04 | 4.62E-03 |
| *Itgb8* | 268.77 | 1.30 | 0.25 | -5.18 | 2.20E-07 | 3.50E-05 |
| *Dixdc1* | 124.77 | 1.30 | 0.35 | -3.68 | 2.29E-04 | 7.27E-03 |
| *Spata1* | 113.16 | 1.29 | 0.36 | -3.56 | 3.72E-04 | 1.07E-02 |
| *Bmp2* | 87.98 | 1.29 | 0.42 | -3.09 | 2.03E-03 | 3.88E-02 |
| *Pcdhb9* | 47.02 | 1.29 | 0.41 | -3.11 | 1.90E-03 | 3.68E-02 |
| *Sh2d5* | 340.01 | 1.29 | 0.43 | -3.01 | 2.61E-03 | 4.69E-02 |
| *Pde1c* | 99.62 | 1.28 | 0.41 | -3.14 | 1.67E-03 | 3.33E-02 |
| *Tbc1d24* | 161.75 | 1.27 | 0.32 | -4.04 | 5.39E-05 | 2.24E-03 |
| *Slc2a3* | 98.60 | 1.27 | 0.29 | -4.38 | 1.17E-05 | 6.59E-04 |
| *Slc16a1* | 508.80 | 1.27 | 0.24 | -5.25 | 1.52E-07 | 2.59E-05 |
| *Celf5* | 34.12 | 1.26 | 0.37 | -3.36 | 7.66E-04 | 1.87E-02 |
| *Magi2* | 74.49 | 1.26 | 0.29 | -4.29 | 1.82E-05 | 9.36E-04 |
| *Rtn2* | 61.13 | 1.25 | 0.40 | -3.11 | 1.87E-03 | 3.65E-02 |
| *Dclk1* | 166.37 | 1.25 | 0.38 | -3.25 | 1.15E-03 | 2.54E-02 |
| *Snrpn* | 384.06 | 1.24 | 0.33 | -3.75 | 1.76E-04 | 5.88E-03 |
| *Guca1a* | 1094.43 | 1.24 | 0.34 | -3.69 | 2.20E-04 | 7.04E-03 |
| *Car12* | 138.73 | 1.23 | 0.30 | -4.04 | 5.27E-05 | 2.21E-03 |
| *Sgtb* | 84.47 | 1.22 | 0.35 | -3.51 | 4.41E-04 | 1.23E-02 |
| *Vldlr* | 440.65 | 1.22 | 0.20 | -5.95 | 2.63E-09 | 1.17E-06 |
| *Kif5a* | 156.12 | 1.22 | 0.39 | -3.14 | 1.68E-03 | 3.34E-02 |
| *Jun* | 2294.82 | 1.21 | 0.40 | -3.03 | 2.41E-03 | 4.45E-02 |
| *Serpina3h* | 79.64 | 1.21 | 0.34 | -3.52 | 4.36E-04 | 1.22E-02 |
| *Shisa4* | 90.26 | 1.21 | 0.30 | -4.11 | 4.03E-05 | 1.78E-03 |
| *Chrnb2* | 72.24 | 1.21 | 0.40 | -3.02 | 2.52E-03 | 4.60E-02 |
| *Nrep* | 305.33 | 1.21 | 0.38 | -3.17 | 1.52E-03 | 3.11E-02 |
| *Rab3a* | 347.85 | 1.20 | 0.39 | -3.07 | 2.16E-03 | 4.08E-02 |
| *Chn1* | 92.44 | 1.19 | 0.39 | -3.09 | 2.03E-03 | 3.88E-02 |
| *Kcnk3* | 58.75 | 1.19 | 0.32 | -3.77 | 1.61E-04 | 5.53E-03 |
| *Apbb1* | 266.50 | 1.19 | 0.39 | -3.09 | 2.00E-03 | 3.83E-02 |
| *Espn* | 81.01 | 1.19 | 0.30 | -3.97 | 7.30E-05 | 2.87E-03 |
| *Tmem98* | 391.90 | 1.19 | 0.36 | -3.31 | 9.38E-04 | 2.18E-02 |
| *Khdrbs3* | 234.03 | 1.18 | 0.26 | -4.61 | 4.07E-06 | 2.93E-04 |
| *Prkar1b* | 180.88 | 1.18 | 0.34 | -3.48 | 5.03E-04 | 1.34E-02 |
| *Fbxo27* | 90.03 | 1.17 | 0.30 | -3.86 | 1.12E-04 | 4.09E-03 |
| *6430571L13Rik* | 52.08 | 1.17 | 0.37 | -3.20 | 1.37E-03 | 2.89E-02 |
| *Kirrel3* | 32.58 | 1.17 | 0.36 | -3.25 | 1.15E-03 | 2.54E-02 |
| *Cp* | 455.30 | 1.17 | 0.31 | -3.78 | 1.56E-04 | 5.38E-03 |
| *Frem1* | 37.43 | 1.16 | 0.37 | -3.15 | 1.66E-03 | 3.31E-02 |

| | | | | | | |
|---|---|---|---|---|---|---|
| *Tmem8* | 72.16 | 1.16 | 0.38 | -3.04 | 2.40E-03 | 4.43E-02 |
| *Epb4.1l3* | 172.60 | 1.16 | 0.30 | -3.83 | 1.30E-04 | 4.62E-03 |
| *Slc4a8* | 84.02 | 1.16 | 0.31 | -3.71 | 2.07E-04 | 6.65E-03 |
| *Pnmal2* | 369.55 | 1.16 | 0.36 | -3.24 | 1.19E-03 | 2.59E-02 |
| *Snph* | 91.94 | 1.15 | 0.30 | -3.81 | 1.41E-04 | 4.94E-03 |
| *Ccdc136* | 199.53 | 1.15 | 0.28 | -4.14 | 3.48E-05 | 1.58E-03 |
| *Magee1* | 122.62 | 1.15 | 0.28 | -4.08 | 4.54E-05 | 1.97E-03 |
| *Peli3* | 171.11 | 1.14 | 0.30 | -3.85 | 1.19E-04 | 4.29E-03 |
| *Pip5k1b* | 161.90 | 1.14 | 0.35 | -3.30 | 9.80E-04 | 2.26E-02 |
| *Ank2* | 482.11 | 1.14 | 0.23 | -4.97 | 6.53E-07 | 7.61E-05 |
| *Ttll11* | 48.51 | 1.13 | 0.36 | -3.15 | 1.64E-03 | 3.29E-02 |
| *Elmo1* | 97.67 | 1.13 | 0.33 | -3.37 | 7.60E-04 | 1.86E-02 |
| *Rufy3* | 357.13 | 1.12 | 0.26 | -4.38 | 1.18E-05 | 6.62E-04 |
| *Sox9* | 368.93 | 1.11 | 0.36 | -3.09 | 2.03E-03 | 3.88E-02 |
| *Klhl13* | 40.47 | 1.11 | 0.36 | -3.07 | 2.13E-03 | 4.03E-02 |
| *Pclo* | 94.96 | 1.11 | 0.32 | -3.50 | 4.64E-04 | 1.27E-02 |
| *Whrn* | 258.14 | 1.10 | 0.35 | -3.18 | 1.47E-03 | 3.04E-02 |
| *A730017C20Rik* | 107.65 | 1.10 | 0.36 | -3.01 | 2.66E-03 | 4.74E-02 |
| *Wdr78* | 77.36 | 1.10 | 0.27 | -4.05 | 5.04E-05 | 2.12E-03 |
| *Cadm2* | 203.99 | 1.09 | 0.34 | -3.18 | 1.46E-03 | 3.02E-02 |
| *Mapt* | 159.97 | 1.08 | 0.36 | -3.04 | 2.34E-03 | 4.35E-02 |
| *Asrgl1* | 124.97 | 1.08 | 0.33 | -3.26 | 1.12E-03 | 2.49E-02 |
| *Nt5e* | 363.78 | 1.08 | 0.32 | -3.33 | 8.57E-04 | 2.04E-02 |
| *Mdga1* | 86.09 | 1.08 | 0.32 | -3.35 | 7.99E-04 | 1.94E-02 |
| *Pcmtd2* | 1063.20 | 1.08 | 0.26 | -4.15 | 3.31E-05 | 1.53E-03 |
| *Pla2r1* | 191.49 | 1.07 | 0.33 | -3.28 | 1.04E-03 | 2.36E-02 |
| *Tnfaip3* | 1165.02 | 1.07 | 0.33 | -3.22 | 1.26E-03 | 2.72E-02 |
| *Epb4.1l2* | 1092.79 | 1.07 | 0.20 | -5.44 | 5.29E-08 | 1.16E-05 |
| *Rundc3a* | 254.20 | 1.07 | 0.28 | -3.85 | 1.19E-04 | 4.30E-03 |
| *4931428F04Rik* | 73.32 | 1.06 | 0.31 | -3.43 | 6.00E-04 | 1.53E-02 |
| *Synpo2* | 132.80 | 1.06 | 0.29 | -3.59 | 3.29E-04 | 9.76E-03 |
| *Ankrd6* | 50.84 | 1.06 | 0.32 | -3.28 | 1.05E-03 | 2.37E-02 |
| *Ptgs1* | 1192.84 | 1.05 | 0.33 | -3.19 | 1.42E-03 | 2.96E-02 |
| *Pfkp* | 606.21 | 1.05 | 0.28 | -3.69 | 2.21E-04 | 7.05E-03 |
| *Kcnd3* | 79.57 | 1.05 | 0.35 | -3.03 | 2.43E-03 | 4.48E-02 |
| *Lurap1l* | 131.66 | 1.03 | 0.26 | -3.91 | 9.32E-05 | 3.47E-03 |
| *Atp6v0e2* | 508.31 | 1.03 | 0.22 | -4.61 | 4.05E-06 | 2.93E-04 |
| *Ahi1* | 216.26 | 1.02 | 0.23 | -4.40 | 1.11E-05 | 6.32E-04 |
| *Hk1* | 1631.02 | 1.02 | 0.18 | -5.68 | 1.33E-08 | 4.44E-06 |

| | | | | | | |
|---|---|---|---|---|---|---|
| *Mapk10* | 99.04 | 1.01 | 0.30 | -3.36 | 7.89E-04 | 1.92E-02 |
| *Mgst2* | 47.73 | -1.05 | 0.35 | 2.99 | 2.77E-03 | 4.89E-02 |
| *Fkbp5* | 5209.34 | -1.05 | 0.30 | 3.54 | 4.03E-04 | 1.15E-02 |
| *Pla2g2f* | 76.81 | -1.05 | 0.26 | 4.01 | 6.03E-05 | 2.45E-03 |
| *Hmgcs1* | 8715.87 | -1.06 | 0.34 | 3.09 | 2.02E-03 | 3.87E-02 |
| *Wdr62* | 97.81 | -1.10 | 0.25 | 4.41 | 1.02E-05 | 5.93E-04 |
| *Stag3* | 260.06 | -1.13 | 0.30 | 3.73 | 1.93E-04 | 6.30E-03 |
| *Insig1* | 2814.59 | -1.14 | 0.36 | 3.20 | 1.38E-03 | 2.92E-02 |
| *Cyp4a12a* | 467.07 | -1.38 | 0.40 | 3.47 | 5.12E-04 | 1.36E-02 |
| *Tsc22d3* | 2502.93 | -1.40 | 0.40 | 3.49 | 4.80E-04 | 1.29E-02 |
| *Gm17334* | 29.66 | -1.45 | 0.48 | 3.04 | 2.39E-03 | 4.43E-02 |
| *Fut4* | 686.68 | -1.70 | 0.49 | 3.44 | 5.92E-04 | 1.52E-02 |
| *Lzts1* | 76.10 | -1.81 | 0.57 | 3.16 | 1.56E-03 | 3.17E-02 |
| *Wnt11* | 381.85 | -1.86 | 0.54 | 3.46 | 5.46E-04 | 1.43E-02 |
| *Gm13445* | 14.15 | -2.59 | 0.68 | 3.82 | 1.36E-04 | 4.81E-03 |
| *Gm6166* | 147.65 | -4.88 | 1.52 | 3.20 | 1.37E-03 | 2.89E-02 |

RNAseq analysis was performed at day 7 of treatment

**Table S3. Enriched TF regulons in lacripep-treated corneas at day 7**

| ***Isl1*** | ***Rest*** | ***Rreb1*** |
|---|---|---|
| *Pcbp4* | *Snap25* | *Rufy3* |
| *Cabp4* | *Phyhipl* | *Nrxn2* |
| *Gpr152* | *Rtbdn* | *Car8* |
| *Cabp2* | *Chrnb2* | *Rdh12* |
| *Nr2e1* | *Cadm3* | *Kcnab2* |
| *Ccdc136* | *Fam155a* | *Lrrc16b* |
| *Bai1* | *Gabrg2* | *Pcbp4* |
| *Lrit2* | *Prmt8* | *Rims1* |
| *Tulp1* | *Cacna1b* | *Shank2* |
| *Gnb3* | *Srrm4* | *Vldlr* |
| *Isl1* | *Cacng3* | *Lingo3* |
| *Mamld1* | *Kcnh2* | *Slc6a1* |
| *Slc12a5* | *Paqr4* | *Zmat4* |
| *Bmp2* | *Cdk5r2* | *Myt1* |
| *Unc119* | *Nrsn1* | *Gem* |
| *Usp13* | *Ptprn* | *Scgn* |
| *Samd14* | *Mgat5b* | *Doc2b* |
| *Bhlhe22* | *A330050F15Rik* | *Stk32b* |
| *Cacna2d4* | *Myt1* | *Akap6* |
| *Impdh1* | *Gad1* | *Nfasc* |
| *Obscn* | *Rph3a* | *Hcn4* |
| *Rho* | *Sez6* | *Kcne2* |
| *Cspg5* | *Olfm3* | *Vat1l* |
| *Paqr4* | *Phf21b* | *Syngr3* |
| *Abca4* | *Fam163b* | *Zfp385b* |
| *Fhod3* | *Celf3* | *Wscd1* |
| *Sec14l2* | *Hcn1* | *Lrrn3* |
| *Phyhipl* | *Sybu* | *4833424O15Rik* |
| *Cngb1* | *Gprin1* | *Ppargc1b* |
| *Rtn4rl1* | *Grm5* | *Gng4* |
| *Sv2a* | *Chgb* | *Slc38a3* |
| *C2cd2l* | *Nefh* | *Fam196a* |
| *Slc8a3* | *Vgf* | *Nrcam* |
| *Fam57b* | *Pcbp3* | *Ctnnd2* |
| *Lrit1* | *Scamp5* | *Igsf21* |
| *Gm11744* | *Ctnna2* | *Cdh22* |

| | | |
|---|---|---|
| *Srrm3* | *Nfasc* | *Nlgn1* |
| *Gucy2e* | *Slc12a5* | *Neurod2* |
| *Sall3* | *C1ql2* | *Phactr3* |
| *Gucy2f* | *Cntnap2* | *Tdrkh* |
| *BC027072* | *Iglon5* | *Cacng5* |
| *Coro2b* | *Crmp1* | *Kcnh6* |
| *Faim* | *Bex2* | *Kirrel3* |
| *Esrrg* | *Chga* | *Pak3* |
| *Slc22a8* | *Gabrb3* | *Cacna1e* |
| *Msi1* | *Dner* | *Drd2* |
| *Rgr* | *St18* | *Epha5* |
| *Wdr17* | *Cacna1e* | *Impdh1* |
| *Bin1* | *Drd2* | *5430419D17Rik* |
| *Cdk5r2* | *Ank1* | *Slc8a3* |
| *Rasgrf2* | *Apbb1* | *Mcf2l* |
| *Igsf21* | *Cdh22* | *Abcg4* |
| *Pou3f1* | *Cdk5r1* | *Fam163b* |
| *Ntng2* | *Fam57b* | *St8sia1* |
| *Cdhr1* | *Acsl6* | *Sgip1* |
| *Gnat2* | *Celsr3* | *Glp2r* |
| *Chd7* | *Hpca* | *Frem1* |
| *Glp2r* | *Mmp24* | *Ppp2r2b* |
| *Cplx4* | *Wasf1* | *Gpr75* |
| *Whrn* | *Dpp6* | *Elfn2* |
| *Iglon5* | *Car10* | *Wfikkn2* |
| *6530402F18Rik* | *Gria2* | *Cdhr1* |
| *Kcnh2* | *Gabbr2* | *Zfp536* |
| *Rcvrn* | *Galnt13* | *Trpc1* |
| *Arhgef26* | *Ptprr* | *Rho* |
| *Pde6g* | *Igsf21* | *Slitrk2* |
| *Slc16a1* | *Bex1* | *Evl* |
| *Samd7* | *Scrt1* | *Whrn* |
| *Cdh23* | *Sarm1* | *Rp1l1* |
| *Gjd2* | *Syp* | *Mamld1* |
| *Kcnj14* | *Espn* | *Add2* |
| *Gabra1* | *Lrrc16b* | *Vstm2b* |
| *Otx2* | *Pclo* | *Ncald* |
| *Ggt7* | *Smpd3* | *Zswim5* |
| *Nlgn1* | *Nrxn3* | *Nrg3* |

| | | |
|---|---|---|
| *Wfikkn2* | *Rasl10b* | *Mapk10* |
| *Pcdh9* | *Nptx2* | *Fabp12* |
| *Trnp1* | *Apc2* | *Fam167a* |
| *Pcbp3* | *Slc24a2* | *Efr3b* |
| *Abcg4* | *Cadps* | *Kcnj10* |
| *Nat8l* | *Trim9* | *Tmem98* |
| *Slc38a3* | *Camk2n2* | *Nrxn3* |
| *Vsx1* | *Celf4* | *Chrnb2* |
| *Rp1l1* | *Ap3b2* | *Nrsn1* |
| *Sema7a* | *Elfn2* | *Slc16a6* |
| *Dlg4* | *Rho* | *Dusp8* |
| *Gabrr3* | *Gpr123* | *Igsf9* |
| *Neurl1a* | *Nrxn2* | *Epb4.1l2* |
| *Ano2* | *Kcnc3* | *Gnaz* |
| *Klhl23* | *Ntng2* | *Kcna5* |
| *Lingo3* | *6530402F18Rik* | *Apba1* |
| *Rgs9bp* | *Cntn4* | *Bbs9* |
| *D630045J12Rik* | *Zdhhc22* | *Rtn1* |
| *Cnr1* | *Rundc3a* | *Scamp5* |
| *Pitpnm1* | *Kcnma1* | *Elovl4* |
| *Cerkl* | *St8sia2* | *Cacng4* |
| *Dpysl5* | *Slc6a13* | *Glra2* |
| *Igsf9* | *Kcnq2* | *Rcan2* |
| *Scamp5* | *Nhlh2* | *Elmo1* |
| *Celf3* | *Myh7b* | *Srrm3* |
| *Rtbdn* | *Atp2b2* | *Tmem72* |
| *Chrna4* | *Psd* | *Dpysl5* |
| *Ptprr* | *Nr2e1* | *Hpcal4* |
| *Slc1a7* | *Ttll11* | *Ptprt* |
| *Myh7* | *Slitrk1* | *Prnp* |
| *Kcnh6* | *Nol4* | *Chst3* |
| *Sez6* | *Cxxc4* | *Elmod1* |
| *Syt1* | *Mypn* | *6530402F18Rik* |
| *Amph* | *Sh2d1a* | *Pde4dip* |
| *C1ql1* | *Kcnd3* | *Tmem108* |
| *Gpr137c* | *Lingo3* | *Sv2a* |
| *Slitrk3* | *Kcnb1* | *AI118078* |
| *Kcnj10* | *Ank2* | *Psd* |
| *Lrp2* | *Sult4a1* | *Ccdc85a* |

| | | |
|---|---|---|
| *Pvrl3* | *Rims2* | *Crb1* |
| *Cadm3* | *Mef2c* | *Kcnma1* |
| *Ntm* | *Ankrd6* | *Fscn2* |
| *Sox2* | *Rab3a* | *Phf21b* |
| *A330050F15Rik* | *Fam167a* | *Tnfrsf8* |
| *Slc4a8* | *Car2* | *Zfp804a* |
| *Ankrd6* | *Esrrg* | *Dlgap1* |
| *Pde4dip* | *Srrm3* | *Gpr162* |
| *Mdga1* | *Slc6a1* | *Ahi1* |
| *St18* | *Hcn4* | *Dnm3* |
| *Ric8b* | *Sv2b* | *Pcdh9* |
| *Cntn1* | *Celf5* | *Snap25* |
| *Elmo1* | *Kcnj10* | *Nsg1* |
| *Esrrb* | *Epb4.1l3* | *Negr1* |
| *Neurod2* | *Qpct* | *Tmem132d* |
| *Ptp4a3* | *Cacng5* | *Gpr37* |
| *Cacng3* | *Snrpn* | *Gas7* |
| *Lgi1* | *Zfp365* | *Atp2b2* |
| *St6galnac5* | *Podxl* | *Cdh8* |
| *Rax* | *Hist1h2be* | *Thsd7a* |
| *Gad1* | *Cerkl* | *Kcnd3* |
| *Sall2* | *Shank2* | *Wasf1* |
| *Scgn* | *Ncald* | *Csrp3* |
| *Pdc* | *Fstl5* | *Zfp488* |
| *Klhl13* | *Pde8b* | *Areg* |
| *Cdh8* | *Lrfn2* | *Cabp1* |
| *Slc16a6* | *Whrn* | *Phyhipl* |
| *Crb1* | *Chd7* | *Lgi1* |
| *Tmem136* | *Rp1* | *Srrm4* |
| *Zfp536* | *AI593442* | *Hspa12a* |
| *Rtn1* | *Gpr179* | *Wdr17* |
| *Hpcal4* | *Pacsin1* | *Kcnj12* |
| *Snrpn* | *Bnip3* | *Kcnq2* |
| *Cdh22* | *Fam78b* | *Syt9* |
| *Xirp1* | *Slc4a10* | *Cecr2* |
| *Car2* | *Zfp385b* | *Pak7* |
| *Fam78b* | *Kcnh6* | *Traf3ip3* |
| *Sybu* | *Pigr* | *Me1* |
| *Cecr2* | *Rtn2* | *Trpc4* |

| | | |
|---|---|---|
| *Nxnl1* | *Sez6l2* | *Nmnat2* |
| *Rbp3* | *Zfp536* | *Rlbp1* |
| *Fam196a* | *Rorb* | *St18* |
| *Ppp1r9a* | *Bai1* | *Mtmr7* |
| *Tmem27* | *Scn1a* | *Dennd5b* |
| *Tmem35* | *Pde4dip* | *Mdga1* |
| *Dusp8* | *Rp1l1* | *Fam155a* |
| *Pgbd5* | *Got1* | *Ntng2* |
| *Arid3b* | *Dpysl4* | *Dpf3* |
| *Lrrc4c* | *Kcnc1* | *Iqsec3* |
| *Zfp385b* | *Ccdc136* | *Rorb* |
| *Ttr* | *Chst3* | *Pacsin1* |
| *Ank1* | *Cacng4* | *Sybu* |
| *Gprin1* | *Chrna4* | *Atcay* |
| *Alpk2* | *Hpcal4* | *Rgs8* |
| *Slc6a13* | *Me1* | *Gabra3* |
| *Snap91* | *Abcg4* | *Tbc1d24* |
| *Grm5* | *Ptprn2* | *Scrt1* |
| *Dpp6* | *Lrrc4c* | *Ric3* |
| *Fam19a3* | *Ano2* | *Chn1* |
| *Tmem108* | *Igsf9* | *Adcy1* |
| *Gramd1b* | *Gng4* | *Dclk1* |
| *Car10* | *Nrg3* | *Acsl6* |
| *Nrxn3* | *Elfn1* | *Cacna1s* |
| *Tmem59l* | *Zfyve28* | *Cntn1* |
| *Aipl1* | *Ppp2r2b* | *Sh2d1a* |
| *Celf4* | *Pcbp4* | *Rom1* |
| *Dlgap1* | *Ly6h* | *Susd3* |
| *Dusp10* | *Glp2r* | *Mdm1* |
| *Itgb8* | *Nrcam* | *Pde6h* |
| *Prmt8* | *St8sia3* | *Cnr1* |
| *Elovl2* | *C1ql1* | *C1ql1* |
| *St3gal1* | *Slc12a5* | *Slc12a5* |
| *Dpf3* | *Ttll11* | *Ttll11* |
| *Kcnk3* | *Alpk2* | *Alpk2* |
| *Pgm2l1* | *Mpp3* | *Mpp3* |
| *Gnb5* | *Zfp365* | *Zfp365* |
| *Rgs9* | *Mmd* | *Mmd* |
| *Scrt1* | *Ntm* | *Ntm* |

| | | |
|---|---|---|
| *Zdhhc22* | *Bsn* | *Bsn* |
| *Lin7a* | *Ric8b* | *Ric8b* |
| *Eno2* | *Ank2* | *Ank2* |
| *Pde6h* | *Syt1* | *Syt1* |
| *Gm11961* | *Galnt13* | *Galnt13* |
| *Crmp1* | *Pdc* | *Pdc* |
| *Cacna1e* | *Cdk5r1* | *Cdk5r1* |
| *Gas7* | *Rph3a* | *Rph3a* |
| *Rhot1* | *Cxxc4* | *Cxxc4* |
| *Cacna1h* | *Tmem56* | *Tmem56* |
| *Phactr3* | *Eml5* | *Eml5* |
| *Sh2d1a* | *Gpm6a* | *Gpm6a* |
| *Kcnb1* | *Cntnap2* | *Cntnap2* |
| *4833424O15Rik* | *Crmp1* | *Crmp1* |
| *Cnga1* | *Rasl10b* | *Rasl10b* |
| *Igsf11* | *Nol4* | *Nol4* |
| *Nrxn2* | *Sema6a* | *Sema6a* |
| *Add2* | *Cyp1b1* | *Cyp1b1* |
| *Amigo2* | *Stxbp1* | *Stxbp1* |
| *Crocc* | *Lrfn2* | *Lrfn2* |
| *Rcan2* | *Lhx4* | *Lhx4* |
| *Fam3c* | *Csn3* | *Csn3* |
| *Insm1* | *Cadps* | *Cadps* |
| *Hkdc1* | *Myh3* | *Myh3* |
| *Stxbp1* | *Sox2* | *Sox2* |
| *Slc1a2* | *Bai1* | *Bai1* |
| *Magi2* | *Cacna2d4* | *Cacna2d4* |
| *Ccdc126* | *Cdk14* | *Cdk14* |
| *Rgs8* | *Fam161a* | *Fam161a* |
| *Rorb* | *Nhlh2* | *Nhlh2* |
| *Frmd3* | *Slc19a1* | *Slc19a1* |
| *Brsk2* | *Cttnbp2* | *Cttnbp2* |
| *Wasf1* | *Gng3* | *Gng3* |
| *Ttyh1* | *Tdrd9* | *Tdrd9* |
| *Rgs16* | *Pou3f1* | *Pou3f1* |
| *Cabp1* | *Snrpn* | *Snrpn* |
| *Pacsin1* | *Cntn4* | *Cntn4* |
| *Ush2a* | *Ildr2* | *Ildr2* |
| *Drd2* | *Pclo* | *Pclo* |

| | | |
|---|---|---|
| *Hcn4* | *Faim2* | *Faim2* |
| *Rps6kl1* | *Jun* | *Jun* |
| *Fabp12* | *Lrrc4c* | *Lrrc4c* |
| *Pde8b* | *Ryr2* | *Ryr2* |
| *Dnm3* | *Tagln3* | *Tagln3* |
| *Rims1* | *C1ql3* | *C1ql3* |
| *Nrl* | *Slc1a2* | *Slc1a2* |
| *Pigr* | *Ggt7* | *Ggt7* |
| *Pde6a* | *Cacnb2* | *Cacnb2* |
| *Ppargc1b* | *Pitpnm1* | *Pitpnm1* |
| *Cxxc4* | *Ahrr* | *Ahrr* |
| *Dusp4* | *Kcnk3* | *Kcnk3* |
| *Fam161a* | *Epha8* | *Epha8* |
| *C1ql3* | *Prox1* | *Prox1* |
| *Epha5* | *Ppm1e* | *Ppm1e* |
| *Kcnma1* | *Coro2b* | *Coro2b* |
| *Rs1* | *Pcmtd2* | *Pcmtd2* |
| *Fsd1l* | *Pde8b* | *Pde8b* |
| *Trpc1* | *Prrt3* | *Prrt3* |
| *Hpca* | *Nsg2* | *Nsg2* |
| *P4htm* | *Amph* | *Amph* |
| *Lrrc2* | *Pde10a* | *Pde10a* |
| *Mypn* | *Gjd2* | *Gjd2* |
| *Trim67* | | |
| *Snap25* | | |
| *Cntn4* | | |
| *Kif5a* | | |
| *Synpr* | | |
| *Ncald* | | |
| *Epb4.1l2* | | |
| *Sox9* | | |
| *Tmem72* | | |
| *Gucy1a3* | | |
| *Arc* | | |
| *Svop* | | |
| *Dgke* | | |
| *Gabrr2* | | |
| *Mmp12* | | |
| *Mak* | | |

| |
|---|
| *Atp1b2* |
| *Tbx3* |
| *Wscd1* |
| *Agap2* |
| *Stk39* |
| *Zfp385c* |
| *Thsd7a* |
| *Slc4a5* |
| *Rtn4rl2* |
| *Mycn* |
| *Cdk5r1* |
| *Ctnnd2* |
| *Lhx4* |
| *Cntnap2* |
| *Gngt1* |
| *Rlbp1* |
| *Nhlh2* |
| *2610034M16Rik* |
| *Dscaml1* |
| *Rpe65* |
| *Ptprt* |
| *Rpgrip1* |
| *Bai3* |
| *Shank2* |
| *Ahi1* |
| *Kcna5* |
| *Gpr37* |
| *Sgtb* |
| *Ank2* |
| *Me1* |
| *Tnfrsf8* |
| *Apba1* |
| *Cacng2* |
| *Frem1* |
| *Susd3* |
| *Eepd1* |
| *Dusp26* |
| *Fgf13* |
| *Herc3* |

| |
|---|
| *Ppm1n* |
| *Drd4* |
| *Elfn1* |
| *Kcne2* |
| *Slc1a3* |
| *Car12* |
| *Ppp3cc* |
| *Efnb3* |
| *Pde1a* |
| *Spock1* |
| *Ttn* |
| *Abcc8* |
| *Cttnbp2* |
| *Rgs6* |
| *Scg2* |
| *Syt9* |
| *Rph3a* |
| *Aqp4* |
| *4931428F04Rik* |
| *Nrsn1* |
| *Slitrk1* |
| *Lrrn3* |
| *Pak3* |
| *Mdm1* |
| *Prox1* |
| *Kcnd3* |
| *Slc24a2* |
| *Stxbp6* |
| *Tceal5* |
| *Phf21b* |
| *Slc6a1* |
| *Srrm4* |
| *Guca1b* |
| *Gem* |
| *Gabbr2* |
| *Tmem130* |
| *Camk2n2* |
| *5430419D17Rik* |
| *Podxl* |

| |
|---|
| *Dclk1* |
| *Nmnat2* |
| *Spata1* |
| *Guca1a* |
| *Lrfn2* |
| *Fam163b* |
| *Kif1a* |
| *Nfasc* |
| *Kirrel3* |
| *Prdm8* |
| *Pde1b* |
| *Gpr75* |
| *Olfm3* |
| *Nrcam* |
| *Tbc1d24* |
| *Pde6c* |
| *Pcmtd2* |
| *Cyp1b1* |
| *Ryr2* |
| *Chn1* |
| *Zmat4* |
| *Epb4.1l3* |
| *Tox2* |
| *Ppef2* |
| *Prnp* |
| *Chst3* |
| *Chrnb2* |
| *Elmod1* |
| *Sgip1* |
| *Clic6* |
| *Grik1* |
| *Pcdh10* |
| *Espn* |
| *Fam184b* |
| *Hist2h4* |
| *Slc17a7* |
| *Efr3b* |
| *Plch1* |
| *Myrip* |

| |
|---|
| *Ppm1e* |
| *Slc4a10* |
| *Crx* |
| *Lhfpl4* |
| *Krt18* |
| *Kcnv2* |
| *Kif21b* |

**Table S4. Unique gene targets within TF regulons enriched in lacripep-treated corneas at day 7**

| ***Isl1*** | ***Rreb1*** | ***Rest*** |
|---|---|---|
| *Klhl13* | *Dennd5b* | *Sarm1* |
| *Cabp2* | *Gng3* | *Chga* |
| *Sall2* | *Bbs9* | *Gpr179* |
| *Stxbp6* | *Hspa12a* | *Smpd3* |
| *Snap91* | *Rom1* | *Sv2b* |
| *Pgm2l1* | *Iqsec3* | *Nptx2* |
| *Herc3* | *Nsg1* | *Kcnc1* |
| *Pde6a* | *Pde10a* | *Sult4a1* |
| *Gm11744* | *Mpp3* | *Bnip3* |
| *Rgs9bp* | *Doc2b* | *Rest* |
| *Elovl2* | *Tdrd9* | *Gpr123* |
| *Ush2a* | *Tagln3* | *Nefh* |
| *Ccdc126* | *Slitrk2* | *Rab3a* |
| *Myrip* | *Pak7* | *Celf5* |
| *Mycn* | *Gpr162* | *Hist1h2be* |
| *Samd7* | *Csn3* | *Celsr3* |
| *Slc1a3* | *Sema6a* | *Kcnc3* |
| *Atp1b2* | *Zfp488* | *Apc2* |
| *Spata1* | *Bsn* | *Vgf* |
| *Pcdh10* | *Csrp3* | *Ctnna2* |
| *Kif21b* | *Ric3* | *Trim9* |
| *Lrp2* | *Myh3* | *Qpct* |
| *Zfp385c* | *Tmem98* | *Hcn1* |
| *Slc4a5* | *Cacna1s* | *Fstl5* |
| *Pde6g* | *Vldlr* | *Rundc3a* |
| *Slc22a8* | *Epha8* | *Mgat5b* |
| *Dusp26* | *Trpc4* | *AI593442* |
| *Ttyh1* | *AI118078* | *Scn1a* |
| *Pde1a* | *Syngr3* | *St8sia3* |
| *Gm11961* | *Nsg2* | *Cacna1b* |
| *C2cd2l* | *Ildr2* | *Apbb1* |
| *Gabra1* | *Gabra3* | *Gria2* |

| | | |
|---|---|---|
| *Kcnj14* | *Mcf2l* | *Got1* |
| *Eepd1* | *Rdh12* | *Dner* |
| *Slc1a7* | *Vat1l* | *Chgb* |
| *4931428F04Rik* | *Akap6* | *Rp1* |
| *Hist2h4* | *Cdk14* | *Ptprn2* |
| *Tbx3* | *Kcnj12* | *Ly6h* |
| *Itgb8* | *Eml5* | *Ptprn* |
| *Sall3* | *Rreb1* | *Gabrg2* |
| *Arid3b* | *Gnaz* | *Mef2c* |
| *Pgbd5* | *Cacnb2* | *Dpysl4* |
| *Myh7* | *Negr1* | *Zfyve28* |
| *Nxnl1* | *Rufy3* | *Mmp24* |
| *Lrit1* | *Atcay* | *St8sia2* |
| *Igsf11* | *Faim2* | *Gabrb3* |
| *Rasgrf2* | *Stk32b* | *Bex2* |
| *Sec14l2* | *Zswim5* | *Myh7b* |
| *Rgs9* | *Prrt3* | *Ap3b2* |
| *Svop* | *Fscn2* | *C1ql2* |
| *Slc17a7* | *Tdrkh* | *Bex1* |
| *Tmem35* | *Mapk10* | *Sez6l2* |
| *Brsk2* | *Kcnab2* | *Syp* |
| *Rps6kl1* | *Zfp804a* | *Rtn2* |
| *Hkdc1* | *Slc19a1* | *Rims2* |
| *Cdh23* | *Mtmr7* | |
| *Aqp4* | *St8sia1* | |
| *Arc* | *Glra2* | |
| *Vsx1* | *Ccdc85a* | |
| *Ppp1r9a* | *Jun* | |
| *Frmd3* | *Car8* | |
| *Gucy2e* | *Elovl4* | |
| *Pde1b* | *Mmd* | |
| *Gabrr2* | *Gpm6a* | |
| *Rtn4rl1* | *Vstm2b* | |
| *Samd14* | *Tmem132d* | |
| *Neurl1a* | *Ahrr* | |
| *Tceal5* | *Adcy1* | |

| | |
|---|---|
| *Sox9* | *Tmem56* |
| *Cspg5* | *Evl* |
| *Rtn4rl2* | *Areg* |
| *Abca4* | *Traf3ip3* |
| *Usp13* | |
| *Rgr* | |
| *Bmp2* | |
| *Msi1* | |
| *Aipl1* | |
| *Gabrr3* | |
| *Trim67* | |
| *Car12* | |
| *Otx2* | |
| *Bin1* | |
| *Eno2* | |
| *Abcc8* | |
| *Amigo2* | |
| *Dusp10* | |
| *Grik1* | |
| *Synpr* | |
| *Rhot1* | |
| *Esrrb* | |
| *Dgke* | |
| *Tulp1* | |
| *Crx* | |
| *Efnb3* | |
| *Rpe65* | |
| *Agap2* | |
| *Spock1* | |
| *Lrit2* | |
| *Rgs6* | |
| *Tmem59l* | |
| *Gnat2* | |
| *Cnga1* | |
| *Tmem27* | |
| *Tmem136* | |
| *Kcnv2* | |

| |
|---|
| *Slc4a8* |
| *Fam184b* |
| *Rcvrn* |
| *Pde6c* |
| *P4htm* |
| *Lhfpl4* |
| *Gngt1* |
| *Trnp1* |
| *Cacng2* |
| *Bai3* |
| *Gnb3* |
| *Gucy1a3* |
| *Unc119* |
| *Faim* |
| *Xirp1* |
| *Tox2* |
| *Rbp3* |
| *Arhgef26* |
| *Dlg4* |
| *Ptp4a3* |
| *Crocc* |
| *Gucy2f* |
| *2610034M16Rik* |
| *D630045J12Rik* |
| *Ppm1n* |
| *Gpr152* |
| *St6galnac5* |
| *Isl1* |
| *Stk39* |
| *Gramd1b* |
| *Cngb1* |
| *Magi2* |
| *Gpr137c* |
| *Fam3c* |
| *Rpgrip1* |

| |
|---|
| *Lin7a* |
| *Drd4* |
| *Slitrk3* |
| *Fam19a3* |
| *Ttr* |
| *Clic6* |
| *Mak* |
| *Mmp12* |
| *Ppef2* |
| *Kif5a* |
| *Tmem130* |
| *Cabp4* |
| *Scg2* |
| *BC027072* |
| *Fgf13* |
| *Cplx4* |
| *Rs1* |
| *Kif1a* |
| *Rax* |
| *Fhod3* |
| *Guca1b* |
| *Cacna1h* |
| *Slc16a1* |
| *Ppp3cc* |
| *Ttn* |
| *Nat8l* |
| *Prdm8* |
| *Fsd1l* |
| *Pvrl3* |
| *Klhl23* |
| *Krt18* |
| *Gnb5* |
| *Lrrc2* |
| *Plch1* |
| *Sema7a* |
| *Bhlhe22* |
| *St3gal1* |

| |
|---|
| *Sgtb* |
| *Dusp4* |
| *Nrl* |
| *Rgs16* |
| *Insm1* |
| *Dscaml1* |
| *Guca1a* |
| *Obscn* |

**Data S1. (separate file)**
Readcounts at 7 days

**Data S2. (separate file)**
Gene Ontology terms for lacripep- versus PBS-treated *Aire* KO corneas
